# Decoding causal genes and programs from regulatory variants in aortic valve disease

**DOI:** 10.64898/2026.05.26.727981

**Authors:** Mewen Briend, Anne Rufiange, Valentine Duclos, Samuel Mathieu, Thibaut Kanmacher, Dominique K. Boudreau, Nathalie Gaudreault, Victoria Saavedra Armero, François Dagenais, Christian Couture, Philippe Joubert, Sébastien Thériault, Yohan Bossé, Patrick Mathieu

## Abstract

Aortic valve disease is common, yet its regulatory mechanisms remain poorly understood. We performed multi-omic profiling of human aortic valve interstitial cells (HAVICs), identifying 11,891 allele-specific chromatin accessibility QTLs (as-caQTLs), 48% novel to this cell type. These variants were enriched in active enhancers, disrupted transcription factor (TF) motifs, particularly AP-1, TEAD and GATA families, and were validated by allele-specific TF binding assays. A fine-tuned deep DNA sequence model prioritized common and rare variants at risk loci predicted to impact chromatin accessibility. Single-cell CRISPRi perturbation of 247 variants identified cis-target genes at 55 as-caQTL elements, including loci without eQTLs. We demonstrate that common regulatory variants controlling elastin and fibrillin impact the development of the aortic valve apparatus. We provide genetic evidence and a mechanistic framework for the contribution of a reduced aortic root size to CAVD risk. Perturbations identified core cell programs led by upstream regulators *AHNAK*, *PDIA6*, and *RNFT1* converging on extracellular matrix production and iron transport.

## Introduction

The aortic valve (AV) apparatus, comprising the valve leaflets and associated anatomical structures such as the aortic root, annulus, and sino-tubular junction (STJ), ensures unidirectional blood flow from the heart to the systemic circulation^1^. Calcific aortic valve disease (CAVD), the most common heart valve disorder, is characterized by progressive thickening and mineralization of the valve leaflets^2,3^. Given the structural and functional complexity of the AV apparatus, pathological processes that manifest in the leaflets interact with other components of the AV apparatus. For example, STJ-to-annulus diameter ratio and geometry of the aortic root shape the hemodynamic environment experienced by the AV impacting leaflets mechanics and stress^4,5^. Despite these considerations, the genetic programs and cellular mechanisms governing the coordinated development of the AV apparatus, and their implications for CAVD, remain largely unknown.

Recent genome-wide association studies (GWAS) have identified 261 risk loci for CAVD^6^ and more than 130 loci associated with aortic root size and peak flow velocity at the STJ^7,8^. As with many complex traits, most of these variants lie in noncoding regions of the genome^9^. In parallel, whole-genome sequencing (WGS) in the UK Biobank has uncovered rare noncoding variants linked to disorders of the AV apparatus^10^. Together, these findings highlight a central challenge of polygenic traits-disorders: identifying the causal variants, the genes they regulate, and the core cellular programs through which they influence the AV. Here we address these questions through multi-omic profiling and functional genomics of human aortic valve interstitial cells (HAVICs). We identify allele-specific chromatin accessibility quantitative trait loci (as-caQTLs), which capture regulatory variants that directly alter chromatin accessibility. These variants were prioritized for high-content single-cell CRISPR interference (CRISPRi) perturbation screens to link disease-associated variants to their target genes and downstream cell programs. In parallel, we use deep learning model to predict regulatory grammar and variant effects on chromatin accessibility in HAVICs. Integrating these approaches with functional validation, including production of extra-cellular matrix (ECM), reporter assays and transcription factor (TF) binding, reveals variant gene connections. These analyses identify key regulators of aortic root development, and core HAVIC gene programs underlying CAVD.

## Results

### as-caQTLs map the regulatory landscape of HAVICs

We isolated HAVICs from a total of 40 biological replicates and performed comprehensive multi-omic profiling totaling ∼ 30 billion sequenced reads **(Figure 1A, Supplementary Table 1)**. RNA-seq expression confirmed canonical fibroblast (*FAP*, *TNC*, *S100A4*), smooth muscle (*ACTA2*, *TAGLN*, *CALD1*), and mesenchymal stem cell markers (*CD44*, *THY1*, *PDGFRA*), consistent with HAVIC identity **(Figure 1B)**. We performed ATAC-seq on 16 biological replicates and, after confirming high inter-sample correlation, defined a consensus set of 326,594 chromatin-accessible regions **(Supplementary Figure 1)**. Leveraging intrasample allelic imbalance at heterozygous single nucleotide variants (SNVs) (**Supplementary Figure 2A, methods)**^11–13^, which mitigates technical and biological noise, we identified 11,891 allele-specific chromatin accessibility QTLs (as-caQTLs, FDR < 0.1), representing 3.1% of 382,722 testable heterozygous SNVs (**Figure 1C, Supplementary Figure 2B, Supplementary Table 2**). Increasing ATAC-seq coverage improved detection of as-caQTLs with smaller effect sizes (**Figure 1D**, **Supplementary Figure 2C**). Most ATAC peaks contained a single as-caQTL, whereas a subset harbored two or more (**Figure 1E**). Notably, 48% of these as-caQTLs had not been reported in existing chromatin accessibility QTL resources across hundreds of cell types (**Figure 1F**), underscoring the cell-type specificity of this regulatory layer^11,14,15^.

**Figure 1:**
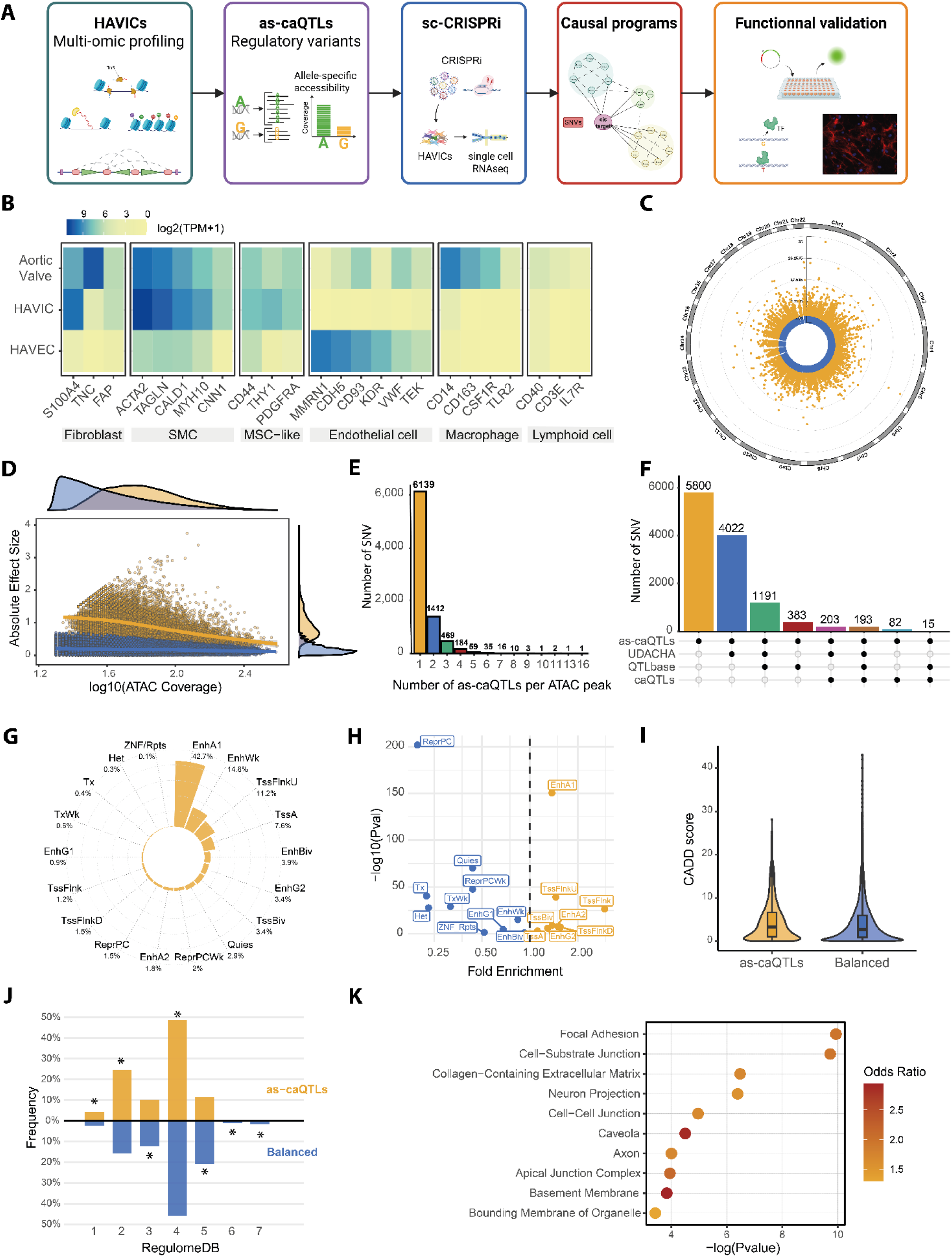
Allele-specific chromatin accessibility QTLs map the regulatory landscape of human aortic valve interstitial cells. **A**, Overview of the multi-omic profiling strategy in HAVICs (n = 40 biological replicates, ∼30 billion sequenced reads). **B**, RNA-seq expression of canonical fibroblast (*FAP, TNC, S100A4*), smooth muscle (*ACTA2, TAGLN, CALD1, MYH10, CNN1*), mesenchymal stem cell (*CD44, THY1, PDGFRA*), endothelial (*MMRN1, CDH5, CD93, KDR, VWF, TEK*), macrophage (*CD14, CD163, CSF1R, TLR2*) and lymphoid markers (*CD40, CD3E, IL7R*) in bulk aortic valve tissue, isolated HAVICs and isolated HAVECs. **C**, Manhattan plot of as-caQTLs identified from allelic imbalance in ATAC-seq at heterozygous SNVs (n = 11,891 as-caQTLs at FDR < 0.1, out of 382,722 testable SNVs). **D**, Detection of as-caQTLs effect size as a function of increasing ATAC-seq sequencing depth, showing improved sensitivity for variants with smaller effect sizes. **E**, Distribution of the number of as-caQTLs per ATAC-seq peak. **F**, Overlap of as-caQTLs with previously reported chromatin accessibility QTLs across cell types; 48% are novel to HAVICs. **G**, Fraction of as-caQTLs across ChromHMM chromatin states derived from ChIP-seq of six histone marks in HAVICs. **H**, Enrichment of as-caQTLs relative to balanced heterozygous SNVs across chromatin states (Fisher’s exact test). **I**, CADD scores for as-caQTLs compared to balanced SNVs (P = 7.78E-37, Wilcoxon rank-sum test). **J**, RegulomeDB scores for as-caQTLs versus balanced SNVs; asterisks indicate significant enrichment (Fisher’s exact test). **K**, Gene Ontology enrichment analysis for genes closest to as-caQTLs (hypergeometric test).

An 18-state ChromHMM^16^ model from ChIP-seq of six histone marks in HAVICs (**Supplementary Figure 2D)** showed that active enhancers harbored the greatest fraction of as-caQTLs (42.7%) **(Figure 1G)** and were significantly enriched relative to balanced heterozygous SNVs (OR: 1.37, P = 3.7E-151, Fisher’s exact test), as were flanking TSS regions (OR: 1.45, P = 6.6E-40, Fisher’s exact test) (**Figure 1H**), highlighting the potential of as-caQTLs to regulate gene expression from both distal and proximal elements. Consistently, as-caQTLs had higher combined annotation dependent depletion (CADD) scores (P = 7.78E-37, Wilcoxon rank-sum test) **(Figure 1I)** and lower RegulomeDB scores than balanced SNVs, reflecting greater predicted regulatory potential (**Figure 1J**).

Genes closest to as-caQTLs were enriched for focal adhesion (OR: 1.98, P = 1.15E-10, hypergeometric test), collagen-containing ECM (OR: 1.73, P = 3.32E-7, hypergeometric test) (**Figure 1K**) and fetal cardiac fibroblast-specific gene programs (OR: 1.91, P = 2.19E-05, hypergeometric test)^17^. Importantly, as-caQTL-mapped genes were overrepresented among known AV disorder genes from DISGENET^18^ (OR: 1.20, P = 0.009, hypergeometric test) and monogenic thoracic aorta and aortic root disorders catalogued in PanelApp^19^ and GENTAC^20^ (OR: 1.93, P = 0.004, hypergeometric test), reinforcing pathway convergence between common polygenic and rare monogenic disease mechanisms.

### as-caQTLs reside in conserved regions and alter transcription factor binding

Flanking regions of as-caQTLs (±50 bp) showed higher evolutionary conservation than balanced SNVs, with elevated PhastCons scores (P = 3.9E-11, Wilcoxon rank-sum test) (**Figure 2A**) and enrichment in high-constraint regions as defined by the Genomic Evolutionary Rate Profiling (GERP) score^21^ (GERP ≥ 4; OR: 1.32, P = 1.96E-05, Fisher’s exact test). TOBIAS^22^ algorithm in HAVIC ATAC-seq data showed enriched TF footprints at and near as-caQTLs (P < 1E-308, Kolmogorov–Smirnov test) (**Figure 2B**).

**Figure 2:**
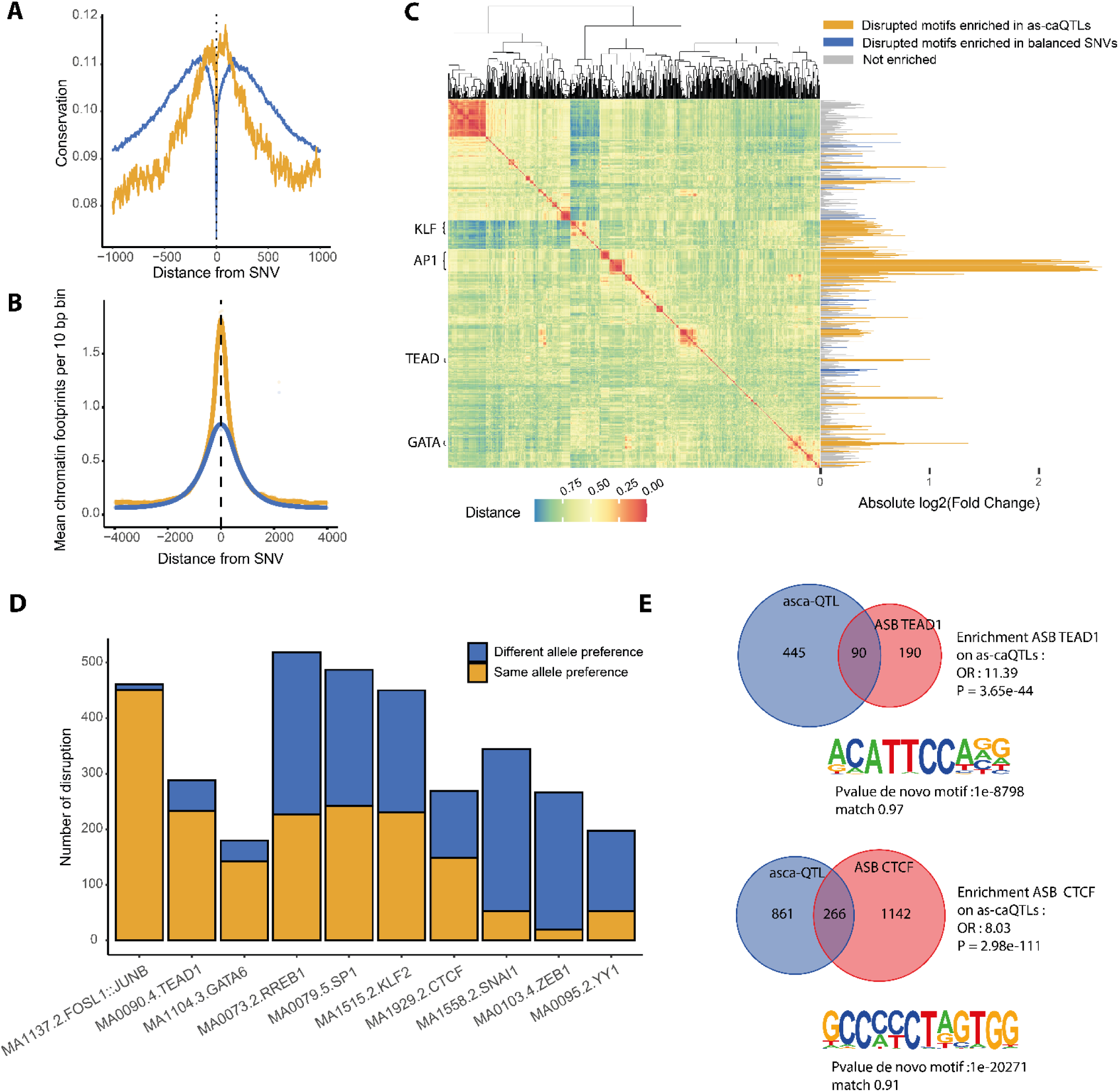
as-caQTLs reside in evolutionarily conserved regions and alter transcription factor binding. **A**, Tag density plot of PhastCons conservation scores in flanking regions of as-caQTLs compared to balanced SNVs (PhastCons score as-ca vs balanced ±50 bp, P = 3.9E-11, Wilcoxon rank-sum test). **B**, Tag density plot of TF footprint density (TOBIAS) around as-caQTLs versus balanced SNVs in HAVIC ATAC-seq data (P < 1E-308, Kolmogorov–Smirnov test). **C**, Enrichment of disrupted TF motifs (JASPAR) at as-caQTLs. Left: heatmap of motif distance with hierarchical clustering. Right: log₂(fold change) of motif disruption versus balanced SNVs (Fisher’s exact test). **D**, Frequency of concordant versus discordant allele preference between chromatin accessibility and predicted TF binding, distinguishing activator from repressor TFs. **E**, Validation of allele-specific TF binding in HAVICs and *denovo* motif of ChIPseq peaks (HOMER). Top: enrichment of as-caQTLs for allele-specific TEAD1 binding (Fisher’s exact test). Bottom: enrichment of as-caQTLs for allele-specific CTCF binding (Fisher’s exact test).

Using JASPAR, we found that as-caQTLs disrupt TF motifs^13^ with strongest enrichments for FOSL1 (OR: 5.96, P = 8.70E-175, Fisher’s exact test) and related AP-1 family members, alongside KLF, TEAD and GATA family TFs with established roles in heart development (**Figure 2C, Supplementary Table 3**)^23–26^. The relationship between allelic preference for chromatin accessibility and TF motif scores revealed distinct patterns: for activator TFs (AP-1, TEAD, GATA) the accessibility-increasing allele was typically the preferred motif allele, whereas for repressor TFs (SNAI1, ZEB1, YY1) allelic preferences were often discordant (**Figure 2D**), consistent with a model in which activators promote and repressors constrain chromatin accessibility^27,28^. We validated these predictions using the database of allelic dosage-corrected allele-specific human TF binding sites (ADASTRA)^13^, finding a 4.10-fold enrichment of as-caQTLs for allele-specific TF binding (TF-ASB) relative to balanced SNVs (P < 1E-308, Fisher’s exact test). ChIP-seq for TEAD1 (n=3) and CTCF (n=2) in HAVICs identified 2,369 TF-ASB events, as-caQTLs showed an 8.24-fold enrichment for TF-ASB (P = 4.0E-149, Fisher’s exact test) with 94% concordance in effect direction (**Figure 2E, Supplementary Table 4-5**), providing orthogonal validation that allelic chromatin imbalance captures molecular events directly linked to TF recruitment.

### as-caQTLs capture regulatory elements controlling transcriptional activity

Activity-by-contact (ABC)^29^ using in situ HiC in HAVICs predicted enhancer-gene connections and revealed enrichment of as-caQTLs within these regulatory elements (REs) (OR: 1.64, P = 1.8E-129, Fisher’s exact test). Consistently, as-caQTLs were also enriched in H3K27ac-HiChIP loops linking distal REs to gene promoters (OR: 1.18, P = 1.67E-56, Fisher’s exact test), supporting a role for these variants in distal enhancer regulation of target genes. Active REs frequently produce short RNAs^30^. To determine whether as-caQTLs mark sites of transcription initiation, we performed 5′-capped small RNA sequencing (5′csRNA-seq)^31^ in HAVICs from seven donors, identifying 38,329 transcription start sites (TSSs), including 24,099 proximal (within 500 bp of annotated genes) and 14,230 distal TSSs. The accuracy of TSS identification was confirmed by the canonical initiator YR dinucleotide and the TATA box motifs peaking 20–30 nucleotides upstream (**Supplementary Figure 3A-B**). The spatial organization of TF motifs around TSSs was consistent with their regulatory roles: activator motifs (AP-1, KLF, NFY) peaked ∼50 bp upstream, while the repressor YY1 peaked ∼10 bp downstream of TSSs (**Supplementary Figure 3C**), supporting a model where spatial relationships between TFs and transcription initiation define activator versus repressor function^32^. as-caQTLs were significantly enriched near active TSSs compared to random expectation and balanced SNVs (P < 1E-308 and P = 2.7E-10, Kolmogorov–Smirnov test) (**Supplementary Figure 3D**), with their peak density upstream of TSSs mirroring TF spatial distribution (**Figure 3A, Supplementary Figure 3E**). Nascent transcript levels were significantly elevated around as-caQTLs compared to balanced SNVs (P < 1E-308, Wilcoxon rank-sum test) (**Figure 3B**).

**Figure 3:**
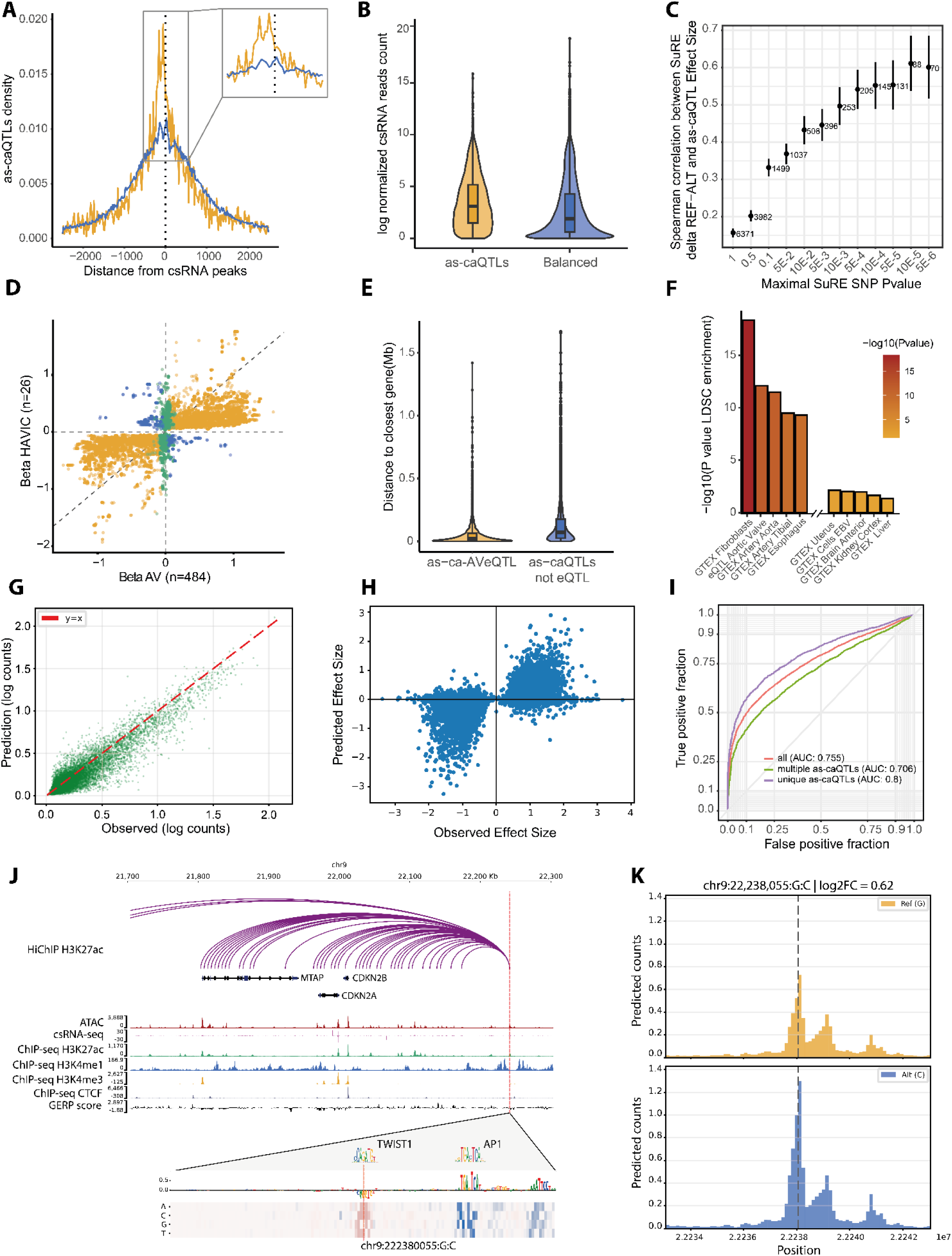
as-caQTLs mark transcriptionally active regulatory elements and associate with gene expression. **A**, Spatial distribution of as-caQTL (yellow) density relative to active transcription start sites (TSSs) identified by 5′csRNA-seq, compared to balanced SNVs (blue). **B**, Nascent transcript levels (5′csRNA-seq) within ±250 bp of as-caQTLs compared to balanced SNVs (P < 1E-308, Wilcoxon rank-sum test). **C**, Spearman correlation between SuRE reporter activity and as-caQTL allelic effects across categories of significant SuRE SNVs. **D**, Comparison of HAVICs eQTLs (n = 26) with bulk aortic valve tissue (n = 484), identifying 2,567 HAVIC-specific QTL pairs (green) and 1,040 pairs with opposite effects (blue) (Z-test). **E**, Distance to TSS for as-caQTLs shared with AV eQTLs versus unshared as-caQTLs (P < 1E-308, Wilcoxon rank-sum test). **F**, Stratified LD score regression showing heritability enrichment of as-caQTLs across GTEx tissues and aortic valve eQTLs (full results in Supplementary Figure 4F). **G**, Performance of the fine-tuned Enformer model on held-out HAVIC ATAC-seq test data (Pearson’s r = 0.93). **H**, Correlation between Enformer-predicted allelic fold-changes and experimentally observed as-caQTL effects (n = 11,891, Spearman’s ρ = 0.57). **I**, Classification performance (AUC) for prediction of as-caQTL allelic direction. **J**, Rare noncoding variant at the 9p21. Tracks show, from top to bottom: H3K27ac HiChIP loops, gene annotations, ATAC-seq, 5′csRNA-seq, H3K27ac, H3K4me1, H3K4me3 and CTCF ChIP-seq, and GERP conservation score. In silico saturation mutagenesis predicts disruption of a TWIST1 binding motif by the rare allele. **K**, Enformer-predicted chromatin accessibility at chr9:22238055:G:C for both alleles, showing that the C allele increases chromatin accessibility (predicted log₂FC = 0.62).

We further validated the transcriptional impact of as-caQTLs using the Survey of Regulatory Elements (SuRE)^33^, a massively parallel reporter assay evaluating 5.9 million SNVs. as-caQTLs were 2.2-fold enriched in significant SuRE variants (P = 1.89E-52, Fisher’s exact test), with allelic effects on chromatin accessibility correlating with reporter expression (Spearman’s ρ = 0.61 for top-significant hits, P < 1E-308) (**Figure 3C, Supplementary Figure 4A**).

### as-caQTLs and gene expression

RNA-seq from 26 genotyped HAVICs identified 34,115 significant expression QTLs (eQTLs) (FDR < 10%) for 1,306 expressed genes (eGenes). The strongest association was rs7629767 with *ULK4* expression (P_eQTL_ = 6.72E-15), a variant associated with pulse pressure (P_GWAS_ = 1.37E-95) and ascending aorta diameter (P_GWAS_ = 1.02E-17). Comparison with our published eQTL dataset from 484 bulk aortic valves (AVs)^34^ revealed 2,567 HAVIC-specific QTL pairs and 1,040 pairs with opposite effects for 60 eGenes (**Figure 3D, Supplementary Table 6**), highlighting cell-type-specific regulation masked in whole-tissue measurements. For instance, the blood pressure risk allele C-rs2493132 (P_GWAS_ = 1.18E-12) is associated with higher *AGT* expression in HAVICs (β = 0.74, P_eQTL_ = 8.63E-05) but lower expression in bulk AV (β = −0.10, P_eQTL_ = 2.23E-04), reflecting opposing regulatory effects across cell populations. Among HAVIC eQTLs, 200 as-caQTLs tagged 124 eGenes. Leveraging the larger AV dataset, eQTLs were enriched in the vicinity of as-caQTLs (P < 1E-308, Kolmogorov-Smirnov test) (**Supplementary Figure 4B).** In total, 6,504 as-caQTLs (55%) were shared with AV eQTLs (as-ca-AVeQTLs), linking 7,116 eGenes with 58.8% directional concordance. Shared as-ca-AVeQTLs were closer to TSSs (54.7 kb vs. 139.6 kb, P < 1E-308, Wilcoxon rank-sum test) (**Figure 3E**), showed higher minor allele frequency (MAF) and were enriched in ABC-predicted REs (OR: 4.74, P < 1E-308, Fisher’s exact test) (**Supplementary Figure 4C**). The remaining 45% of as-caQTLs with no eQTL were enriched in distal enhancers **(Supplementary Figure 4D)**, and had larger chromatin accessibility effect sizes (0.95 ± 0.41 vs. 0.86 ± 0.37, P = 9.99E-32, Wilcoxon rank-sum test) **(Supplementary Figure 4E)**, indicating that as-caQTLs capture a regulatory layer extending beyond expression-based approaches^35,36^. Stratified LD score regression^37^ showed heritability enrichment for AV and fibroblast (GTEx) eQTLs (8.24 and 10.17) with a stronger enrichment for the top 1,000 as-caQTLs **(Figure 3F, Supplementary Figure 4F-G, Supplementary Table 7)**. as-caQTLs were also enriched for the heritability of CAVD (OR: 8.24, P_S-LDSC_ = 7.8E-4). These findings, together with the observation that 45% of as-caQTLs escape eQTL detection, motivated us to develop a sequence-based approach to predict allele-specific chromatin accessibility directly from DNA sequence and prioritize causal variants at GWAS loci.

### Deep neural network models predict causal chromatin accessibility variants

We fine-tuned the transformer-based Enformer^38^ model on HAVIC ATAC-seq data, achieving excellent predictive accuracy on held-out test data (Pearson’s r = 0.93) **(Figure 3G)**. Critically, the model captured allele-specific effects: predicted allelic fold-changes correlated with experimentally observed as-caQTL effects (Spearman’s ρ = 0.57) **(Figure 3H)**, with stronger concordance when the causal variant was unambiguously assigned as the sole SNV within its ATAC-seq peak (Spearman’s ρ = 0.67) **(Supplementary Figure 5A)**. Classification of as-caQTLs versus balanced SNVs yielded an AUC of 0.68, rising to 0.71 for unique variants per peak **(Supplementary Figure 5B)**, and directional prediction reached an AUC of 0.80 for the highest-confidence subsets **(Figure 3I)**.

We applied this model to prioritize causal variants at GWAS loci. Among 24,203 genome-wide significant CAVD SNVs^6^, 3,028 overlapped with ATAC-seq peaks, of which 194 were predicted to alter chromatin accessibility at high specificity (0.95), including 13 lead SNVs **(Supplementary Table 8)**. These variants recapitulated hallmark properties of experimental as-caQTLs, including elevated CADD scores (P =1.6E-09, Wilcoxon rank-sum test) **(Supplementary Figure 5C)**, disruption of AP-1 family motifs **(Supplementary Table 9)**, enrichment for TF-ASB (ADASTRA fold change = 3.8, P = 4.46E-12, Fisher’s exact test) and enrichment in fine-mapped credible sets from the probabilistic identification of causal SNVs (PICS) (PICS 99%; fold change = 1.42, P = 0.01, Fisher’s exact test). Among these, the T allele of rs178004, a lead CAVD variant (P_GWAS_ = 1.08E-18), was predicted to increase chromatin accessibility and is connected by H3K27ac-HiChIP to the promoter of *GATA6*, a key cardiac developmental TF, located ∼335 kb downstream^39^. In silico saturation mutagenesis revealed a composite RE consisting of a canonical AP-1 motif (TGAGTCA) followed by a 7-bp spacer and a second motif (AGTCAT) harboring rs178004. Attention weight analysis showed that the risk C allele disrupts the second motif and attenuates the contribution of the first, suggesting a cooperative architecture in which both sites are required for full enhancer activity **(Supplementary Figure 5D)**. We then extended the model to rare noncoding variation from UK Biobank WGS^10^, interrogating 864 rare variants (MAF < 0.01, P_GWAS_ < 1E-05) associated with CAVD and 955 with aortic aneurysm and dissection. Of these, 52 and 31 overlapped ATAC-seq peaks, from which the model flagged 2 and 5 causal candidate variants impacting chromatin accessibility, respectively **(Supplementary Table 10)**. The strongest signal was a genome-wide significant variant for CAVD at the 9p21 risk locus (chr9:22238055:G:C, P_GWAS_ = 9.5E-10, MAF = 2.5E-04), located in a highly conserved region (GERP score = 4.9). In silico saturation mutagenesis at this position predicted disruption of a TWIST1 binding motif by the rare allele, leading to increased chromatin accessibility, a prediction supported by overlapping TWIST1 ChIP-seq peaks from ChIP-Atlas. TWIST1 is a master regulator of heart valve development, and its role in CAVD has been supported by Mendelian randomization^34,40^. In HAVICs, H3K27ac-HiChIP identified chromatin loops connecting this rare variant region to the promoters of *MTAP*, *CDKN2A*, and *CDKN2B* (**Figure 3J-K**).

### Single-cell CRISPRi perturbation resolve target genes and disease mechanisms at CAVD risk loci

To empirically link regulatory variants to target genes, we implemented single-cell RNA-seq CRISPR interference (CRISPRi) perturbation in primary HAVICs from two donors stably expressing KRAB-dCas9-MeCP2^41^ **(Figure 4A)**. We performed two complementary screens. In a first screen (library 1), we targeted 38 CAVD GWAS variants prioritized by functional annotations, to establish the CRISPRi perturbation framework in HAVICs and validate previously nominated candidate genes (mean of 205 cells per gRNA; 8,075 cells after QC; **Supplementary Figure 6, Supplementary Table 11**). In a second expanded screen (library 2), we targeted 220 as-caQTLs across 190 REs associated with GWAS loci (P_GWAS_ < 5E-08), suggestive associations (5E-08 ≤ P_GWAS_ < 1E-05), or fine-mapped by PICS^9^ for AV traits (mean of 575 cells per gRNA; 24,082 cells after QC; **Supplementary Figure 7, Supplementary Table 12**). Both libraries used four guide RNAs (gRNAs) per target at elevated multiplicity of infections (MOI)^41,42^. By design, library 2 leveraged as-caQTLs, which capture direct allelic regulatory events and minimize biases from linkage disequilibrium (LD). Using SCEPTRE^43^, which is well calibrated for high-MOI screens, to link REs to cis-target genes within 1 Mb, library 1 identified 24 significant SNV-gene pairs across 15 SNVs, nominating 21 CRISPRi genes (ciGenes) (**Supplementary Figure 6J, Supplementary Table 13)**. Library 2 identified 55 significant SNV–gene pairs across 50 REs (59 SNVs), totaling 48 unique ciGenes (FDR < 10%) **(Figure 4B, Supplementary Table 14)**. Unless otherwise noted, downstream analyses refer to library 2 results.

**Figure 4:**
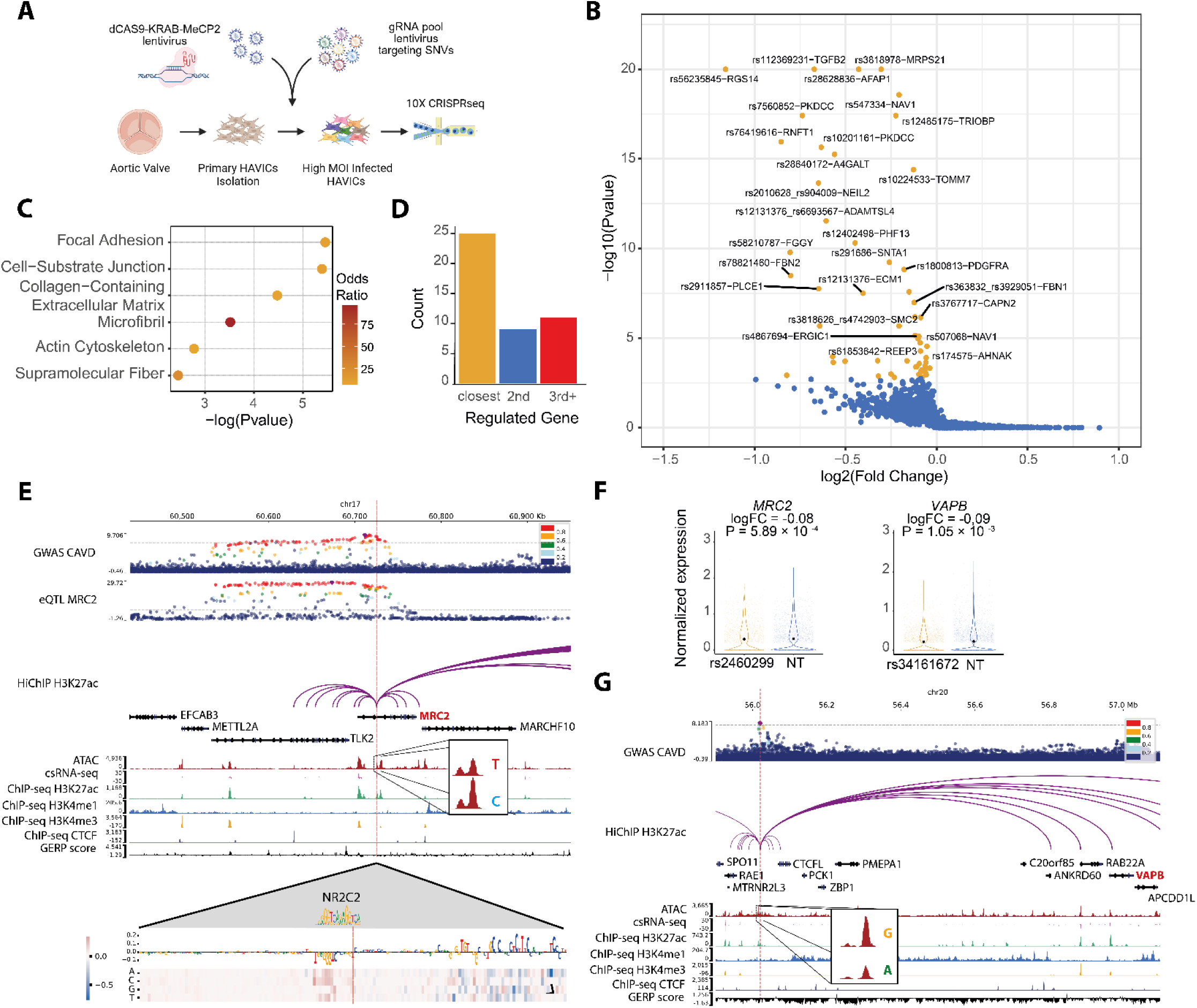
Single-cell CRISPRi perturbation of as-caQTLs identifies cis-target genes at risk loci. **A**, Experimental design for CRISPRi perturbation in primary HAVICs stably expressing KRAB-dCas9-MeCP2, with gRNA libraries targeting SNVs. **B**, Volcano plot of the second gRNA library; yellow points represent significant SNV–gene pairs (SCEPTRE, FDR < 10%). **C**, Gene Ontology enrichment of the 48 identified ciGenes (hypergeometric test). **D**, Frequency of ciGene position relative to the targeted variant (closest gene, second closest, or third and beyond). **E**, CRISPRi of rs2460299-RE downregulates *MRC2* (log₂FC = −0.08, P = 5.89E-4). Tracks show, from top to bottom: GWAS LocusZoom, *MRC2* eQTL LocusZoom, H3K27ac HiChIP loops, gene annotations, ATAC-seq, 5′csRNA-seq, H3K27ac, H3K4me1, H3K4me3 and CTCF ChIP-seq, and GERP conservation score. Insets show allelic imbalance at rs2460299 for ATAC-seq. In silico saturation mutagenesis predicts disruption of an NR2C2 repressor binding motif. **F**, Violin plots of *MRC2* (left) and *VAPB* (right) expression in cells targeted by gRNAs against the respective variant-associated regulatory elements compared to non-targeting controls. **G**, CRISPRi of rs34161672-RE downregulates *VAPB* (log₂FC = −0.09, P = 1.05E-3). Tracks show, from top to bottom: GWAS LocusZoom, H3K27ac HiChIP loops, gene annotations, ATAC-seq, 5′csRNA-seq, H3K27ac, H3K4me1, H3K4me3 and CTCF ChIP-seq, and GERP conservation score. Insets show allelic imbalance at rs34161672 for ATAC-seq.

Library 2 ciGenes were enriched for focal adhesion and collagen-containing ECM **(Figure 4C)**, and for ∼45% of REs, the ciGene was not the closest gene **(Figure 4D)**. **Supplementary Figure 8** summarizes the as-caQTL CRISPRi screen results.

### CRISPRi validates TWAS/MR-nominated genes

Library 1 confirmed several previously nominated candidate genes for CAVD. CRISPRi validated *PALMD*^44^ (via rs6702619-RE, ∼65 kb downstream, consistent with previous CRISPRa analysis), *NAV1*^45^ (via rs507068-RE, P_GWAS_ = 1.18E-27, with an additional ciGene *RNPEP* not annotated by eQTL), and *ATP13A3*^34^ (via rs1706003, log2FC = −0.84, P_SCEPTRE_ = 1.81E-67), a polyamine transporter, along with two additional ciGenes (*TMEM44*, *LINC00884*). Notably, the CAVD risk allele G-rs1706003 is associated with higher *ATP13A3* expression in bulk AV (β = 0.20, P_eQTL_ = 2.39E-20) but trends in the opposite direction in HAVICs (β = −0.097, P_eQTL_ = 0.13), highlighting possible cell-type-specific regulatory effects.

### Perturbation at loci with eQTL support

Among as-caQTLs targeted in library 2 with existing eQTL support, 117 eGenes had been nominated. CRISPRi resolved 48 ciGenes, of which 19 overlapped with eGenes and 29 represented novel target gene assignments.

CRISPRi of rs2460299-RE downregulated *MRC2* (log₂FC = −0.08, P_SCEPTRE_ = 5.8E-4). Rs2460299 (P_GWAS_ = 1.63E-9) is in strong LD (r² = 0.99) with CAVD lead SNV rs2465417 **(Figure 4E–F)**. The C allele increases chromatin accessibility (effect size = 0.68, P_as-caQTL_ = 2.20E-6) and *MRC2* expression in AV (β = 0.17, P_eQTL_ = 4.40E-24), and is predicted to disrupt an NR2C2 repressor binding site, corroborated by in silico saturation mutagenesis. CRISPRi of intronic rs12485175-RE, a CAVD GWAS variant (P_GWAS_ = 7.81E-15), downregulated *TRIOBP* (log₂FC = −0.22, P_SCEPTRE_ = 3.91E-18) and *LGALS1* (log₂FC = − 0.03, P_SCEPTRE_ = 4.7E-4). TRIOBP stabilizes F-actin^46^, whereas LGALS1 (galectin) has been associated with inflammation, ECM remodelling and mineralization in CAVD^47^.

### Perturbation resolves cis-target genes at loci without eQTL support

Nearly half of ciGene associations (29/59) were either absent from eQTL datasets (AV or HAVICs) (19%, 11/59) or identified a different target gene (30%, 18/59), underscoring that perturbation screens at as-caQTLs resolve regulatory relationships and target genes^36^.

Among loci lacking eQTLs, the promoter variant rs1800813 (P_GWAS_ = 1.39E-19; LD r² = 1.0 with the lead SNV) regulates *PDGFRA* (log2FC = −0.17, P_SCEPTRE_ = 1.5E-09) (**Supplementary Figure 9A**). The protective G allele increases chromatin accessibility (effect size = 0.41; P_as-caQTL_ = 1.8E-04) and transcriptional activity in luciferase assay, supporting a regulatory role (**Supplementary Figure 9B**). Consistent with *PDGFRA* as a mesenchymal stemness marker^48,49^, single-cell data across 29,741 HAVICs from 3 donors show it is downregulated during HAVIC-to-myofibroblast differentiation, suggesting the G allele may help preserve stem-like states and limit myofibroblast transition (**Supplementary Figure 9C-F**).

Intergenic rs34161672 is an as-caQTL (P_as-caQTL_ = 8.5E-08) and a lead SNV for CAVD (P_GWAS_ = 1.61E-08) with no eQTL in HAVIC or AV. CRISPRi of rs34161672-RE reduced the expression of *VAPB* (log2FC = −0.09, P_SCEPTRE_ = 0.001), located ∼943 kb downstream, a connection confirmed by long-range H3K27ac-HiChIP loops **(Figure 4F-G)**. *VAPB* encodes a transmembrane endoplasmic reticulum (ER) protein involved in protein export and the unfolded protein response (UPR)^50,51^. Similarly, gRNAs targeting rs477748-RE, a CAVD risk locus (P_GWAS_ = 7.74E-10) with no eQTL (AV, HAVICs), downregulate *ERGIC1*, a component of the ER–Golgi intermediate compartment^52^. Together, these loci implicate intracellular transport and ER homeostasis as functional features of HAVICs linked to CAVD.

In other cases, CRISPRi identified target genes distinct from eQTL-nominated candidates. Rs35613341, a lead CAVD SNV (P_GWAS_ = 9.67E-13) intronic to *ZFPM1*, whose G allele increases chromatin accessibility (effect size = 0.43, P_as-caQTL_ = 1.4E-03) and is associated with higher *ZFPM1* expression in AV (β = 0.27, P_eQTL_ = 2.06E-28). However, CRISPRi of this rs35613341-RE regulated not *ZFPM1* but *SLC7A5*, located ∼625 kb downstream. The above data are consistent with experimental evidence showing that SLC7A5 is involved in fibrosis of cardiac fibroblasts^53^.

### Genetic control of aortic root size and CAVD

The regulatory variant rs6693567 (LD r² = 0.90 with lead SNV rs4970935, within PICS credible set) is associated with aorta flow velocity at the STJ (AoVmax, P_GWAS_ = 6.0E-9) and, consistent with Bernoulli’s principle, indexed aortic area at the STJ (iAoAstj, P_GWAS_ = 5.5E-6). This variant is also associated with CAVD (P_GWAS_ = 3.9E-13), pulse pressure (P_GWAS_ = 3.5E-14), coronary artery disease (P_GWAS_ = 2.9E-8) and coronary artery dissection (P_GWAS_ = 9.7E-29), identifying it as a pleiotropic regulatory hotspot linking hemodynamics, vascular stiffness, and cardiovascular pathology (**Figure 5A**).

**Figure 5:**
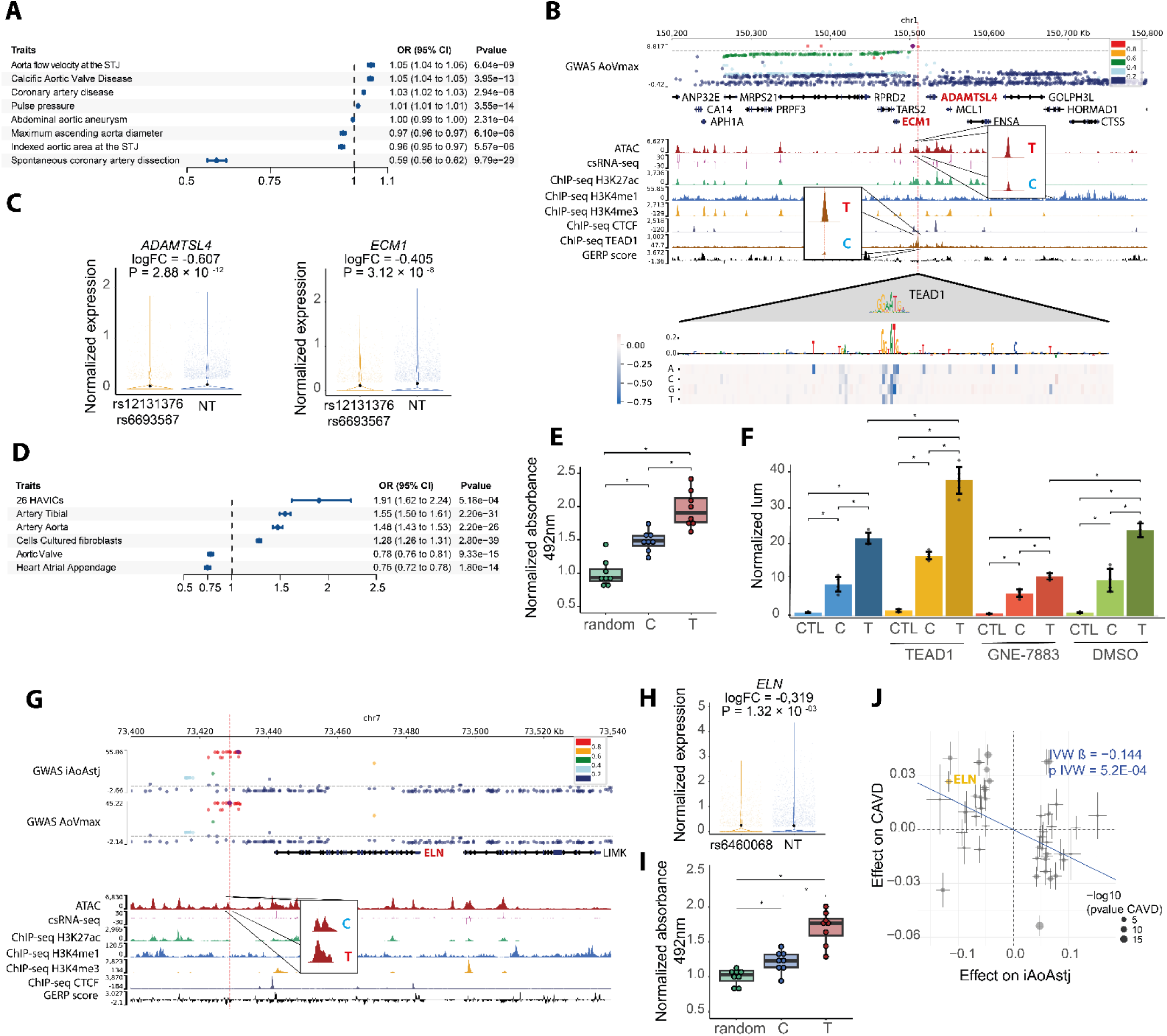
Regulatory variants at elastogenesis loci control aortic root size and CAVD risk. **A**, Forest plot of rs6693567 associations with cardiovascular traits, including aortic flow velocity at the STJ, CAVD, coronary artery disease, pulse pressure, abdominal aortic aneurysm, maximum ascending aorta diameter, indexed aortic area at the STJ and coronary artery dissection (Effect Allele = T). **B**, CRISPRi of rs12131376/rs6693567-RE downregulates *ADAMTSL4* (log₂FC = −0.607, P = 2.88E-12) and ECM1 (log₂FC = −0.405, P = 3.12E-8). Tracks show, from top to bottom: aortic Vmax GWAS LocusZoom, H3K27ac HiChIP loops, gene annotations, ATAC-seq, 5′csRNA-seq, H3K27ac, H3K4me1, H3K4me3 and CTCF ChIP-seq, and GERP conservation score. In silico saturation mutagenesis predicts disruption of a TEAD1 motif. Insets show allelic imbalance at rs6693567 for ATAC-seq (top) and TEAD1 ChIP-seq (bottom). **C** Violin plots of *ADAMTSL4* (left) and *ECM1* (right) expression in cells targeted by gRNAs against the variant-associated regulatory element compared to non-targeting control cells. **D**, Forest plot of rs6693567 eQTL effects on *ADAMTSL4* expression across cardiovascular tissues (Effect Allele = T). **E**, DPI-ELISA showing allele-dependent TEAD1 binding at rs6693567 (n =6, Wilcoxon rank sum test, *P<0.05). **F**, Luciferase reporter assay comparing a random sequence control to a 200-bp enhancer element carrying the C or T allele. Blue: control; yellow: TEAD1 overexpression; red: GNE-7883 treatment; green: DMSO vehicle. (Wilcoxon rank sum test, n=6, 2 donors, *P<0.05). **G**, CRISPRi of rs6460068-RE downregulates *ELN* (log₂FC = −0.319, P = 1.32E-3). Tracks show, from top to bottom: indexed aortic area at the STJ GWAS LocusZoom, aortic Vmax GWAS LocusZoom, gene annotations, ATAC-seq, 5′csRNA-seq, H3K27ac, H3K4me1, H3K4me3 and CTCF ChIP-seq, and GERP conservation score. Inset shows allelic imbalance for ATAC-seq. **H**, Violin plot of *ELN* expression in cells targeted by gRNAs against the variant-associated regulatory element compared to non-targeting controls. **I**, DPI-ELISA showing allele-dependent GATA4 binding at rs6460068 (n=6, Wilcoxon rank sum test, *P<0.05). **J**, Mendelian randomization of the aortic root area at the STJ on CAVD showing that genetically determined smaller iAoAstj is associated with increased CAVD risk (β = −0.14, P_IVW = 5.3E-4).

CRISPRi of rs6693567-RE reduced expression of *ADAMTSL4* (log2FC = −0.60, P_SCEPTRE_ = 2.8E-12) and *ECM1* (log2FC = −0.40, P_SCEPTRE_ = 3.1E-8) (**Figure 5B-C**). Notably, the T allele of rs6693567 was associated with higher *ADAMTSL4* expression in HAVICs (β = 0.64, P_eQTL_ = 5.1E-04), consistent with GTEx data in vascular tissues and fibroblasts, but with lower expression in bulk AV tissue (β = −0.24, P_eQTL_ = 9.3E-15), mirroring the cell-type-specific directional effects observed at some loci and likely reflecting opposing regulatory contributions from non-HAVIC/mesenchymal cell populations (**Figure 5D**). Rs6693567 is predicted to alter a TEAD1 binding motif, with the T allele strengthening TEAD1 recruitment, a prediction supported by Enformer in silico saturation mutagenesis. Experimentally, DNA-protein interaction (DPI)-ELISA confirmed allele-dependent TEAD1 binding (**Figure 5E)**, and TEAD1 ChIP-seq in HAVICs revealed strong allele-specific binding at this position (effect size = 3.76, P_TF-ASB_ = 5.1E-18) (**Figure 5B, Supplementary Table 4**). Consistent with HAVIC eQTL direction, reporter assay showed T allele-dependent activity, which was enhanced by TEAD1 overexpression and reduced by pharmacological inhibition of TEAD with GNE-7883^54^ (**Figure 5F**). The variant lies in a conserved enhancer (GERP = 4.7), suggesting evolutionary constraints and developmental gene regulation. In fetal cardiac stromal cells, single cell ATAC-seq co-accessibility links rs6693567-RE to the *ADAMTSL4* promoter^55^, and in the human fetal heart (week 11)^56^ **(Supplementary Figure 10A)**, *ADAMTSL4* and elastogenesis genes (*ELN*, *FBLN5*, *LOX*, *FBN1*) are enriched in the outflow tract, a region involved in the development of the AV (**Supplementary Figure 10B-F**). These findings are consistent with data showing that ADAMTSL4 binds to FBN1 and control microfibril assembly and elastogenesis^57^.

At the *ELN* locus, intergenic as-caQTL rs6460068 is strongly associated with ascending aorta size (P_GWAS_ = 1.80E-76) and iAoAstj (P_GWAS_ = 2.00E-53), and suggestively with CAVD (P_GWAS_ = 4.15E-06), yet had no eQTL in HAVIC or AV. CRISPRi of rs6460068-RE resolved this gap, identifying *ELN* as the target gene (log2FC = −0.31, P_SCEPTRE_ = 0.001) (**Figure 5G-H**). The T allele creates a GATA4 binding motif, confirmed by DPI-ELISA **(Figure 5I)**. Thus, both the *ADAMTSL4* and *ELN* loci converge on elastogenesis, *ADAMTSL4* controlling microfibril and elastin fibre assembly and *ELN* encoding its principal substrate, each under allele-specific regulatory control in HAVICs. This convergence, together with the shared pleiotropy between aortic root size and CAVD across multiple loci, prompted us to formally test whether aortic root geometry causally influences disease risk. Mendelian randomization supported a bidirectional relationship between iAoAstj and CAVD. Genetically predicted iAoAstj was inversely associated with CAVD risk (β = −0.14, P_IVW_ = 5.3E-04) **(Figure 5J, Supplementary Table 15, Supplementary Figure 10G)**, whereas genetically predicted CAVD was associated with lower iAoAstj (β = −0.06, P_IVW_ = 1.3E-03) **(Supplementary Table 15)**. Collectively, these findings implicate impaired elastogenesis as a determinant of reduced aortic root dimensions, thereby increasing susceptibility to CAVD.

### Convergence of common regulatory and rare coding variants at fibrillin genes

Given its high haploinsufficiency score (pHaplo = 0.99)^58^, *FBN1* harbors one of the strongest rare loss-of-function (LOF) variant burden associations with cardiovascular disease in UK Biobank whole exome sequencing (WES) data^59^, including congenital malformation syndromes (P_burden_ = 2.01E-27), aortic aneurysm and dissection (P_burden_ = 6.86E-11), aortic valve repair or replacement (P_burden_ = 5.55E-7), and nonrheumatic aortic valve disorders (P_burden_ = 6.08E-6). Clinically, pathogenic mutations in *FBN1* cause Marfan syndrome^60^. We tested whether common GWAS variants also converge on *FBN1* through regulatory mechanisms. CRISPRi identified rs363832/rs3929051-RE as the strongest RE controlling *FBN1* expression (log2FC = −0.12, P_SCEPTRE_ = 1.03E-7) (**Supplementary Figure 11A**). These two as-caQTLs are in strong LD (r² = 0.97) within the same intronic ATAC peak. The rs363832-C allele increases pulse pressure (P_GWAS_ = 2.82E-20), decreases iAoAstj (P_GWAS_ = 4.98E-7), and is predicted to disrupt a motif for MEIS2, a homeobox TF involved in aortic root development^61^.

At 5q23, CRISPRi of rs73348278-RE and rs78821460-RE reduced expression of *FBN2* (log2FC = −0.64, P_SCEPTRE_ = 2.1E-6 and log2FC = −0.80, P_SCEPTRE_ = 3.2E-09) (**Supplementary Figure 11B**), which encodes the *FBN1* paralog fibrillin-2 with overlapping functions in microfibril assembly^62,63^. The 73348278-C allele increases chromatin accessibility (effect size = 1.13, P_as-caQTL_ = 1.78E-4) and *FBN2* expression in AV (β = 1.14, P_eQTL_ = 9.34E-69), and is associated with lower CAVD risk (P_GWAS_ = 7.52E-9). Together, these results demonstrate convergence between rare coding mutations and common regulatory variants at fibrillin genes in controlling the development of the AV apparatus and risk of CAVD^64,65^.

### Identifying causal programs for aortic valve disease

Genome-wide, CRISPRi of variants can resolve trans-regulated ciGenes. For a perturbation, we considered trans-networks consisting in at least 50 ciGenes for downstream analysis. To link trans-regulated ciGenes to disease, co-expressed downstream ciGenes were grouped into functional programs^41^. Inspired by recent integrative genetic–perturbation frameworks^66^, we used LOF burden associations from UK Biobank whole-exome sequencing (WES) to derive a genetically informed perturbation program score (GIPPS). GIPPS harmonizes association effects to perturbation direction, providing a signed empirical –log10 P-value summarizing program-level effects on disease risk (|GIPPS| > 1.30; P < 0.05) **(Methods, Supplementary Figure 12A)**. In total we identified seven programs from upstream variant perturbations, of which three were core programs causally associated to CAVD (|GIPPS| > 1.30).

CRISPRi of rs174575-RE (P_GWAS_ = 2.80E-26), in moderate LD (r² = 0.55) with the lead SNV at a CAVD risk locus, regulated *FADS2* (log2FC = −0.15, P_SCEPTRE_ = 2.64E-8) and *AHNAK* (log2FC = −0.05, P_SCEPTRE_ = 2.88E-5) in cis, the latter connected by long-range HiChIP loops ∼721 kb downstream (**Figure 6A-B**). This locus produced a large trans-network of 153 ciGenes (**Figure 6C, Supplementary Table 16**). Co-expression analysis revealed three programs, of which only the green program (59 ciGenes including cis-regulated gene *AHNAK*), enriched in ECM organization (OR = 25.75, P = 2.7E-16, hypergeometric test) (**Figure 6D**), was significantly associated with CAVD risk (GIPPS = −1.54), suggesting that rs174575-AHNAK-controlled program regulates fibrogenesis (**Figure 6E**). All ciGenes in this program were downregulated by CRISPRi, including myofibroblast markers (*MYH9*, *CNN1*, *FN1*). This program was enriched for SMAD3^67^ ChIP peaks from ChIP-Atlas^68^ (OR = 3.44, P = 0.002) and TEAD1 peaks in HAVICs (OR = 9.42, P_TF-ASB_ = 2.37E-11) (**Supplementary Figure 13A**). To assess whether TEAD1 directly controls trans-ciGenes, we designed gRNAs to the *TEAD1* promoter, identifying 88 genome-wide ciGenes. Twenty ciGenes from the green AHNAK-program were among the TEAD1 targets (OR = 62.55, P = 1.11E-31) **(Supplementary Table 17)**, including six with TEAD1 ChIP peak enrichment (*CLIC4*, *SPARC*, *VIM*, *COL4A2*, *COL1A1*, *MYH9*). In HAVICs, CRISPRi of *AHNAK* reduced TGF-β-induced collagen production in the scar-in-a-jar assay, providing direct functional evidence that AHNAK controls an ECM program in accordance with the direction of GIPPS (**Figure 6F, Supplementary Figure 13B-C**).

**Figure 6:**
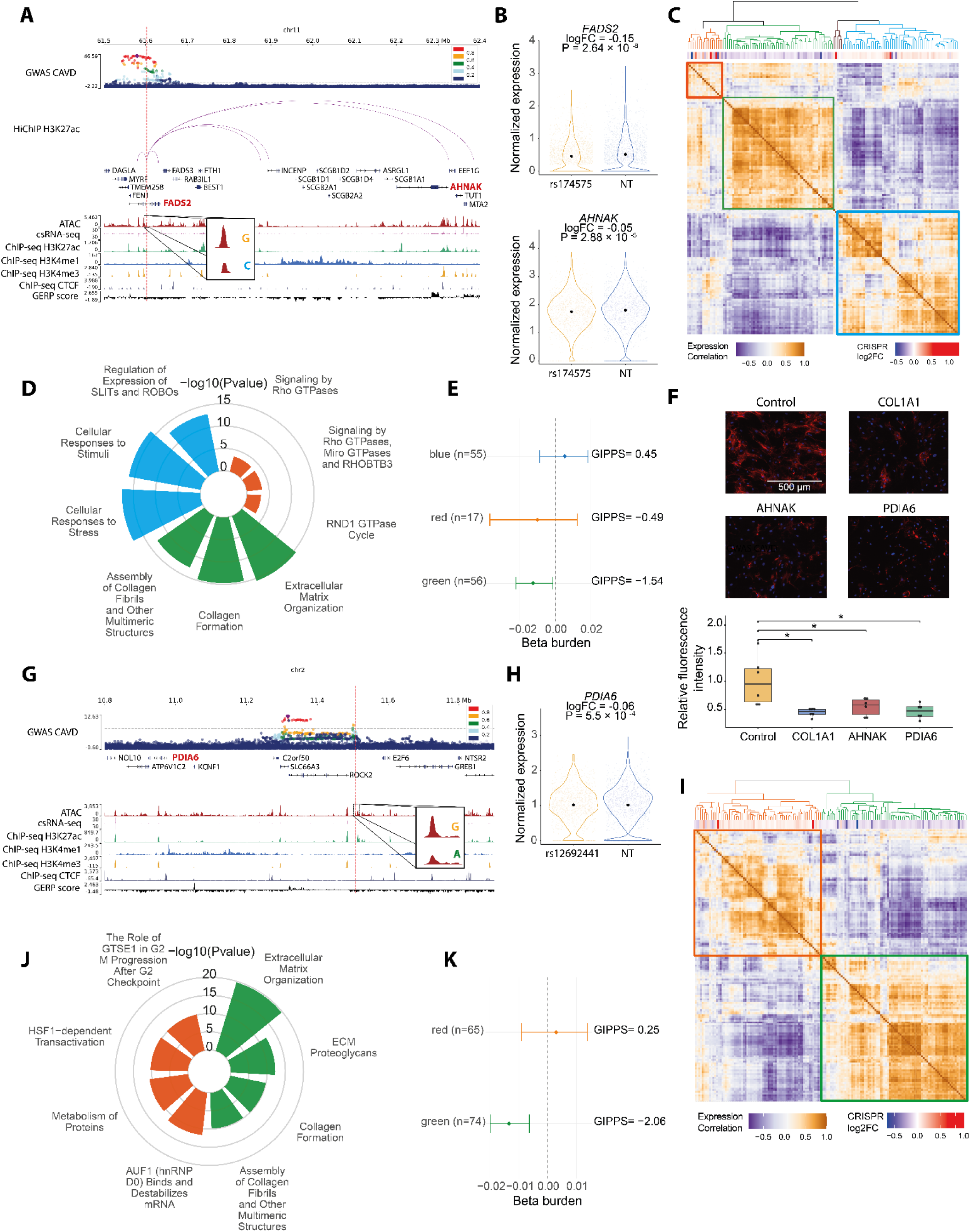
CRISPRi perturbation identifies upstream regulators of core HAVIC programs causal for CAVD. **A**, CRISPRi of rs174575-RE downregulates *FADS2* (log₂FC = −0.15, P = 2.64E-8) and *AHNAK* (log₂FC = −0.05, P = 2.88E-5). Tracks show, from top to bottom: CAVD GWAS LocusZoom, H3K27ac HiChIP loops, gene annotations, ATAC-seq, 5′csRNA-seq, H3K27ac, H3K4me1, H3K4me3 and CTCF ChIP-seq, and GERP conservation score. Inset shows allelic imbalance for ATAC-seq. **B**, Violin plots of *FADS2* (top) and *AHNAK* (bottom) expression in cells targeted by gRNAs against the variant-associated regulatory element compared to non-targeting controls. **C**, Heatmap of co-expression among 153 trans-regulated ciGenes downstream of rs174575-RE perturbation, with CRISPRi effect sizes and hierarchical clustering identifying three programs. **D**, Reactome pathway enrichment analysis for each program (top three enrichments per program; hypergeometric test). **E**, Forest plot of the genetically informed perturbation program score (GIPPS) for the three programs; the ECM organization program is significantly associated with CAVD risk (GIPPS = (log₂FC = −0.05, P = 2.88E-5). Tracks show, from top to 1.54). Number of genes overlapping each program and LOF data for GIPPS analysis was reported. **F**, Scar-in-a-jar collagen production assay following CRISPRi perturbation with non-targeting, *COL1A1*, *AHNAK* and *PDIA6* gRNAs (n = 6 replicates from 2 donors) (Wilcoxon signed rank test, *P<0.05). **G**, CRISPRi of rs12692441-RE downregulates *PDIA6* (log₂FC = (log₂FC = −0.05, P = 2.88E-5). Tracks show, from top to 0.06, P = 5.5E-4). Tracks show, from top to bottom: CAVD GWAS LocusZoom, gene annotations, ATAC-seq, 5′csRNA-seq, H3K27ac, H3K4me1, H3K4me3 and CTCF ChIP-seq, and GERP conservation score. Inset shows allelic imbalance for ATAC-seq. **H**, Violin plot of *PDIA6* expression in cells targeted by gRNAs against the variant-associated regulatory element compared to non-targeting control cells. **I**, Heatmap of co-expression among 160 trans-regulated ciGenes downstream of rs12692441-RE perturbation, with CRISPRi effect sizes and hierarchical clustering identifying two programs. **J,** Reactome pathway enrichment analysis for each program (top three enrichments per program; hypergeometric test). **K**, Forest plot of the genetically informed perturbation program score (GIPPS) for the two programs; the ECM program (green) is associated with CAVD risk (GIPPS = (log₂FC = −0.05, P = 2.88E-5). Tracks show, from top to 2.06). Number of genes overlapping each program and LOF data for GIPPS analysis was reported.

CRISPRi of rs12692441-RE, an intergenic as-caQTL associated with CAVD at suggestive threshold (P_GWAS_ = 1.06E-6) with no eQTL (AV, HAVIC), reduced in cis *PDIA6* (∼530 kb upstream), a gene involved in the UPR^69^, and trans-regulated 160 ciGenes (**Figure 6G-I, Supplementary Table 18**). Co-expression analysis delineated two programs: a heat shock response program (red) and an ECM program (green) containing the collagen chaperone *FKBP10* as top downregulated gene (**Figure 6J**). Only the ECM program was associated with CAVD risk (GIPPS = (log₂FC = −0.05, P = 2.88E-5). Tracks show, from top to 2.06) (**Figure 6K**). Consistent with the direction of GIPPS, CRISPRi of *PDIA6* reduced collagen production in the scar-in-a-jar assay, providing functional evidence that PDIA6 controls ECM-program and fibrogenesis (**Figure 6F, Supplementary Figure 13B-D**).

CRISPRi of rs76419616-RE associated with CAVD (P_GWAS_ = 3.87E-6), downregulated in cis *RNFT1* (log2FC = (log₂FC = −0.05, P = 2.88E-5). Tracks show, from top to 0.85, P_SCEPTRE_ = 1.11E-16), encoding a E3 ubiquitin-protein ligase involved in the ER-associated degradation pathway (ERAD)^70^, and impacted genome-wide expression of 103 ciGenes (**Supplementary Figure 13E, Supplementary Table 19)**. Co-expression analysis identified two programs (blue and red) **(Supplementary Figure 13F)**. Only the blue program, which contains *RNFT1* and is enriched in iron uptake and transport (OR: 37.5, P = 5.31E-07, hypergeometric test) **(Supplementary Figure 13G)**, was associated with CAVD (GIPPS = 2.41) **(Supplementary Figure 13H)**. These findings are consistent with evidence that heme-derived iron released from intra-leaflet microhemorrhages during CAVD impact HAVIC function and support a role for ER-stress in cell iron handling^71^.

## Discussion

The findings of this work show that as-caQTLs in HAVICs reflect direct regulatory events associated with TF binding and are enriched for genetic variants contributing to CAVD heritability. By integrating genomic annotations with CRISPRi perturbation screens targeting prioritized variants displaying allelic imbalance, we identified cis-regulated target genes at loci relevant to AV function. This approach also revealed mechanistic links between structural features, such as aortic root size and CAVD risk. In parallel, perturbation experiments uncovered HAVIC core regulatory programs and their upstream regulators associated with CAVD **(Supplementary Figure 14)**.

as-caQTLs provide a direct readout of molecular events regulating chromatin accessibility, while mitigating confounding effects such as LD^72^. Although as-caQTLs offer a robust framework for identifying causal regulatory mechanisms, their application is limited to heterozygous variants present in the sampled individuals. To overcome this limitation and extend allelic imbalance predictions to unseen sequences, we leveraged knowledge transfer with the Enformer model, which predicted strong regulatory effects for both common and rare variants. When combined with HAVIC-specific functional annotations, TF allelic imbalance analyses, and CRISPR perturbation screens, as-caQTLs enabled the identification of regulatory mechanisms at several loci. Supported by experimental validation, we identified key TFs, including TEAD1 and GATA4, along with their respective target genes, *ADAMTSL4* and *ELN*. Both loci are associated with aortic root size and CAVD. *ADAMTSL4* plays an important role in microfibril and elastin assembly, highlighting a potential link between ECM organization and disease risk. This relationship between microfibril biology and CAVD is further supported by the presence of rare LOF and common noncoding variants impacting *FBN1* and *FBN2*. Together, these observations raise the possibility that the anatomical structure of the aortic root, determined by microfibril and elastin assembly, may influence susceptibility to CAVD^73^. Consistent with this hypothesis, Mendelian randomization analyses showed that genetically predicted lower aortic root size is associated with an increased risk of CAVD.

A central objective of these perturbation screens was to identify trans-regulated gene networks, define their organization into HAVIC programs, and determine which are causal for CAVD. We identified three variants that robustly regulate seven co-regulated programs through upstream cis-regulated genes. To test causality, we leveraged LOF burden data from UK Biobank WES^59^. Adapting a recently proposed framework^66^, we derived an empirical score (GIPPS) to quantify the relationship between upstream regulatory perturbation and CAVD risk for each program. This approach identified three upstream peripheral regulators controlling core programs converging on ECM organization/production and iron uptake. Using the scar-in-a-jar assay, we validated that ECM programs are regulated by the cis-regulated genes *AHNAK* and *PDIA6*. AHNAK, a large multifunctional protein, regulates an ECM program enriched for SMAD3 and TEAD1. The strong impact of *PDIA6* further implicates the UPR in HAVIC fibrogenesis. Intra-leaflet microhemorrhages, commonly observed in severely remodeled CAVD cusps, release heme-derived iron into the tissue environment^74^. Although iron promotes HAVIC dysfunction, whether iron handling represents a causal mechanism remained unresolved. Here, we identify *RNFT1* as an upstream regulator of a core causal program governing cellular iron uptake and transport. Together, these findings support an omnigenic architecture in which peripheral regulators converge on core gene programs that drive CAVD **(Supplementary Figure 12B)** ^75^.

Over the past five years, GWASs have markedly expanded the number of loci associated with traits and diseases of the AV apparatus. Leveraging these discoveries, we developed an integrative framework in HAVICs to prioritize causal variants, identify their target genes, and define the gene programs mediating CAVD risk. This approach uncovered novel upstream regulators of core programs and revealed links between genetic determinants of aortic valve anatomy/function, ECM biology, and disease susceptibility. These findings provide a mechanistic framework for CAVD genetics and highlight microfibril‒elastin/ECM regulation and iron handling as key pathways.

## ONLINE METHODS

### Cell lines and culture

HEK293T cells (ATCC #CRL-11268) and immortalized human bone marrow mesenchymal stromal cells (iMSC3; ABM #T0529) were cultured in DMEM high glucose (Gibco #11995-065) supplemented with 10% FBS (VWR #97068-085), 1% penicillin–streptomycin (Gibco #15240-062), 0.2% mycozap (Lonza #VZA-2032), 10 mM sodium pyruvate (Gibco #11360-070) and 20 mM L-glutamine (Gibco #25030-081). Cells were passaged using trypsin–EDTA 0.5% (Gibco #15400-054) at a 1:5 dilution.

Human aortic valve interstitial cells (HAVICs) and endothelial cells (HAVECs) were isolated from aortic valves obtained from patients who underwent cardiac surgery at the Institut universitaire de cardiologie et de pneumologie de Québec – Université Laval. All patients provided informed consent, and the study was approved by the IUCPQ ethics committee. Non-mineralized valves were obtained following heart transplantation; mineralized valves were obtained during valve replacement. Leaflets were washed in HBSS 1X and incubated with 2.6 mg/ml type II collagenase (Gibco #171010-015) in complete DMEM at 37°C for 10 min with rotation. HAVECs were detached by gentle rubbing with a cotton swab, pelleted at 200 rcf for 5 min, and resuspended in endothelial cell growth BBE medium (ATCC #PCS-100-030, #PCS-100-040). Leaflets were then minced and incubated with 3 mg/ml type I collagenase (Gibco #17100-017) in HBSS 1X at 37°C for 45 min with rotation. Digested tissue was filtered through a 70 µm mesh, pelleted, and resuspended in complete DMEM. HAVICs and HAVECs were used between passages 3 and 7.

Mouse aortic valve interstitial cells (mAVICs) were isolated from pooled aortic valves of 5–10 C57BL/6 mice. Tissues were subjected to sequential collagenase digestion: first at 1 mg/ml type I collagenase for 30 min, then at 4.5 mg/ml for 30 min, both at 37°C with agitation in a 1:1 mixture of serum-free DMEM and HEPES. Cells were plated at minimal volume in 24-well plates and left undisturbed for several days to maximize adhesion efficiency.

### Lentivirus preparation

Lentiviral particles were produced in HEK293T cells (passage <20) seeded at 7 × 10⁶ cells per 10 cm dish. Transfection was performed in Opti-MEM supplemented with 5% FBS and 200 µM sodium pyruvate using Lipofectamine 3000 (Thermo Fisher Scientific). Per dish, 12 µg transfer plasmid, 9 µg psPAX2, and 3 µg pMD2.G were combined with P3000 reagent (mix A) and Lipofectamine 3000 (mix B), incubated 20–30 min, and added dropwise to cells. Medium was replaced after 4 h and 1/500 ViralBoost Reagent (Alstem #VB100) was added. Viral supernatants were collected at 24 and 48 h post-transfection, pooled, filtered through 0.45 µm, concentrated with Lenti-X concentrator (Takara #631231) following recommendation and either used immediately or snap-frozen at (log₂FC = −0.05, P = 2.88E-5). Tracks show, from top to 80°C. Target cells were transduced in packaging medium supplemented with 8 µg/ml polybrene. Antibiotic selection was initiated 48 h post-transduction and maintained throughout the experiment.

### RNA-seq

HAVICs were isolated from 14 CAVD patients undergoing valve replacement and 13 controls undergoing heart transplantation (10 women, 17 men). HAVECs were obtained from four transplant donors (two women, two men). RNA was extracted using the E.Z.N.A. Total RNA Kit I (Omega Bio-Tek #R6834-02) and quality-assessed on a Bioanalyzer (Agilent RNA 6000 Nano Kit; all RIN ≥9.2). Libraries were prepared from 1 µg RNA using the Illumina Stranded mRNA Prep Ligation Kit (#20040532) and sequenced as 100 bp paired-end reads.

### Genotyping

Genomic DNA was extracted from buffy coats using the QIAamp DNA Blood Midi Kit (Qiagen) and quantified using PicoGreen (Invitrogen #P7589). Samples were diluted to 50 ng/µl and genotyped on the Illumina Infinium Global Screening Array-24 v3.0 at Genome Quebec. All patients provided informed consent, and the study was approved by the IUCPQ ethics committee.

### ATAC-seq

ATAC-seq libraries were prepared from freshly trypsinized HAVICs from 10 control and 6 CAVD valves, following the Kaestner Lab ATAC-seq Protocol and Brunton et al. with modifications^76^. Aliquots of 40,000 cells were lysed in cold lysis buffer (10 mM Tris-HCl pH 7.5, 10 mM NaCl, 3 mM MgCl₂, 0.1% NP-40, 0.1% Tween-20, 0.01% digitonin) for 10 min on ice. Nuclei were pelleted at 1,200 rcf, resuspended in 20 µl of Tn5 transposition reaction mix (Illumina #20034197), and incubated at 37°C for 1–1.5 h at 1,250 rpm. Transposed fragments were purified (MicroElute GEL/PCR Purification Kit, GeneBio Systems), amplified with barcoded primers, and purified using 1.8× SPRIselect beads (Beckman #B23317). Libraries were quantified on a Bioanalyzer High Sensitivity DNA chip and sequenced as 150 bp paired-end reads.

### ChIP-seq

ChIP-seq was performed on 2 million freshly trypsinized HAVICs per replicate (n = 3 biological replicates for histone marks and TEAD1; n = 2 for CTCF), adapted from the RELACS protocol^77^. Cells were cross-linked with 1% formaldehyde for 10 min, quenched with 125 mM glycine, and lysed in nuclear extraction buffer (10 mM Tris-HCl pH 8.0, 10 mM NaCl, 0.2% NP-40, 1× protease inhibitors) for 1 h at 4°C. Nuclei were treated with 0.5% SDS at 62°C for 10 min, quenched with 1.5% Triton X-100, and chromatin was digested with DpnII (NEB #R0543M) for 6 h at 37°C. Chromatin was released by sonication (QSONICA Q800R3, 11 min, 30 s on/30 s off, 30% amplitude, 4°C) and diluted in Diagenode iC1 buffer.

Immunoprecipitation was performed overnight at 4°C using antibodies against H3K27ac (Abcam #ab4729), H3K4me3 (Abcam #ab8580), H3K4me1 (CellSignaling #5326S), H3K27me3 (Diagenode #C15410195), H3K36me3 (Diagenode #C15410192), H3K9me3 (Abcam #ab8898), TEAD1 (Active Motif #61644), and CTCF (Cell Signaling #2899). Immune complexes were captured with Dynabeads Protein G (Invitrogen #10004D) for 3 h and sequentially washed with Diagenode iW1–iW4 buffers. DNA was eluted, treated with RNase A and Proteinase K, de-cross-linked overnight at 65°C, and purified using the Zymo ChIP DNA Clean & Concentrator Kit (#D5205).

Libraries were prepared from 50 µl purified DNA using the NEBNext Ultra II DNA Library Prep Kit (NEB #E7645S) with modifications: size selection was omitted and adapter-ligated fragments were purified using 1× SPRIselect beads. USER enzyme treatment and PCR enrichment (12 cycles) were performed in a single step. Libraries were sequenced as 150 bp paired-end reads.

### Hi-C

Hi-C was performed on 2 million HAVICs, adapted from Lafontaine et al. (2021)^78^ and Gryder et al. (2020)^79^. Cells were cross-linked with 1% formaldehyde and processed as for ChIP-seq through DpnII digestion. After heat inactivation of DpnII (62°C, 20 min), digested ends were filled in with biotin-14-dATP (Invitrogen #19524016) using Klenow fragment (NEB #M0210) at 37°C for 1 h. Proximity ligation was performed overnight at room temperature with T4 DNA Ligase (NEB #M0202) in the presence of 1% Triton X-100. Nuclei were pelleted, resuspended in shearing buffer, and sonicated (QSONICA Q800R3, 7 min, 30 s on/30 s off, 30% amplitude, 4°C). DNA was treated with RNase A and Proteinase K, de-cross-linked at 67°C for 3 h, and purified using the Zymo ChIP DNA Clean & Concentrator Kit.

Biotin pull-down was performed using Streptavidin C-1 beads (2 µl/µg DNA) in 2× Biotin Binding Buffer (10 mM Tris-HCl pH 7.5, 1 mM EDTA, 2 M NaCl) for 15 min at room temperature. Libraries were prepared on-bead using the NEBNext Ultra II DNA Library Prep Kit (NEB #E7645S) and purified using 1× SPRIselect beads. Libraries were deeply sequenced as 150 bp paired-end reads.

### HiChIP

HiChIP was performed on 1 million freshly trypsinized HAVICs from four biological replicates, adapted from Mumbach et al. (2016)^80^. Cells were cross-linked with 1% formaldehyde for 10 min at room temperature, quenched with 125 mM glycine, and lysed in nuclear extraction buffer (10 mM Tris-HCl pH 8.0, 10 mM NaCl, 0.2% NP-40, 1× protease inhibitors) for 1 h at 4°C. Nuclei were treated with 0.5% SDS at 62°C for 10 min, quenched with 1.5% Triton X-100, and chromatin was digested with MboI (NEB #R0147) for 2 h at 37°C. After heat inactivation (62°C, 20 min), digested ends were filled in with biotin-14-dATP (Invitrogen #19524016) using Klenow fragment (NEB #M0210) at 37°C for 1 h. Proximity ligation was performed with T4 DNA Ligase (NEB #M0202) for 4 h at room temperature in the presence of 1% Triton X-100.

Nuclei were pelleted, resuspended in nuclear lysis buffer (50 mM Tris-HCl pH 7.5, 1% SDS, 10 mM EDTA), and chromatin was released by sonication (QSONICA Q800R3, 3 min, 30 s on/30 s off, 20% amplitude, 9°C). Sonicated chromatin was diluted in ChIP dilution buffer and immunoprecipitated overnight at 4°C with 4 µg anti-H3K27ac (Abcam #ab4729) and Dynabeads Protein G (Invitrogen #10004D). Beads were sequentially washed with low-salt, high-salt, and LiCl wash buffers, and DNA was eluted in 50 mM NaHCO₃/1% SDS, treated with Proteinase K, de-cross-linked at 67°C, and purified using the Zymo ChIP DNA Clean & Concentrator Kit (#D5205).

Biotinylated ligation junctions were captured on Streptavidin C-1 beads in 2× Biotin Binding Buffer (10 mM Tris-HCl pH 7.5, 1 mM EDTA, 2 M NaCl) for 15 min at room temperature. On-bead tagmentation was performed with Tn5 (Illumina #20034198) at 55°C for 10 min, with the Tn5 amount scaled to input DNA as described in Mumbach et al. (2016). Beads were washed sequentially with 50 mM EDTA, Tween Wash Buffer, and 10 mM Tris pH 7.5, then resuspended in PCR master mix containing Phusion HF (NEB #M0531S) and Nextera indexed primers. Libraries were amplified (72°C for 5 min; 98°C for 1 min; cycles of 98°C for 15 s, 63°C for 30 s, 72°C for 1 min), purified using the Zymo DNA Clean & Concentrator Kit, quantified by Qubit and Bioanalyzer, and sequenced as 150 bp paired-end reads.

### 5′-capped small RNA-seq (5′csRNA-seq)

Transcription initiation profiling was performed on HAVICs from 7 donors (5 controls, 2 CAVD), following the Meyer et al. protocol^81^ with minor modifications. Total RNA was extracted with Qiazol (Qiagen #79306) and chloroform, precipitated with isopropanol overnight at (log₂FC = −0.05, P = 2.88E-5). Tracks show, from top to 20°C, and quality-assessed on a Bioanalyzer (all RIN ≥9.2). Small RNAs (∼20–55 nt) were size-selected on a 15% TBE-urea gel (Bio-Rad #4566055), excised, eluted in 0.5 M NaCl, and precipitated with ethanol.

5′-cap enrichment was performed by sequential treatment with Terminator exonuclease (LGC Genomics #TER51020; 30°C, 1 h) and CIP phosphatase (NEB #M0525S; 37°C, 1 h plus 45 min), following Meyer et al. (2026). RNA was purified with Trizol LS/chloroform extraction and isopropanol precipitation. Libraries were prepared using the NEBNext Small RNA Library Prep Kit (NEB #E7330S), with RppH-mediated decapping (37°C, 2 h), sequential 3′ and 5′ adapter ligation, reverse transcription, and PCR amplification (14 cycles). Libraries were size-selected on a 10% TBE gel (140–175 bp), eluted in diffusion buffer, and precipitated with ethanol. Sequencing was performed as 150 bp paired-end reads.

Total RNA-seq control libraries for csRNA-seq normalization were prepared from 500 ng total RNA using the Illumina Stranded Total RNA Prep Ligation Kit with Ribo-Zero Plus (#20040529).

### RT-qPCR

Total RNA was extracted using the E.Z.N.A. Total RNA Kit (Omega Bio-Tek) and quantified by Nanodrop. Reverse transcription was performed on 1 µg RNA using the qScript cDNA Synthesis Kit (Quantabio). Quantitative PCR was performed on a Corbett Rotor-Gene 6000 using PerfeCTa SYBR Green SuperMix (Quantabio) with 5 µl of 1:10-diluted cDNA per reaction and 2 µl of Qiagen primers (HPRT1 #QT00059066, AHNAK #QT01680994, PDIA6 #QT00037086). Cycling conditions: 95°C for 15 min; 40 cycles of 94°C for 10 s, 55°C for 30 s, 72°C for 30 s. Relative expression was quantified using the ΔΔCt method normalized to HPRT1 housekeeping gene.

### Perturb-seq

Perturb-seq was performed on HAVICs from 2 donors, following the CROPseq^82^ protocol for library construction and the STING-seq^41^ protocol for the plasmid and the 10x single cell 5’ CRISPR capture.

*Guide RNA library construction.* A custom sgRNA library designed with Flashfry^83^ targeting candidate regulatory elements was synthesized as oligonucleotide pools (IDT (library1) or Agilent HiFi OLS (library2)). The lentiGuide-FE-Puro backbone (Addgene #170069) was linearized with BsmBI and purified according to Datlinger protocol^82^. Oligonucleotides were cloned by Gibson assembly (NEBuilder HiFi DNA Assembly Master Mix, 1:200 vector:insert molar ratio, 50°C, 1 h), desalted on a 0.05 µm MCE membrane, and electroporated into Lucigen Endura cells (BioRad Micropulser, EC1 program). Transformants were plated at ∼100× library coverage and plasmid DNA was extracted by maxiprep.

*Library quality control*. sgRNA representation was verified by targeted sequencing on an Illumina MiniSeq, following Datlinger recommendations. The sgRNA cassette was amplified from the plasmid pool (10 ng/µl), with optimal cycle number determined by qPCR to prevent overamplification. Products were purified using 2.0× SPRI beads and quantified by Qubit HS dsDNA assay.

*CRISPRi perturbation and single-cell capture*. HAVICs were co-transduced with lentiviral particles carrying dCas9-KRAB-MeCP2 (lentiCRISPRi(v2)-Blast, Addgene #170068; MOI ∼ 5) and the sgRNA library at the desired MOI. Selection with puromycin and blasticidin was initiated 48 h post-transduction and maintained for 10 days. Single-cell gene expression and sgRNA capture libraries were prepared using the Chromium Next GEM Single Cell 5′ Kit v2 with the Chromium Next GEM Chip K Single Cell Kit (10x Genomics). sgRNA-derived cDNA was separated from gene expression cDNA by SPRIselect size selection according to 10x gRNA capture protocol. Both libraries were sequenced as 150 bp paired-end reads.

### Single-cell RNA-seq of TGFβ-induced myofibroblast differentiation

To profile myofibroblast differentiation at single-cell resolution, HAVICs from three independent donors were cultured in Fibroblast Growth Medium 3 (PromoCell #C-23025) and treated with either 10 ng/ml TGFβ1 (Gibco #PHG9214) for 48 h or vehicle control. Cells were then harvested and processed for single-cell RNA sequencing using the Chromium Next GEM Single Cell 3′ Kit v3.1 with the Chromium Next GEM Chip G Single Cell Kit (10x Genomics), following the manufacturer’s instructions. Libraries were sequenced on an Illumina NextSeq 2000 P3.

### DPI-ELISA

Transcription factor–DNA binding was assessed by streptavidin-capture ELISA^84^. Complementary biotinylated oligonucleotides (30 nt centered on the SNV, IDT) were annealed at 2 µM in annealing buffer (40 mM Tris-HCl pH 8.0, 20 mM MgCl₂, 50 mM NaCl). Nuclear extracts were prepared from 5–10 × 10⁶ cells by sequential lysis in Gough I buffer (with 0.65% NP-40) and high-salt buffer C (400 mM NaCl, 25% glycerol). Streptavidin-coated 96-well plates were loaded with 100 pmol biotinylated probes per well, blocked with 5% BSA, and incubated with 10–50 µg nuclear extract. Bound proteins were detected with primary antibodies (1:500, TEAD1 (Active Motif #61644), GATA4 (ActiveMotif #39894)) followed by HRP-conjugated secondary antibodies (0.01 µg/ml) (TransGen #HS101-01). Colorimetric detection was performed using OPD substrate (Thermo Fisher #34005) and absorbance measured at 492 nm.

### Luciferase reporter assays

Dual-luciferase reporter assays were performed using the pLenti-basP:fLuc-TK:rLuc lentiviral reporter vector (Addgene # 138369), which contains a firefly luciferase gene driven by a minimal promoter for enhancer activity measurement and a constitutively expressed Renilla luciferase cassette under the TK promoter as an internal control for lentiviral integration. The original Gateway cloning cassette was removed by double digestion with SbfI (NEB #R3642S) and ApaI (NEB #R0114S) and replaced with a unique BamHI (NEB #R3136S) restriction site to enable Gibson assembly–based cloning. Enhancer sequences (200 bp centered on variants of interest) were cloned into the BamHI-linearized vector by Gibson assembly (1:7 vector:insert ratio) and verified by Sanger sequencing. Lentiviral particles were produced in HEK293T cells and used to transduce HAVICs. Three days post-transduction, firefly and Renilla luciferase activities were measured using the Dual-Luciferase Reporter Assay System (Promega #E1910). Enhancer activity was expressed as the ratio of firefly to Renilla luminescence and compared between allelic constructs and a random sequence control.

### Scar-in-a-jar fibrosis assay

HAVICs were seeded in 12-well plates (15,000 cells/well) and cultured for 48 h^85^. Fibrosis was induced with assay medium containing 100 µM L-ascorbic acid 2-phosphate (Sigma #A8960), macromolecular crowding agents (37.5 mg/ml Ficoll 70 and 25 mg/ml Ficoll 400), and 5 ng/ml TGFβ1 (Gibco #PHG9214) for 6 days, refreshed every 2–3 days. Cells were fixed in ice-cold methanol (−20°C, 10 min), blocked with 5% BSA/0.05% Tween-20, and stained with anti-type I collagen (Sigma #C2456, 1:500) for 2 h at room temperature, followed by Alexa Fluor 647 secondary antibody (Invitrogen #A-21236, 1:1,000) and DAPI (1:1,000) overnight at 4°C. Images (4 × 4 grid per well) were acquired on a Zeiss Axio Observer Z1 (10× objective). Nuclei were counted using StarDist 2D, and collagen integrated density was normalized to nuclei count using ImageJ.

### Sequencing

This publication includes data generated at the UC San Diego IGM Genomics Center utilizing both Illumina NovaSeq 6000 and Illumina NovaSeq X Plus that were both purchased with funding from a National Institutes of Health SIG grant (#S10 OD026929).

## QUANTIFICATION AND STATISTICAL ANALYSIS

### Statistics

Statistical analyses were performed using R. Comparisons of continuous distributions between two groups were performed using two-sided Kolmogorov-Smirnov tests. Comparisons of continuous variables between two groups were performed using two-sided Wilcoxon rank-sum tests. Categorical comparisons between two groups (e.g., as-caQTLs versus balanced variants) were performed using two-sided Fisher’s exact tests. Gene set enrichment in predefined pathways was assessed using hypergeometric tests. Linear correlations were quantified using Pearson’s correlation coefficient, and Spearman’s rank correlation was used when relationships were non-linear or when variables were not normally distributed. Unless otherwise specified, p-values were adjusted for multiple testing using the Benjamini-Hochberg procedure.

### RNA-seq data processing

Paired-end reads were trimmed using fastp with default parameters. Reads were aligned to the reference genome (Genecode V35 lift 37) with STAR v2.7.9a and sorted with samtools. Gene-level quantification was performed with featureCounts (Subread) in paired-end mode with junction read counting and overlapping feature assignment.

### ATAC-seq data processing

Reads were trimmed with fastp and aligned to hg19 using Bowtie2. Peaks were called with MACS2 (--broad --broad-cutoff 0.01). A consensus peak set (326,594 regions after removing of Encode black list regions) was generated from merged BAM files. Signal tracks were produced with bamCoverage (deeptools), and fraction of reads in peak (FRiP) was calculated using bedtools multicov.

#### Transcription factor footprinting

Transcription factor footprinting analysis was performed using TOBIAS^22^. Duplicate reads were removed using Picard MarkDuplicates. Transcription factor binding for 755 position frequency matrices from the JASPAR 2024 CORE Homo sapiens collection were predicted.

### Allele-specific chromatin accessibility

Heterozygous SNVs within ATAC-seq peaks were identified using bcftools mpileup/call on hg19-aligned BAMs. Only biallelic autosomal SNVs with quality ≥20 were retained. Reference mapping bias was mitigated using the WASP pipeline^86^. Variants in ENCODE blacklist regions were excluded and remaining SNVs were annotated with dbSNV build 151. Allele-specific counts were quantified with GATK ASEReadCounter (v4.1.8.1; minimum 5 reads per allele with a minimum of 20 total reads). Background allelic dosage was estimated using BAbachi, retaining only diploid regions^13^. Allelic imbalance was tested using MixALime^11^ with a Marginalized Compound Negative Binomial. Significant SNVs (FDR < 0.10) were defined as allele-specific caQTLs and non significant SNVs (Pval > 0.10) were defined as balanced.

### Haplotype phasing and allele-specific signal tracks

Read-based haplotype phasing was performed using WhatsHap ^87^. Heterozygous variants from the allelic imbalance VCF files were phased using whatshap phase. Reads were assigned to parental haplotypes and haplotype-specific BAM files were generated. Haplotype-resolved signal tracks (bigWig) were produced from each haplotype BAM using bamCoverage (deeptools).

### ChIP-seq data processing

Reads were trimmed with fastp and aligned to hg19 using Bowtie2. Narrow peak calling was used for TEAD1 and CTCF; broad peak calling (--broad --broad-cutoff 0.01) for histone modifications. Chromatin states were annotated using ChromHMM^16^ with the 18-state expanded model based on six histone marks (H3K27ac, H3K27me3, H3K4me1, H3K4me3, H3K36me3, H3K9me3), following Roadmap Epigenomics Consortium recommendations^88^. Allelic imbalance in ChIP-seq data was assessed using the same pipeline as ATAC-seq, with a minimum of 20 overlapping reads and at least 3 reads per allele.

### HiChIP and Hi-C data processing

Raw reads from four HiChIP-H3K27ac samples and two Hi-C samples were trimmed with fastp and processed through HiC-Pro^89^ with default settings (MIN_MAPQ = 10, MboI restriction enzyme). Chromatin loops were called from merged HiChIP data using FitHiChIP^90^ in peak-to-all mode (5 kb bins, 20 kb-2 Mb range, FDR ≤ 0.01).

### Activity-By-Contact (ABC) model

Enhancer-gene connections were predicted using the ABC model^29^. Enhancer activity was estimated by combining ATAC-seq accessibility and H3K27ac signal. Three-dimensional contact frequency was derived from Hi-C matrices. Enhancer-gene pairs with an ABC score ≥0.025 were retained as high-confidence regulatory connections.

### csRNA-seq data processing

Reads were trimmed using HOMER to remove 3′ adapter sequences (maximum 2 mismatches, minimum match length 4). Reads were aligned to hg19 with STAR v2.7.9 using parameters optimized for short capped RNA fragments. Transcription start sites were identified using the HOMER findcsRNATSS.pl module, leveraging csRNA-seq enrichment relative to matched total RNA-seq and input background.

### Genotyping and eQTL analysis

Samples failing quality control (low call rate, sex mismatch, genotype discordance) were excluded. Variants were filtered by call rate (<0.97), MAF (<0.01), and HWE (P < 1E-07), yielding 454,278 variants. Imputation was performed using the TOPMed r3 reference panel (Rsq > 0.3). cis-eQTL mapping was performed for SNVs within ±100 kb of TSS (MAF > 0.1) using QTLtools v1.3.1, with sex, disease status, three expression PCs, and one genotype PC as covariates. Significance was assessed at FDR < 10%. Directionality of effects was compared with bulk aortic valve eQTLs (n = 484) using Z-tests.

### Transcription factor motif disruption analysis

The impact of genetic variants on TF binding motifs was assessed using PerfectosAPE^13^. For each variant, a 50 bp sequence centered on the SNV was extracted from hg19. Motif models were obtained from the JASPAR 2024 CORE Homo sapiens collection (755 matrices). Allele-specific changes in binding affinity were quantified as fold changes between reference and alternative alleles with PERFECTOS-APE.

### Stratified LD score regression (S-LDSC)

Partitioned heritability enrichment analysis was performed using stratified LD score regression (S-LDSC)^91^. Binary annotations derived from HAVIC as-caQTL data (±500bp) were generated for each autosome by assigning a value of 1 to HapMap3 SNVs (from the baselineLD v2.2 template) overlapping annotation intervals using bedtools map. Per-chromosome LD scores were computed with ldsc.py --l2 using the 1000 Genomes Phase 3 European reference panel, a 1 cM LD window, and HapMap3 SNVs as the SNV set. GWAS summary statistics were harmonized with munge_sumstats.py against the HapMap3 reference. Partitioned heritability was estimated for each GWAS trait using ldsc.py --h2 with the custom LD scores, HapMap3 regression weights excluding the HLA region, 1000 Genomes Phase 3 European allele frequencies, and the --overlap-annot flag to control for the baselineLD v2.2 annotations. Heritability enrichment, standard errors, and p-values were extracted for each annotation-GWAS pair.

### Phenome-wide association scans

Phenome-wide association scans for candidate variants were performed using the Cardiovascular Disease Knowledge Portal. For each variant, cross-trait associations were retrieved from the variant-level summary pages, and significant associations were reported.

### SuRE data

SuRE (Survey of Regulatory Elements) allele-specific regulatory activity data from HepG2 cells were obtained from van Arensbergen et al. (2019)^92^. Allele-specific regulatory effects were extracted for candidate variants.

### ADASTRA allele-specific TF binding

Allele-specific transcription factor binding (TF-ASB) calls were retrieved from the ADASTRA database^13^. Variants with FDR < 0.05 were considered significant TF-ASB variants.

### UDACHA and QTLbase

Allele-specific chromatin accessibility calls were retrieved from UDACHA^11^ by merging ATAC-seq, DNase-seq, and FAIRE-seq data; variants with FDR < 0.1 were retained. Cross-tissue molecular QTLs were retrieved from QTLbase^15^ using the variants reported in the resource. Additional caQTLs from Wenz et al. (2026)^93^ were included, retaining variants at FDR < 0.05.

### Whole-genome sequencing data

Summary statistics from the most recent UK Biobank whole-genome sequencing association study^10^ were obtained and lifted over from GRCh38 to GRCh37/hg19 using the UCSC liftOver tool. Variants were filtered on minor allele frequency (MAF < 0.01), annotated with the Ensembl Variant Effect Predictor (VEP) to exclude exonic SNVs, and intersected with the HAVIC ATAC-seq consensus peak set using bedtools intersect to retain variants located in open chromatin regions.

### Mendelian randomization

Two-sample Mendelian randomization (MR) analyses were performed using the LaScaMolMR Julia package^94^. Instrumental variables (IVs) were selected from exposure GWAS summary statistics at a genome-wide significance threshold of P < 5E-08. IVs were clumped for linkage disequilibrium using 1000 Genomes Phase 3 European reference genotypes at an r² threshold of 0.01. Causal effects were estimated using three complementary methods: inverse-variance weighted (IVW), MR-Egger, and weighted median. MR-Egger intercept was used to assess the presence of pleiotropy.

### Locus plot

Multi-track locus plots were generated using pyGenomeTracks^95^, a custom LocusZoom-style association plots displaying GWAS -log₁₀(p-values) with linkage disequilibrium-based coloring relative to the lead variant was added.

### Tag density and signal enrichment plots

Tag density profiles at genomic features of interest were generated using HOMER annotatePeaks.pl with the -hist option to compute average signal density centered on as-caQTLs or TSSs. The density of eQTLs and footprints in the vicinity of allele-specific chromatin accessibility QTLs (as-caQTLs) were computed using the GenomicRanges R package^96^ and visualized using ggplot2.

### Microarray data processing

Publicly available microarray expression data from human fetal heart (GSE75985) were processed using the NetworkAnalyst platform^97^. Raw data were background-corrected, log₂-transformed, and VSN-normalized. Probe-to-gene mapping was performed using the platform annotation file, and probes mapping to multiple genes were removed. Differential expression analysis between OFT and the four heart chambers was performed using the limma framework, and genes with an adjusted p-value < 0.05 were considered differentially expressed.

### Predictive model for chromatin accessibility

The Enformer^38^ deep learning model (transformer-based, 196,608 bp input, 128 bp resolution, 896 output bins) was fine-tuned on HAVIC ATAC-seq data using the gReLU^98^ framework with a custom lazy-loading function. ATAC-seq peaks from 16 merged samples served as training intervals, split by chromosome (chr10: validation; chr11: test). Fine-tuning proceeded in two phases: (1) head-only training (Adam, lr = 1 × 10⁻³, Poisson NLL loss) with early stopping (patience = 3), then (2) end-to-end training of all 11 transformer layers and head (Adam, lr = 5 × 10⁻⁵, gradient clipping max norm = 1.0) on a single NVIDIA RTX 5090 GPU. Performance was evaluated by Pearson and Spearman correlations on chr11. Variant effects were quantified as log₂ fold change between alternative and reference predictions at the three central bins; variants with |log₂FC| > 0.24 (95% specificity) were considered significant.

### Perturb-seq analysis

Libraries were processed with Cell Ranger v7.0.0 (GRCh38-2024-A reference). Differential expression analysis was performed using SCEPTRE^43^ with high-MOI settings, left side and union gRNA assignment. Discovery pairs were constructed in cis (±1 Mb); positive controls targeted known gene TSS. Quality filters included ≤5% mitochondrial reads, ≥15 non-zero cells per group, and UMI counts within the 5^th^-95^th^ percentile range. Trans-regulatory effects were assessed genome-wide for each perturbation target independently.

### Gene co-expression network analysis

Weighted gene co-expression network analysis (WGCNA)^99^ was performed on single-cell data normalized by library size and denoised with MAGIC^100^. Biweight midcorrelation was computed between gene pairs identified from trans-regulatory analyses, and programs were defined by hierarchical clustering (average linkage).

### Transcription factor enrichment analysis

TF binding enrichment was assessed using a distance-weighted approach adapted from ChEA3^101^. ChIP-seq peaks from ChIP-Atlas were used to compute binding scores for protein-coding genes, defined as the sum of distance to TSS-weighted peak scores (1 - |d|/50,000) within 50 kb of each TSS. Target genes were the top 5% of scored genes (maximum 1,500).

### Genetically Informed Perturbation Program Score (GIPPS)

For each co-expressed ciGene program, gene-level loss-of-function (LOF) burden effect sizes were obtained from UK Biobank whole-exome sequencing (n = 394,841; ICD code I35)^59^. Effect sizes were transformed to gene dosage (βdosage = -βLOF) and harmonized with CRISPRi perturbation direction. The program effect (β_program_) was defined as the mean of harmonized β_dosage_ values. Significance was assessed by permutation testing (10,000 iterations, sampling without replacement from background genes), yielding an empirical two-sided P-value. Results are reported as GIPPS (signed -log₁₀ P-value); |GIPPS| > 1.30 corresponds to P < 0.05. Confidence intervals for βprogram were derived from 10,000 within-program bootstrap resamples, reported as 2.5^th^-97.5^th^ percentiles.

### Single-cell RNA-seq analysis

Raw sequencing reads were aligned and quantified using Cell Ranger v7.0.0 (10x Genomics) with the GRCh38-2020-A reference transcriptome. Downstream analyses were performed in R using Seurat v5.3.1. Low-quality cells were filtered based on the following criteria: nFeature_RNA > 200 and < 14,000, percentage of mitochondrial reads < 5%, percentage of ribosomal protein S (RPS) reads < 15%, and percentage of ribosomal protein L (RPL) reads < 15%. Gene expression was normalized using NormalizeData, and highly variable features were identified with FindVariableFeatures. Data were scaled using ScaleData, and principal component analysis was performed with RunPCA (n = 100 components). Batch effects across samples were corrected using IntegrateLayers with the CCAIntegration method on log-normalized data.

Single-cell trajectories were inferred by first constructing a diffusion map from the integrated embedding using the DiffusionMap function from the destiny R package. Pseudotime and lineage structures were then computed on the diffusion map coordinates using slingshot.

## Author contributions

M.B. and P.M. conceived the project and designed the experiments. M.B, A.R. and V.D. performed HAVICs and HAVECs isolation and culture from aortic valve. A.R. performed HiChIP H3K27ac, HiC, ATAC-seq and genotyping experiments. A.R. and V.D. performed csRNA-seq experiments. A.R. and M.B performed ChIP-seq and perturb-seq experiments. D.K.B., N.G and V.S.A. performed HAVICS to myofibroblast single cell experiment. M.B. and P.M. performed data analysis with support from S.M. and T.K. All authors reviewed the manuscript and provided significant intellectual input. M.B. and P.M. drafted the manuscript.

## Data availability

All data generated in this study are included in the manuscript and its Supplementary Information. Sequencing data will be deposited in the NCBI Gene Expression Omnibus upon acceptance. The fine-tuned Enformer model for HAVIC ATAC-seq with code for model fine-tuning, and as-caQTLs discovery script, are available at Zenodo (accession: 10.5281/zenodo.20396493).

## References

1. Schäfers, H.-J. & Konstantinov, I. E. Surgical anatomy of aortic root: Toward precise and durable aortic, neo-aortic, and truncal valve repairs. J. Thorac. Cardiovasc. Surg. 169, 1287–1295 (2025).

2. Calcific aortic stenosis. Nat. Rev. Dis. Primer 2, 16007 (2016).

3. Moncla, L.-H. M., Briend, M., Bossé, Y. & Mathieu, P. Calcific aortic valve disease: mechanisms, prevention and treatment. Nat. Rev. Cardiol. 20, 546–559 (2023).

4. Marom, G. et al. Numerical model of the aortic root and valve: optimization of graft size and sinotubular junction to annulus ratio. J. Thorac. Cardiovasc. Surg. 146, 1227–1231 (2013).

5. Hayashi, H. et al. Influence of aneurysmal aortic root geometry on mechanical stress to the aortic valve leaflet. Eur. Heart J. Cardiovasc. Imaging 22, 986–994 (2021).

6. Small, A. M. et al. Genomic and transcriptomic analyses of aortic stenosis enhance therapeutic target discovery and disease prediction. Nat. Genet. 58, 57–66 (2026).

7. Gomes, B. et al. Genetic architecture of cardiac dynamic flow volumes. Nat. Genet. 8. 56, 245–257 (2024).

8. Pirruccello, J. P. et al. Deep learning enables genetic analysis of the human thoracic aorta. Nat. Genet. 54, 40–51 (2022).

9. Farh, K. K.-H. et al. Genetic and epigenetic fine mapping of causal autoimmune disease variants. Nature 518, 337–343 (2015).

10. Carss, K. et al. Whole-genome sequencing of 490,640 UK Biobank participants. Nature 645, 692–701 (2025).

11. Buyan, A. et al. Statistical framework for calling allelic imbalance in high-throughput sequencing data. Nat. Commun. 16, 1739 (2025).

12. Liang, Y., Aguet, F., Barbeira, A. N., Ardlie, K. & Im, H. K. A scalable unified framework of total and allele-specific counts for cis-QTL, fine-mapping, and prediction. Nat. Commun. 12, 1424 (2021).

13. Abramov, S. et al. Landscape of allele-specific transcription factor binding in the human genome. Nat. Commun. 12, 2751 (2021).

14. Wenz, B. M. et al. Genotype inference from aggregated chromatin accessibility data reveals genetic regulatory mechanisms. Genome Biol. 26, 81 (2025).

15. Huang, D. et al. QTLbase2: an enhanced catalog of human quantitative trait loci on extensive molecular phenotypes. Nucleic Acids Res. 51, D1122–D1128 (2023).

16. Ernst, J. & Kellis, M. Chromatin-state discovery and genome annotation with ChromHMM. Nat. Protoc. 12, 2478–2492 (2017).

17. Jonsson, M. K. B. et al. A Transcriptomic and Epigenomic Comparison of Fetal and Adult Human Cardiac Fibroblasts Reveals Novel Key Transcription Factors in Adult Cardiac Fibroblasts. JACC Basic Transl. Sci. 1, 590–602 (2016).

18. Hu, Y. et al. DisGeNet: a disease-centric interaction database among diseases and various associated genes. Database J. Biol. Databases Curation 2025, baae122 (2025).

19. Martin, A. R. et al. PanelApp crowdsources expert knowledge to establish consensus diagnostic gene panels. Nat. Genet. 51, 1560–1565 (2019).

20. Holmes, K. W. et al. GenTAC registry report: gender differences among individuals with genetically triggered thoracic aortic aneurysm and dissection. Am. J. Med. Genet. A. **161A**, 779–786 (2013).

21. Cooper, G. M. et al. Distribution and intensity of constraint in mammalian genomic sequence. Genome Res. 15, 901–913 (2005).

22. Bentsen, M. et al. ATAC-seq footprinting unravels kinetics of transcription factor binding during zygotic genome activation. Nat. Commun. 11, 4267 (2020).

23. Afouda, B. A. Towards Understanding the Gene-Specific Roles of GATA Factors in Heart Development: Does GATA4 Lead the Way? Int. J. Mol. Sci. 23, 5255 (2022).

24. Currey, L., Thor, S. & Piper, M. TEAD family transcription factors in development and disease. Development 148, dev196675 (2021).

25. Santoyo-Suarez, M. G. et al. The Involvement of Krüppel-like Factors in Cardiovascular Diseases. Life 13, 420 (2023).

26. Windak, R. et al. The AP-1 Transcription Factor c-Jun Prevents Stress-Imposed Maladaptive Remodeling of the Heart. PLOS ONE 8, e73294 (2013).

27. Berest, I. et al. Quantification of Differential Transcription Factor Activity and Multiomics-Based Classification into Activators and Repressors: diffTF. Cell Rep. 29, 3147–3159.e12 (2019).

28. Klemm, S. L., Shipony, Z. & Greenleaf, W. J. Chromatin accessibility and the regulatory epigenome. Nat. Rev. Genet. 20, 207–220 (2019).

29. Fulco, C. P. et al. Activity-by-contact model of enhancer–promoter regulation from thousands of CRISPR perturbations. Nat. Genet. 51, 1664–1669 (2019).

30. Briend, M. et al. Connectome and regulatory hubs of CAGE highly active enhancers. Sci. Rep. 13, 5594 (2023).

31. Duttke, S. H., Chang, M. W., Heinz, S. & Benner, C. Identification and dynamic quantification of regulatory elements using total RNA. Genome Res. 29, 1836–1846 (2019).

32. Duttke, S. H. et al. Position-dependent function of human sequence-specific transcription factors. Nature 631, 891–898 (2024).

33. van Arensbergen, J. et al. High-throughput identification of human SNPs affecting regulatory element activity. Nat. Genet. 51, 1160–1169 (2019).

34. Thériault, S. et al. Integrative genomic analyses identify candidate causal genes for calcific aortic valve stenosis involving tissue-specific regulation. Nat. Commun. 15, 2407 (2024).

35. Arthur, T. D. et al. Multiomic QTL mapping reveals phenotypic complexity of GWAS loci and prioritizes putative causal variants. Cell Genomics 5, 100775 (2025).

36. Jeong, R. & Bulyk, M. L. Chromatin accessibility variation provides insights into missing regulation underlying immune-mediated diseases. eLife 13, RP98289 (2025).

37. Gazal, S. et al. Linkage disequilibrium–dependent architecture of human complex traits shows action of negative selection. Nat. Genet. 49, 1421–1427 (2017).

38. Avsec, Ž., et al. Effective gene expression prediction from sequence by integrating long-range interactions. Nat. Methods 18, 1196–1203 (2021).

39. Kodo, K. et al. GATA6 mutations cause human cardiac outflow tract defects by disrupting semaphorin-plexin signaling. Proc. Natl. Acad. Sci. U. S. A. 106, 13933–13938 (2009).

40. Chakraborty, S. et al. Twist1 promotes heart valve cell proliferation and extracellular matrix gene expression during development in vivo and is expressed in human diseased aortic valves. Dev. Biol. 347, 167–179 (2010).

41. Morris, J. A. et al. Discovery of target genes and pathways at GWAS loci by pooled single-cell CRISPR screens. Science 380, eadh7699 (2023).

42. Gasperini, M. et al. A Genome-wide Framework for Mapping Gene Regulation via Cellular Genetic Screens. Cell 176, 377–390.e19 (2019).

43. Barry, T., Wang, X., Morris, J. A., Roeder, K. & Katsevich, E. SCEPTRE improves calibration and sensitivity in single-cell CRISPR screen analysis. Genome Biol. 22, 344 (2021).

44. Chignon, A. et al. Enhancer-associated aortic valve stenosis risk locus 1p21.2 alters NFATC2 binding site and promotes fibrogenesis. iScience 24, 102241 (2021).

45. Thériault, S. et al. Genetic Association Analyses Highlight IL6, ALPL, and NAV1 As 3 New Susceptibility Genes Underlying Calcific Aortic Valve Stenosis. Circ. Genomic Precis. Med. 12, e002617 (2019).

46. Zaharija, B., Samardžija, B. & Bradshaw, N. J. The TRIOBP Isoforms and Their Distinct Roles in Actin Stabilization, Deafness, Mental Illness, and Cancer. Molecules 25, 4967 (2020).

47. Jover, E. et al. Sex-dependent expression of galectin-1, a cardioprotective β-galactoside-binding lectin, in human calcific aortic stenosis. FASEB J. Off. Publ. Fed. Am. Soc. Exp. Biol. 38, e23447 (2024).

48. Farahani, R. M. & Xaymardan, M. Platelet-Derived Growth Factor Receptor Alpha as a Marker of Mesenchymal Stem Cells in Development and Stem Cell Biology. Stem Cells Int. 2015, 362753 (2015).

49. Chandrakanthan, V. et al. Mesoderm-derived PDGFRA+ cells regulate the emergence of hematopoietic stem cells in the dorsal aorta. Nat. Cell Biol. 24, 1211–1225 (2022).

50. Lucken-Ardjomande Häsler, S., Vallis, Y., Pasche, M. & McMahon, H. T. GRAF2, WDR44, and MICAL1 mediate Rab8/10/11-dependent export of E-cadherin, MMP14, and CFTR ΔF508. J. Cell Biol. 219, e201811014 (2020).

51. Kanekura, K., Nishimoto, I., Aiso, S. & Matsuoka, M. Characterization of amyotrophic lateral sclerosis-linked P56S mutation of vesicle-associated membrane protein-associated protein B (VAPB/ALS8). J. Biol. Chem. 281, 30223–30233 (2006).

52. Appenzeller-Herzog, C. & Hauri, H.-P. The ER-Golgi intermediate compartment (ERGIC): in search of its identity and function. J. Cell Sci. 119, 2173–2183 (2006).

53. Yang, B. et al. TRIM35 triggers cardiac remodeling by regulating SLC7A5-mediated amino acid transport and mTORC1 activation in fibroblasts. Cell Commun. Signal. CCS 22, 444 (2024).

54. Hagenbeek, T. J. et al. An allosteric pan-TEAD inhibitor blocks oncogenic YAP/TAZ signaling and overcomes KRAS G12C inhibitor resistance. *Nat*. Cancer 4, 812–828 (2023).

55. Cao, J. et al. A human cell atlas of fetal gene expression. Science 370, eaba7721 (2020).

56. Soh, B.-S. et al. Endothelin-1 supports clonal derivation and expansion of cardiovascular progenitors derived from human embryonic stem cells. Nat. Commun. 7, 10774 (2016).

57. Gabriel, L. A. R. et al. ADAMTSL4, a secreted glycoprotein widely distributed in the eye, binds fibrillin-1 microfibrils and accelerates microfibril biogenesis. Invest. Ophthalmol. Vis. Sci. 53, 461–469 (2012).

58. Collins, R. L. et al. A cross-disorder dosage sensitivity map of the human genome. Cell 185, 3041–3055.e25 (2022).

59. Karczewski, K. J. et al. Systematic single-variant and gene-based association testing of thousands of phenotypes in 394,841 UK Biobank exomes. Cell Genomics 2, 100168 (2022).

60. Dietz, H. C. et al. Marfan syndrome caused by a recurrent de novo missense mutation in the fibrillin gene. Nature 352, 337–339 (1991).

61. Ren, P. et al. Heterogeneous Cardiac-Derived and Neural Crest-Derived Aortic Smooth Muscle Cells Exhibit Similar Transcriptional Changes After TGFβ Signaling Disruption. Arterioscler. Thromb. Vasc. Biol. 45, 260–276 (2025).

62. Olivieri, J., Smaldone, S. & Ramirez, F. Fibrillin assemblies: extracellular determinants of tissue formation and fibrosis. Fibrogenesis Tissue Repair 3, 24 (2010).

63. Carta, L. et al. Fibrillins 1 and 2 perform partially overlapping functions during aortic development. J. Biol. Chem. 281, 8016–8023 (2006).

64. Zhou, D., Zhou, Y., Xu, Y., Meng, R. & Gamazon, E. R. A phenome-wide scan reveals convergence of common and rare variant associations. Genome Med. 15, 101 (2023).

65. Wright, S. N., Yang, J. & Ideker, T. Common and rare genetic variants show network convergence for a majority of human traits. EMBO Rep. 10.1038/s44319-026-00733-4 (2026).

66. Ota, M. et al. Causal modelling of gene effects from regulators to programs to traits. Nature 650, 399–408 (2026).

67. Lee, I. H. et al. Ahnak functions as a tumor suppressor via modulation of TGFβ/Smad signaling pathway. Oncogene 33, 4675–4684 (2014).

68. Zou, Z., Ohta, T. & Oki, S. ChIP-Atlas 3.0: a data-mining suite to explore chromosome architecture together with large-scale regulome data. Nucleic Acids Res. 52, W45–W53 (2024).

69. Eletto, D., Eletto, D., Dersh, D., Gidalevitz, T. & Argon, Y. Protein disulfide isomerase A6 controls the decay of IRE1α signaling via disulfide-dependent association. Mol. Cell 53, 562–576 (2014).

70. Kaneko, M. et al. Genome-wide identification and gene expression profiling of ubiquitin ligases for endoplasmic reticulum protein degradation. Sci. Rep. 6, 30955 (2016).

71. Laguna-Fernandez, A. et al. Iron alters valvular interstitial cell function and is associated with calcification in aortic stenosis. Eur. Heart J. 37, 3532–3535 (2016).

72. Grishin, D. & Gusev, A. Allelic imbalance of chromatin accessibility in cancer identifies candidate causal risk variants and their mechanisms. Nat. Genet. 54, 837–849 (2022).

73. Hinton, R. B. et al. Elastin haploinsufficiency results in progressive aortic valve malformation and latent valve disease in a mouse model. Circ. Res. 107, 549–557 (2010).

74. Morvan, M. et al. Relationship of Iron Deposition to Calcium Deposition in Human Aortic Valve Leaflets. J. Am. Coll. Cardiol. 73, 1043–1054 (2019).

75. Boyle, E. A., Li, Y. I. & Pritchard, J. K. An Expanded View of Complex Traits: From Polygenic to Omnigenic. Cell 169, 1177–1186 (2017).

76. Brunton, H., Garner, I. M., Bailey, U.-M., Upstill-Goddard, R. & Bailey, P. J. Using Chromatin Accessibility to Delineate Therapeutic Subtypes in Pancreatic Cancer Patient-Derived Cell Lines. STAR Protoc. 1, 100079 (2020).

77. Arrigoni, L. et al. RELACS nuclei barcoding enables high-throughput ChIP-seq. *Commun*. Biol. 1, 214 (2018).

79. Lafontaine, D. L., Yang, L., Dekker, J. & Gibcus, J. H. Hi-C 3.0: Improved Protocol for Genome-Wide Chromosome Conformation Capture. Curr. Protoc. 1, e198 (2021).

79. Gryder, B. E. et al. Miswired Enhancer Logic Drives a Cancer of the Muscle Lineage. iScience 23, 101103 (2020).

80. Mumbach, M. R. et al. HiChIP: efficient and sensitive analysis of protein-directed genome architecture. Nat. Methods 13, 919–922 (2016).

81. Meyer, M. K. et al. Profiling active RNA polymerase II transcription start sites from total RNA by capped small RNA sequencing (csRNA-seq). Nat. Protoc. 1–25 (2026) doi:10.1038/s41596-025-01285-y.

82. Datlinger, P. et al. Pooled CRISPR screening with single-cell transcriptome readout. Nat. Methods 14, 297–301 (2017).

83. McKenna, A. & Shendure, J. FlashFry: a fast and flexible tool for large-scale CRISPR target design. BMC Biol. 16, 74 (2018).

84. Brand, L. H., Kirchler, T., Hummel, S., Chaban, C. & Wanke, D. DPI-ELISA: a fast and versatile method to specify the binding of plant transcription factors to DNA in vitro. Plant Methods 6, 25 (2010).

85. Stebler, S. & Raghunath, M. The Scar-in-a-Jar: In Vitro Fibrosis Model for Anti-Fibrotic Drug Testing. Methods Mol. Biol. 2299, 147–156 (2021).

86. van de Geijn, B., McVicker, G., Gilad, Y. & Pritchard, J. K. WASP: allele-specific software for robust molecular quantitative trait locus discovery. Nat. Methods 12, 1061–1063 (2015).

87. Martin, M. et al. WhatsHap: fast and accurate read-based phasing. 085050 Preprint at 10.1101/085050 (2016).

88. Kundaje, A. et al. Integrative analysis of 111 reference human epigenomes. Nature 518, 317–330 (2015).

89. Servant, N. et al. HiC-Pro: an optimized and flexible pipeline for Hi-C data processing. Genome Biol. 16, 259 (2015).

90. Bhattacharyya, S., Chandra, V., Vijayanand, P. & Ay, F. Identification of significant chromatin contacts from HiChIP data by FitHiChIP. Nat. Commun. 10, 4221 (2019).

91. Finucane, H. K. et al. Partitioning heritability by functional annotation using genome-wide association summary statistics. Nat. Genet. 47, 1228–1235 (2015).

92. van Arensbergen, J. et al. High-throughput identification of human SNPs affecting regulatory element activity. Nat. Genet. 51, 1160–1169 (2019).

93. Wenz, B. M. et al. Expanded chromatin accessibility mapping explains genetic variation associated with complex traits in liver. Am. J. Hum. Genet. 113, 260–275 (2026).

94. Mathieu, S. et al. Efficient molecular mendelian randomization screens with LaScaMolMR.jl. 2024.08.29.24312805 Preprint at 10.1101/2024.08.29.24312805 (2024).

95. Lopez-Delisle, L. et al. pyGenomeTracks: reproducible plots for multivariate genomic datasets. Bioinformatics 37, 422–423 (2021).

96. Lawrence, M. et al. Software for Computing and Annotating Genomic Ranges. PLOS Comput. Biol. 9, e1003118 (2013).

97. Zhou, G. et al. NetworkAnalyst 3.0: a visual analytics platform for comprehensive gene expression profiling and meta-analysis. Nucleic Acids Res. 47, W234–W241 (2019).

98. Lal, A., Gunsalus, L., Nair, S., Biancalani, T. & Eraslan, G. gReLU: a comprehensive framework for DNA sequence modeling and design. Nat. Methods 22, 2253–2257 (2025).

99. Langfelder, P. & Horvath, S. WGCNA: an R package for weighted correlation network analysis. BMC Bioinformatics 9, 559 (2008).

100. van Dijk, D. et al. Recovering Gene Interactions from Single-Cell Data Using Data Diffusion. Cell 174, 716–729.e27 (2018).

101. Keenan, A. B. et al. ChEA3: transcription factor enrichment analysis by orthogonal omics integration. Nucleic Acids Res. 47, W212–W224 (2019).

