## Supplementary_materials for "Decoding causal genes and programs from regulatory variants in aortic valve disease"

­­**EXPERIMENTAL MODEL DETAILS**

***Cell lines and culture***

HEK293T cells were obtained from American Type Culture Collection (ATCC #CRL-11268). Immortalized Human Bone Marrow Mesenchymal Stromal Cells - hTERT (iMSC3) cells were obtained from Applied Biological Materials Inc. (ABM #T0529).

HEK293T, iMSC, mVICs and HAVICs were cultured in Dulbecco’s Modified Eagle Medium (DMEM) high glucose, pyruvate (Gibco™#11995-065) supplemented with 10 % fetal bovine serum (VWR# 97068-085), 1 % Penicillin-Streptomycin (Gibco™#15240-062), 0.2% mycozap (Lonza #VZA-2032), 10mM sodium pyruvate (Gibco™#11360-070), 20mM L-glutamin (Gibco™#25030-081). Cell passage was performed using a 1:5 dilution of trypsin-EDTA (0.5%) without phenol red (Gibco™#15400-054). HAVECs were cultured in endothelial cell growth medium (ATCC #PCS-100-030) supplemented with a BBE kit (ATCC #PCS-100-040) and passed with a 1:10 dilution of the same trypsin-EDTA.

Human aortic valve interstitial cells (HAVICs) and endothelial cells (HAVECs) were isolated from non-mineralized aortic valves obtained from patients who underwent cardiac surgery at the *Institut universitaire de cardiologie et de pneumologie de Québec – Université Laval* following heart transplantation without aortic valve abnormalities, or from mineralized aortic valves obtained during valve replacement. All patients provided informed consent, and the study was approved by the IUCPQ ethics committee. Aortic leaflets were pre-wash in 10ml HBSS 1X (Gibco #14175-095) and incubated at 37°C and 5% CO2 with rotation for 10 min in 5ml of 2.6mg/ml type II collagenase (Gibco™#171010-015) dilute in complete DMEM. Leaflets were then rubbed with cotton stem to untie HAVECS, centrifuged for 5 min at 200 rcf and resuspend in endothelial cell growth BBE medium (ATCC). Aortic leaflets were then cut into small pieces and incubated at 37°C and 5% CO2 with rotation for 45 min in 5ml of 3mg/ml type I collagenase (Gibco™#17100-017) dilute in HBSS 1X. Digest tissues are filtered through a 70 µm mesh, centrifuged for 5 min at 200 rcf and resuspended in DMEM complete medium. All HAVICs and HAVECs were used between passages 3 to 7.

Mouse aortic valve interstitial cells (mAVIC) were isolated from aortic valves obtained by microdissection. Pooled aortic valves from 5 to 10 C57BL/6 mice were washed in cold 1X HEPES and centrifuged for 5 minutes at 200 rcf. These tissues were then incubated for 30 minutes at 37°C with agitation in a solution of type I collagenase (Gibco™#17100-017) solubilized at 1 mg/ml in a 1:1 mixture of serum-free DMEM and HEPES (+pen/strep). The tissues were washed twice in cold HEPES, centrifuged, and then incubated with agitation for 30 minutes at 37°C and 5% CO2 in a solution of type I collagenase solubilized at 4.5 mg/mL in a 1:1 mixture of serum-free DMEM and HEPES (+pen/strep). The tissues are collected by centrifugation for 5 minutes at 200 rcf and washed three times with complete DMEM medium containing 10% fetal bovine serum (FBS). The pellet is then resuspended in 500µl of DMEM medium containing 10% FBS and the mixture is transferred to two wells of a 24-well plate, ensuring that the volume of medium in the wells is as low as possible, just enough to cover the surface to increase the efficiency of adhesion. More than two wells may be used if necessary. The plate was then incubated undisturbed at 37°C and 5% CO₂ for several days. To prevent the wells from drying out, medium was added slowly when necessary. The medium was changed after one week.

***Lentivirus preparation***

Lentiviral particles were produced in HEK 293T cells (passage < 20). Cells were seeded at 7 × 10⁶ cells per 10 cm dish in DMEM supplemented with 10% FBS and incubated overnight. The following morning, medium was replaced with lentivirus packaging medium [Opti-MEM (Gibco 31985-070) supplemented with 5% FBS and 200 µM sodium pyruvate]. Transfection was performed using Lipofectamine 3000 Transfection Reagent (Thermo Fisher Scientific # L3000015) according to a modified protocol. For each 10 cm dish, a first mix (mix A) containing 12 µg of the transfer plasmid, 9 µg of psPAX2 (packaging plasmid Addgene 12260), 3 µg of pMD2.G (envelope plasmid Addgene 12259), and 36 µl of P3000 reagent was prepared in 1.5 ml Opti-MEM I. A second mix (mix B) containing 42 µl of Lipofectamine 3000 in 1.5 ml Opti-MEM I was prepared separately. The two mixes were combined, incubated for 20–30 minutes at room temperature, and added dropwise to the cells. After 4 hours, medium was replaced with fresh lentivirus packaging medium and 1/500 ViralBoost Reagent (Alstem #VB100) was added. Viral supernatants were collected at 24 and 48 hours post-transfection, pooled, filtered through a 0.45 µm syringe filter and concentrated with Lenti-X concentrator (Takara #631231) following recommendation. Lentiviral aliquots were either used immediately or snap-frozen and stored at −80°C. The target cells were seeded the day before transduction in their respective growth media. The following day, the medium was replaced with lentivirus packaging medium supplemented with 8 µg/mL polybrene (Sigma #H9268-5G), and then the appropriate volume of lentiviral supernatant was added. Twenty-four hours after transduction, the medium was changed, and 48 hours after transduction, antibiotic selection was initiated. Selection was maintained throughout the experiment, with medium changes every 2 to 3 days.

**METHOD DETAILS**

***RNAseq***

HAVICs were isolated from aortic valve leaflets obtained from 14 patients undergoing aortic valve replacement for CAVD and 13 control patients undergoing heart transplantation (10 women and 17 men), as described above. RNA was extracted from HAVIC cells cultured in 100 mm Petri dishes using the E.Z.N.A. Total RNA Kit I (Omega Bio-Tek, #R6834-02). RNA was also extracted from HAVEC cells obtained from four donors who had undergone heart transplantation (two women and two men). The quality of the RNA was assessed using a Bioanalyzer and the Agilent RNA 6000 Nano Kit (#5067-1511). All RNA used for libraries from HAVIC and HAVEC cells had a RIN ≥9.2. The Illumina® Stranded mRNA Prep Ligation Kit (cat. no. 20040532) was used for library preparation with 1 µg of starting material, as described in the protocol. mRNA libraries were accurately quantified using an Invitrogen Qubit 4 Fluorometer with 1X dsDNA High Sensitivity (#Q33230) and analysed using an Agilent 2100 Bioanalyzer with an Agilent DNA HS Kit (cat. no. 5067-4626). Libraries were then pooled in accordance with the recommendations of the UCSD-IGM Genomics Centre (University of California, San Diego, Institute for Genomic Medicine, Genomics Centre), and sequenced using a paired-end approach with a read length of 100 bp on there Illumina NovaSeq S4 platform.

***Genotyping***

DNA was extracted from buffy coat samples prepared by the Biobank of the *Institut Universitaire de Cardiologie et de Pneumologie de Québec (IUCPQ)* Research Centre and stored at −80°C using the QIAamp DNA Blood Midi Kit (Qiagen #). DNA concentrations were assessed using the Quant-iT PicoGreen dsDNA Kit (Invitrogen #P7589) and analysed using FilterMax F3 Multi-Mode Microplate Readers (Molecular Devices). The samples were diluted to 50 ng/μl in 96-well working plates and stored at −30 °C until genotyping, which was performed using Illumina Infinium™ Global Screening Array-24 v3.0 technology at the Genome Quebec Centre of Expertise and Services. All patients signed informed consent for genetic studies to be carried out. The study was approved by the ethics committee of the *Institut Universitaire de Cardiologie et de Pneumologie de Québec.*

***ATACseq library***

ATAC-seq libraries were prepared based on the [Kaestner Lab ATAC-seq Protocol](https://www.med.upenn.edu/kaestnerlab/assets/user-content/documents/ATAC-seq-Protocol-(Omni)-Kaestner-Lab.pdf) and STAR Protocol from Brunton H et al.^1^, with some modifications. In detail, freshly trypsinised HAVIC cells were isolated and counted from the leaflets of ten non-mineralised aortic valves from control subjects and six valves from CAVD patients. Aliquots of 40,000 cells were centrifuged at 200 rcf for five minutes and washed in 1 ml of cold PBS 1x. The cells were pelleted again at 4 °C, resuspended in 100 µl of cold lysis buffer (10 mM Tris-HCl pH 7.5, 10 mM NaCl, 3 mM MgCl2, 0.1% v/v NP-40, 0.1% v/v Tween-20, 0.01% v/v Digitonin) and incubated on ice for 10 minutes. Then, 1 ml of wash buffer (10 mM Tris-HCl pH 7.5, 10 mM NaCl, 3 mM MgCl2, 0.1% v/v Tween-20) was added, after which the nuclei were centrifuged at 1200 rcf. The nuclei were resuspended in 20 µl of the 1x transposition reaction mix (10 µl of 2x buffer and 10 µl of Tagment DNA Enzyme 1 Tn5 Transposase [Illumina kit #20034197]). The Tn5 reaction mix was incubated at 37 °C for 1–1.5 hours on a thermomixer at 1250 rpm. The transposed samples were then purified using a MicroElute GEL/PCR Purification Kit (GeneBio Systems, Inc., #FAEPK001-1BG) and eluted in 11 μl of H₂O. The purified samples were amplified and indexed in a 50 µl PCR mixture using the barcoded primers described in Table 1 of the STAR protocol by Brunton et al. (2020). The following PCR program is used : 72 °C for 5 min, hot start at 98 °C for 1 min, followed by 10 cycles of 98 °C for 15 sec, 63 °C for 30 sec and 72 °C for 1 min. The samples were then purified using a 1.8x SPRIselect bead ratio (Beckman, #B23317). The ATAC-seq libraries were accurately quantified using an Invitrogen Qubit 4 Fluorometer with 1X dsDNA High Sensitivity (#Q33230) and analysed using an Agilent 2100 Bioanalyzer with an Agilent DNA HS Kit (cat. no. 5067-4626). ATAC-seq libraries were pooled in accordance with the recommendations of the UCSD-IGM Genomics Centre (University of California, San Diego, Institute for Genomic Medicine, Genomics Centre), and sequenced using a paired-end approach with a read length of 150 bp on there Illumina NovaSeq X Plus 25B platform.

***ChIPseq***

ChIP-seq library was adapted from the RELACS protocol developed by [Arrigoni L et al.](https://pmc.ncbi.nlm.nih.gov/articles/PMC6281648/)^2^, as it provides reliable results from a small number of primary cells. Specifically, two million freshly trypsinised HAVIC cells from three biological replicates were fixed with a 1% formaldehyde solution (1% formaldehyde (Sigma 252549), 10% FBS in 1x PBS) for 10 minutes at room temperature (RT). The formaldehyde was then quenched with 125 mM glycine for a further 5 minutes at room temperature. The cells were centrifuged for five minutes at 200 rcf and washed with ice-cold 1x PBS. For nuclear extraction, the cells were resuspended in 500 µl of ice-cold lysis buffer (10 mM Tris-HCl, pH 8.0; 10 mM NaCl; 0.2% NP-40; 1x protease inhibitors) and incubated for 1 hour at 4 °C on a rotary wheel. The nuclei were pelleted at 2,200 rcf for five minutes at 4 °C, washed with 500 µl of cold lysis buffer, and pelleted again. The nuclei were then incubated with 100 µl of 0.5% SDS at 62 °C for 10 minutes. The SDS was quenched by adding 335 µl of 1.5% Triton X-100 and incubating for 15 minutes at 37 °C on a rotating wheel. The chromatin was then digested by adding 50 µl of 10x DPNII buffer, 8 µl of DPNII (NEB R0543M) and 7 µl of water, and incubating for 6 hours at 37 °C on a rotating wheel.

The nuclei were pelleted at 2,200 rcf for 5 minutes at 4 °C. The supernatant was removed, and the nuclei pellet was resuspended in 630 µL of shearing buffer (10 mM Tris-HCl, pH 8.0; 0.1% SDS; 1 mM EDTA). A 15 µL aliquot of the sample was set aside for the digestion control, and the remainder was split into two 0.5 mL tubes (BrandTech #781310), each containing 308 µL. Chromatin was released by sonication at 4 °C using a QSONICA Q800R3-110 for 11 minutes (30 seconds ON, 30 seconds OFF), with 30% amplification. Samples were clarified at maximum speed at 4 °C for 10 minutes, after which the two resulting supernatants were pooled. A 15 µl chromatin aliquot was set aside for the sonication control. The remaining 600 µl of sonicated chromatin was diluted in 1,200 µl of 1× Diagenode Buffer iC1 (Diagenode iDeal ChIP-seq Kit for Histones, #C01010173), to which 1× Protease Inhibitor Cocktail 100× (Sigma, #P8340) was added. For the histone mark ChIPs, the diluted chromatin was divided into six tubes, each containing 300 µl (5% of the volume was kept for input and stored at -20 °C). The antibodies of interest were then added: 3 µg of H3K27ac (Abcam, #ab4729), H3K4me3 (Abcam, #ab8580), H3K4me1 (CellSignaling #5326S), H3K27me3 (Diagenode, #C15410195), H3K36me3 (Diagenode, #C15410192) and H3K9me3 (Abcam, #ab8898). The tubes are incubated overnight at 4°C on a rotating wheel. For the ChIP of TEAD1 (three replicates) and CTCF (two replicates), 1,800 µl of diluted chromatin was used in full for one ChIP (5% of the volume was kept for INPUT and stored at -20°C), after which the antibodies of interest were added: 10 µl of TEAD1 (Active Motif #61644) and 3 µl of CTCF (Cell Signalling #2899). The tubes were then incubated overnight at 4 °C on a rotating wheel. Thirty microlitres of prewashed Dynabeads Protein G (Invitrogen™ #10004D) were added to each tube and incubated for three hours at 4 °C on a rotating wheel. After a brief period of centrifugation, the tubes were placed on a magnetic stand and the supernatant was removed. Then, 350 µl of ice-cold iW1 wash buffer was added to the beads, which were placed on a wheel and left to stand for five minutes at 4 °C. The wash buffer was then removed using the magnetic stand. This washing procedure was repeated with each of the different buffers (iW2, iW3 and iW4), using the same volume, as described above. After removing the final wash buffer, 100 µl of iE1 elution buffer was added to the beads, which were then incubated on a low-speed vortex for 30 minutes at room temperature. After a brief spin, the tubes were placed on the magnetic stand and the supernatant was transferred to a new tube. 85 µl of buffer iE1 were also added to the 15 µl input samples. Two microlitres of RNase A (Qiagen #19101) were added to all tubes, which were then incubated at 37 °C for one hour. Then, 4 µl of elution buffer iE2 and 2 µl of proteinase K (Qiagen #19131) were added, after which the tubes were incubated overnight at 65 °C (Thermomixer F1.5 Eppendorf). The ChIP DNA was then purified using the Zymo DNA Clean and Concentrator Kit (Zymo #D5205) and eluted in 53 µl of TE 1x. The purified DNA was accurately quantified using an Invitrogen Qubit 4 fluorometer with 1X dsDNA High Sensitivity (#Q33230) kit. For ChIP-seq library preparation, 50 µl of purified ChIP and input DNA was used. Libraries were prepared using the NEBNext Ultra II DNA Library Prep Kit for Illumina (E7645S, NEB), with some modifications. Following adapter ligation, size selection was omitted and the adapter-ligated fragments were purified using a 1× SPRIselect (Beckman Coulter #B23318) ratio. USER enzyme treatment was performed together with PCR enrichment using the following programme: Incubation at 37 °C for 15 min, hot start at 98 °C for 30 sec, followed by 12 cycles of 98 °C for 10 sec and 65 °C for 75 sec. A final extension was performed at 65 °C for 5 minutes. The amplified libraries were then purified using 1X SPRIselect beads. All libraries were accurately quantified using an Invitrogen Qubit 4 Fluorometer with 1X dsDNA High Sensitivity (#Q33230) and analysed using an Agilent 2100 Bioanalyzer with an Agilent DNA HS Kit (cat. no. 5067-4626). ChIP libraries were pooled in accordance with the recommendations of the UCSD-IGM Genomics Centre (University of California, San Diego, Institute for Genomic Medicine, Genomics Centre), and sequenced using a paired-end approach with a read length of 150 bp on there Illumina NovaSeq X Plus 25B platform.

The digestion and sonication controls (15 µl) were de-crosslinked with a final concentration of 1% SDS and 200 mM NaCl overnight at 65 °C. RNase A (Qiagen #19101) was added to 100 µl of TE buffer and the mixture was incubated for 1 h at 37 °C, followed by a proteinase K reaction (Qiagen #19131) and incubation for 1 h at 56 °C. DNA purification of the control sample was performed using the Zymo ChIP DNA Clean and Concentrator Kit (#D5205), and DNA quality control was performed according to the Agilent DNA HS Kit protocol (cat. no. 5067-4626).

***HiChIP***

HiChIP protocol was adapted from protocol of Mumbach MR et al.^3^ One million freshly trypsinised HAVIC cells from four biological replicates were fixed with a 1% formaldehyde solution (1% formaldehyde, 10% FBS in 1x PBS) for 10 minutes at room temperature (RT). The formaldehyde was then quenched with 125 mM glycine for a further 5 minutes at room temperature. The cells were centrifuged for five minutes at 200 rcf and washed with ice-cold 1x PBS. For nuclear extraction, the cells were resuspended in 500 µl of ice-cold lysis buffer (10 mM Tris-HCl, pH 8.0; 10 mM NaCl; 0.2% NP-40; 1x protease inhibitors) and incubated for 1 hour at 4 °C on a rotary wheel. The nuclei were pelleted at 2,200 rcf for five minutes at 4 °C, washed with 500 µl of cold lysis buffer, and pelleted again. The nuclei were then incubated with 100 µl of 0.5% SDS at 62 °C for 10 minutes. The SDS was quenched by adding 335 µl of 1.5% Triton X-100 and incubating for 15 minutes at 37 °C on a rotating wheel. The chromatin was then digested by adding 50 µl of 10x NEB buffer #2, 8 µl of MboI (NEB R0147) and incubating for 2 hours at 37 °C on a rotating wheel. After digestion, the MboI is heat-inactivated at 62°C for 20 minutes. The Klenow filling reaction mix is added to the sample: 37.5 µl of biotin-dATP (0.4 mM, Invitrogen #19524016), 1.5 µl of each of the other three dNTPs (10 mM, Invitrogen #18254-011, #18255-018 and #18253-013), and 10 µl of DNA Polymerase I Large (Klenow) Fragment (NEB, M0210). The total mixture is incubated at 37 °C for 1 hour with rotation. The reaction is then transferred to a 2 ml LoBind tube (VWR #CA80077-234), to which ligation mix is added: 150 μl of 10x NEB T4 DNA ligase buffer with ATP (NEB B0202), 125 μl of 10% Triton X-100, 3 μl of Ultrapure BSA (50 ng/ml, #AM2616) and 665 μl of H₂O. A 15 µL sample is stored at -20°C for input digestion. 10 µL of 400 U/µL T4 DNA ligase (NEB M0202) is added to the reaction, after which the sample is incubated at room temperature 4h with rotation. A 15 µl sample is stored at -20 °C for input ligation, after which the nuclei are pelleted at 2,200 rcf for 5 min. The nuclei are resuspended in 300 µl of Nuclear Lysis Buffer (50 mM Tris-HCl, pH 7.5; 1% SDS; 10 mM EDTA; 1X protease inhibitors) and the chromatin is released by sonication using a Qsonica Q800R3-110 for 3 minutes at 9 °C with an amplification of 20% and a pulse of 30 seconds ON and 30 seconds OFF. The samples were clarified at maximum speed and at 4 °C for 10 minutes. The resulting 300 µl of sonicated chromatin were diluted in 1,320 µl of ChIP dilution buffer (16.7 mM Tris-HCl, pH 7.5; 167 mM NaCl; 0.01% SDS; 1.1% Triton X-100; 1.2 mM EDTA). A 15 µl chromatin aliquot was set aside for the sonication control. Then, 30 µl of pre-washed Dynabeads Protein G (Invitrogen™, #10004D) and 4 µg of H3K27ac antibody (Abcam, #ab4729) were added to each tube. The tubes were then incubated overnight at 4 °C on a rotating wheel. After spinning briefly, the tubes were placed on a magnetic stand and the supernatant was removed. Then, 500 µl of Low Salt Wash Buffer (0.1% SDS; 1.0 % Triton X-100; 2 mM EDTA; 20 mM Tris-HCl, pH 7.5; 150 mM NaCl) was added to the beads on the magnet at room temperature. The beads were swished back and forth twice by moving the sample relative to the magnet. The supernatant was then removed and the procedure repeated three times. The same washing protocol was used with High Salt Wash Buffer (0.1% SDS; 1.0 % Triton X-100; 2 mM EDTA; 20 mM Tris-HCl, pH 7.5; 500 mM NaCl) and LiCl Wash Buffer (1.0 % NP-40; 1.0 % Na-Doc; 1 mM EDTA; 10 mM Tris-HCl, pH 7.5; 250 mM LiCl). After the final wash buffer was removed, 100 µl of freshly made DNA elution buffer (50 mM NaHCO3; 1% SDS) was added to the beads, which were then incubated on a low-speed vortex for 15 minutes at room temperature. After a brief spin, the tubes were placed on the magnetic stand. The supernatant was transferred to a new tube and the elution process was repeated with another 100 µL of DNA elution buffer. The second 100 µl is pooled with the first elution. There should now be 200 µl of ChIP sample. Then, 10 µl of Proteinase K (Qiagen #19131) is added and the sample is incubated at 55 °C for 45 minutes with shaking at 650 rpm. It is then raised to 67 °C for at least 1.5 hours with shaking. The DNA was then purified using the Zymo DNA Clean and Concentrator Kit (Zymo #D5205) and eluted with 12 µl of water into a 2 ml LoBind tube (VWR #CA80077-234). The purified DNA was accurately quantified using a Qubit 4 fluorometer (Invitrogen, #Q33230) with a 1X dsDNA High Sensitivity kit to calculate the amount of Tn5 (Illumina # 20034198) needed to generate libraries with the correct size distribution (Mumbach et al., 2016). The biotin pull-down is prepared by washing 5 μl of Streptavidin C-1 beads twice with 200 μl of Tween Wash Buffer (5 mM Tris-HCl, pH 7.5; 0.5 mM EDTA; 1 M NaCl; 0.05% Tween 20) in a 1.5 ml Eppendorf tube. The beads are then resuspended in 10 µl of 2x biotin binding buffer (10 mM Tris-HCl, pH 7.5; 1 mM EDTA; and 2 M NaCl) and transferred to 10 µl of samples in a 2 ml LoBind tube (VWR #CA80077-234). The tubes are placed horizontally on a rotating wheel and incubated at room temperature for 15 minutes. The drop should roll down the side of the tube. Next, the beads are placed on a magnet and the supernatant is discarded. The beads are washed twice with 500 µL of Tween Wash Buffer. They are then incubated at 55 °C for two minutes with shaking at 650 rpm. Before the Tn5 reaction, the beads are pre-washed with 100 μL of 1x (from 2x) TD buffer (Illumina # 20034198). The previously calculated amount of Tn5 is added to 25 μL of 2x TD buffer and topped up to 50 μL with water. The mixture is incubated on a Thermomixer F1.5 Eppendorf at 55 °C with intermittent shaking at 650 rpm for 10 minutes (10 seconds every 2 minutes) and then placed on a magnet to remove supernatant. 100 μL of 50 mM EDTA is added to the samples, which are then incubated at 50 °C for 30 minutes. The samples are then quickly placed on a magnet to remove the supernatant and washed twice at 50 °C for three minutes with 100 μL of 50 mM EDTA. This is followed by two washes at 55 °C for two minutes with 100 μL of Tween Wash Buffer and one final wash with 100 μL of 10 mM Tris pH 7.5. The beads are resuspended in 50 μL of PCR master mix (25 μL Phusion HF 2X (NEB #MO531S); 1.25 μL Nextera Ad1.1 [Universal primer]; 1.25 μL Nextera Ad2.x [Barcoded primer]; 22.5 μL WaterThe following PCR programme was used: 72 °C for 5 min, hot start at 98 °C for 1 min, followed by X number of cycles (depending on the amount of DNA quantified previously) of 98 °C for 15 sec, 63 °C for 30 sec, and 72 °C for 1 min (Mumbach, MR. et al., 2016). The DNA was then purified using the Zymo DNA Clean and Concentrator Kit (Zymo #D5205), and the HiChIP libraries were accurately quantified using an Invitrogen Qubit 4 Fluorometer with 1X dsDNA High Sensitivity (ThermoFisher #Q33230) and analysed using an Agilent 2100 Bioanalyzer with an Agilent DNA HS Kit (cat. no. 5067-4626). HiChIP libraries were pooled in accordance with the recommendations of the UCSD-IGM Genomics Centre (University of California, San Diego, Institute for Genomic Medicine, Genomics Centre), and sequenced using a paired-end approach with a read length of 150 bp on there Illumina NovaSeq X Plus 25B platform.

Sample digestion, ligation and sonication control (15µl) were de-crosslinked with 1% SDS final and 200mM NaCl O/N at 65C. RNAse A (Qiagen#19101) was added with 100µl TE and incubated 1h at 37°C fallowed by a proteinase K reaction (Qiagen #19131) incubated 1h at 56C. Control DNA purification was performed using the ZYMO ChIP DNA Clean and Concentrator Kit (#D5205), and DNA quality control was performed according to the Agilent DNA HS Kit protocol (cat#5067-4626).

***HiC***

HiC protocol was adapted from both protocol of Lafontaine, D. L. et al.^4^ and Gryder BE, et al.^5^ 2 million HAVICs were cross-liked for each HiC and process like a ChIP-seq protocol describe previously (this work) until the DPNII digestion 6h at RT. After digestion, DPNII is heat inactivate at 62C for 20 minutes. Is added to the sample 61.3µl of Buffer 10X #2 NEB, 52ul of Klenow filling reaction mix: 37.5µl of biotin-dATP 0.4mM (Invitrogen #19524016), 1.5µl of each 3 other dNTP 10mM (Invitrogen #18254-011, 18255-018, 18253-013) and 10µl DNA Polymerase I Large (Klenow) Fragment (NEB, M0210). The total mixture (613.3 μL) is incubated at 37 °C for 1 hour with rotation. The reaction is then transferred to a 2 ml LoBind tube (VWR #CA80077-234), to which 878 μl of ligation mix is added: 150 μl of 10x NEB T4 DNA ligase buffer with ATP (NEB B0202), 125 μl of 10% Triton X-100, 3 μl of Ultrapure BSA (50 ng/ml, #AM2616) and 600 μl of H₂O. A 15 µL sample is stored at -20°C for input digestion. 10 µL of 400 U/µL T4 DNA ligase (NEB M0202) is added to the reaction (total volume 1,486 µL), after which the sample is incubated at room temperature overnight with rotation. A 15 µl sample is stored at -20 °C for input ligation, after which the nuclei are pelleted at 2,200 rcf for 5 min. The nuclei are resuspended in 300 µl of shearing buffer (10 mM Tris-HCl, pH 8.0; 0.1% SDS; 1 mM EDTA) and the chromatin is released by sonication using a Qsonica Q800R3-110 for 7 minutes at 4 °C with an amplification of 30% and a pulse of 30 seconds on and 30 seconds off. A 15 µl sample is stored at -20°C for input sonication. Then, 10 µl of RNase A (Qiagen #19101) is added to the sample, which is incubated at 37 °C for one hour with agitation (650 rpm). Next, 30 µl of 10% SDS solution (final concentration 1%) and 10 µl of Proteinase K (Qiagen #19131) are added, and the sample is incubated at 67 °C for three hours with shaking (650 rpm). The 335 µl sample was purified using the Zymo Kit ChIP DNA Clean and Concentrator (Zymo #D5205), after which it was eluted with 12 µl of 1X TE buffer in a 2 ml LoBind tube (VWR #CA80077-234). The purified DNA is accurately quantified using an Invitrogen Qubit 4 fluorometer with a 1X dsDNA high sensitivity kit (Q33230). Biotin pull-down is prepared by washing 2 μl/μg of Streptavidin C-1 beads twice with 200 μl of Tween Wash Buffer (5 mM Tris-HCl, pH 7.5; 0.5 mM EDTA; 1 M NaCl; 0.05% Tween 20) in a 1.5 ml Eppendorf tube. The beads are then resuspended in 10 μl of 2x Biotin Binding Buffer (10 mM Tris-HCl pH 7.5, 1 mM EDTA and 2 M NaCl) and transferred to 10 μl of samples in a 2 ml LoBind tube (VWR #CA80077-234). The tubes are placed horizontally on a rotating wheel and incubated at room temperature for 15 minutes. The drop should then roll down the side of the 2 ml tube. Next, the beads are placed on a magnet, the supernatant is discarded, and the beads are washed with 400 μl of 1x biotin binding buffer. The beads are then resuspended in 100 μl of TE, transferred to a new 1.7 ml LoBind tube (VWR #CA80077-230) and separated using a magnet to remove the supernatant. Next, the beads are resuspended in 50 μl of TE in order to proceed with library preparation using the NEBNext Ultra™ II DNA Library Prep Kit for Illumina (#E7645S). The end preparation and adapter ligation steps of the NEB protocol are performed directly on the 50 μl of fragmented DNA bound to the beads, followed by a 15-minute incubation at 20 °C. After adapter ligation, the beads are separated using a magnet to remove the supernatant. They are then washed once with 200 µl of TE and resuspended in a PCR/USER enzyme master mix containing 2 µl of USER enzyme, 25 µl of NEBNext Ultra II Q5 Master Mix, 5 µl of index primer/i7 primer (with different indexes for each sample), 5 µl of universal PCR primer/i5 primer, and 13 µl of H₂O. The PCR programme begins with a 15-minute incubation at 37 °C for the USER enzyme and continues as described in the NEB protocol. After PCR, the libraries were separated using a magnet and transferred to new tubes. The libraries were then purified using SPRIselect beads at a ratio of 1x (50 μl), with two washes of 200 μl of fresh 80% ethanol. The libraries were then dried for two minutes and eluted in 12 μl of 0.1x TE. The purified DNA was accurately quantified using an Invitrogen Qubit 4 Fluorometer with 1X dsDNA High Sensitivity (#Q33230) and analysed using an Agilent 2100 Bioanalyzer with an Agilent DNA HS Kit (cat. no. 5067-4626). Hi-C libraries were pooled in accordance with the recommendations of UCSD-IGM Genomics Centre (University of California, San Diego Institute for Genomic Medicine, Genomics Centre), and sequenced using a paired-end approach with a read length of 150 bp on there Illumina NovaSeq X Plus 25B platform.

Sample digestion, ligation and sonication control (15µl) were de-crosslinked with 1% SDS final and 200mM NaCl O/N at 65C. RNAse A (Qiagen#19101) was added with 100µl TE and incubated 1h at 37°C fallowed by a proteinase K reaction (Qiagen #19131) incubated 1h at 56C. Control DNA purification was performed using the ZYMO ChIP DNA Clean and Concentrator Kit (#D5205), and DNA quality control was performed according to the Agilent DNA HS Kit protocol (cat#5067-4626)

*5’capped-small RNA-seq (5’csRNA-seq)*

A comprehensive profiling of transcription initiation was performed by identifying active transcription start sites (TSS) for HAVIC isolated from 7 donors, including 5 controls and 2 CAVD patients. 5'csRNA-seq was performed according to Duttke SH laboratory, recently publish in Nature Protocol (2026)^6^, with some minor modifications. Total RNA was extracted from a 100 mm HAVIC Petri dish with 2.5 ml of Qiazol (Qiagen #79306), transferred to an RNase-free 5 ml Eppendorf tube and 0.5 ml of chloroform was added before vigorous vortexing. After an incubation period of 2 minutes at room temperature, the sample was centrifuged at maximum speed for 15 minutes at 4 °C. The upper aqueous phase was carefully transferred to new tubes without disturbing the interphase. The volume of the aqueous phase was measured, and an equal volume of isopropanol (approximately 1.5 ml) was added for precipitation. The mixture was then incubated at -20°C overnight. The RNA was centrifuged at maximum speed for 15 minutes at 4 °C. The supernatant was then completely removed and the pellet was air-dried for 15 minutes. Finally, the RNA is resuspended in 20 µl of RNase-free water and stored at -80°C. The quality of the RNA was evaluated using a Bioanalyzer with an Agilent RNA 6000 Nano kit (part no. 5067-1511). All of the RNA used for the libraries from HAVIC had a RIN value of ≥9.2. A 15% TBE-urea gel (Bio-Rad #4566055) was pre-run in 1x TBE at 200 V for 20 minutes, with 2 µL of FLB 2x buffer (95% formamide, 5 mM EDTA, 0.1% bromophenol blue, and 0.1% xylene cyanol) loaded into the first lane. A 1:1 mixture of 10 µg of total RNA and 2x FLB is heated at 75°C for three minutes to denature, then quickly chilled on ice before loading. The RNA is loaded onto the gel with generous spacing (two empty spaces between samples) and 2.5 µl of ssRNA ladder (NEB #0364) and 2.5 µl of 2xFLB are used as a marker. The gels are run for ~40 minutes at 200 V until the bromophenol blue dye has travelled down three quarters of the way. The gels were stained for 5 minutes with SYBR™ Gold Nucleic Acid Gel Stain (Invitrogen #S11494) in TBE 1x and visualised using the BlueBox™ Pro transilluminator (MiniPCR Bio #QP-1700-03-US). The bands are cut below any strong bands that appear on the gel (approximately 55 nucleotides) and below ~20 nucleotides. They are then transferred to pre-labelled Safe-Lock 2 ml RNase-free Eppendorf tubes containing one GelBreaker Tube (IST Engineering Inc. 3388-100). The gel slices are shredded by spinning the cut gel through the Gelbreaker Tubes at maximum g for three minutes at room temperature (RT). Then, under agitation in a thermomixer at RT, 350 µl of 0.5 M NaCl (modified from Pérez, PM, 2021) are added for elution of sRNA. Next, the slurry is transferred to a spin column (UltraFree MC, 0.45 µm, Millipore UFC30HV25) using a wide orifice p1000 pipette tip. The column containing the slurry is transferred to a new 2 ml RNase-free Eppendorf LowBind tube, which contains 2 µl of GlycoBlue™ coprecipitant (ThermoFisher AM9516, 15 mg/ml). The tube is spun at 1000 g for 1 min at room temperature (RT). The column is discarded and >3x volume of 100% EtOH (1.25 ml) is added to the flow-through. This is mixed well and the small RNAs are precipitated at -20°C overnight or for one day at -80°C. The short RNAs are then precipitated by centrifugation at maximum speed at 4 °C for 45–60 minutes. The supernatant is removed and the pellet washed with 500 µl of 75% ethanol. After a second centrifugation at maximum speed and 4 °C for 20–30 minutes, all the supernatant is removed and the pellet is dried for 3–5 minutes at room temperature. The RNA is resuspended in 6 µl of TE-T buffer (0.05% Tween, 1 mM EDTA and 10 mM Tris at pH 7.5) and transferred to RNase-free, thin-walled PCR tubes. The mixture is heated to 75 °C for three minutes and then cooled on ice. 0.5 µl of each sample is set aside as 'Input' in RNase-free, thin-walled PCR tubes. Then, 1 µl of TE-T [0.05% Tween, 0.1 mM EDTA, 10 mM Tris, pH 7.5] is added to the 'Input' samples, and the mixture is stored at -80 °C for subsequent steps. The 5'-CAP enrichment process begins by adding 14.5 µl of the 1X Terminator Enzyme Master Mix (LGC Genomics, LLC #TER51020) to 5 µl of the sample. The mixture is then centrifuged briefly before being incubated at 30 °C for one hour (Meyer, M. K. et al., 2026). The tubes are briefly centrifuged before 30 µL of CIP enzyme (NEB#M0525S) and Master Mix#1 (Meyer, M.K. et al., 2026) are added, after which the tubes are incubated at 37 °C for 1 hour. The samples are then heated to 75 °C for 90 seconds and rapidly cooled on wet ice. Then, 10 µL of CIP enzyme Master Mix #2 (Meyer et al., 2026) is added to the sample and incubated for an additional 45 minutes at 37 °C. The RNA is purified by adding 500 µL of Trizol LS (Invitrogen 10296028), vortexing for five minutes, adding 150 µL of TE-T, vortexing for one minute, adding 150 µL of a 24:1 chloroform/isoamyl alcohol mixture (Sigma C0549-1PT), vortexing again, and finally centrifuging for ten minutes at 14,000 g at room temperature. The resulting supernatant (usually ~450 µL) is carefully removed without disturbing the interphase and transferred to a new tube containing 60 µL of 3 M sodium acetate and 1 µL of Glycoblue. An equivalent volume of isopropanol (typically about 550 µL) is then added, after which the RNA is precipitated overnight at -20°C. The RNA pellet is formed by spinning at >20,000 g at 4 °C for 45–60 minutes. The supernatant is removed, the pellet is washed with 500 µL of 75% ethanol and dried for 3–5 minutes. The pellet can then be frozen at -80°C or the process can proceed directly to library preparation. The libraries are prepared for samples and inputs, as described by Meyer et al. (2026). For the input, half the required volume is used. This protocol uses half the volume suggested by the NEBNext® Ultra™ II RNA Kits - RNA Library Preparation Kits (E7330S). The RNA pellets are resuspended in 3 µl of TE 'T buffer, heated for two minutes at 75 °C, and then immediately placed on ice. 5 µl of Master Mix 1, containing the RppH enzyme (NEB Cat. No. M0356S), 10x T4 RNA ligase buffer (NEB Cat. No.), PEG 8000 and SUPERase-In RNase inhibitor (Invitrogen Cat. No. AM2694), is added to the samples (Meyer et al., 2026), as well as 2.5 µl for the inputs. The samples are then incubated in a PCR machine at 37 °C for 2 hours with the lid at 42 °C. Next, for 3’ adapter ligation (green tubes NEB kit E7330), 4 µl of Master Mix 2 (Meyer, M.K. & al, 2026) is added to the samples. However, for the input, ~0,2 µl of 3’ adapter is used instead of 0.3 µl. The mixture is then incubated at 18°C O/N and lid at 40°C (RppH is inactive at 20°C and therefore does not destroy the adapter). Next, for RT primer hybridization 1 µl of 5 µM SR RT primer (0.5µl for INPUT) is added to samples on ice (the pink tube in NEB kit E7330 can be previously diluted 1 : 1 in nuclease free water) and incubated at 75°C for 2-3min, 37°C for 30min, and finally 25°C for 15min. Before the 5’ adapter ligation, the (yellow) 5´ SR adapter in the NEB kit should be resuspended in 120 µl of nuclease-free water, aliquoted into separate nuclease-free tubes and stored at –80°C. Before use, one of these tubes should be denatured by heating at 70 ˚C for 2 minutes and immediately placed on ice (use within 30 minutes). Four microlitres of ligation MM3 (Meyer et al., 2026) are added to the sample (two microlitres for input), and the mixture is incubated at 25 °C for approximately three hours. Reverse transcription is performed by adding 5.25 µL of RT-MM4 (Meyer et al., 2026) to the sample ligation reaction (2.625 µL for input) and incubating it at 50 °C for ~1.5 hours before placing it on ice. Then, 25.5 µl of PCR mix (Meyer et al., 2026) is added to each sample (12.25 µl for the inputs) plus 2 µl of a unique barcode (1 µl for the input). A 5M betaine solution (Sigma #B0300-1VL) is used at a volume of 0.3 µl in the PCR mix, along with NEB LongAmp Taq 2X Master Mix and NEBNext SR Primer for Illumina (5′) and Barcode NEBNext Unique Index Primer for Illumina (blue tubes NEB kit E7330). The PCR programme is as follows: 94°C for 3 minutes; then 14 cycles: 94 °C for 45 sec, 63 °C for 30 sec, 70 °C for 15 sec and finally 70 °C for 5 min, followed by cooling at 4 °C. The 50 µl samples and 25 µl inputs are precipitated overnight at -20 °C in ethanol, in new 2 ml RNase-free LowBind Eppendorf tubes. This is done with the addition of 1 µl of Glycoblue, 5 µl of 3 M NaOAC (0.1X) and 200 µl of 100% ethanol (>3X by volume). The DNA is pelleted at maximum speed for 45–60 minutes at 4 °C. The supernatant is then removed and the pellet washed with 500 µl of 75% ethanol. The pellet is dried for three to five minutes before being resuspended in 10 µl of water. DNA size selection and purification are performed using a 10% polyacrylamide-TBE gel containing 12 wells. First, 2 µl of 6X loading buffer are added to each sample. Then 5 µl of NEB #E7323 Quick-Load pBR322 MspI-DNA Digest (+ 2µl of 6X blue and 5 µl of H2O) is used as a marker. The input and csRNA-seq samples are loaded next to each other, which is critical for identical size selection. The gel is run for 20 min at 80V, followed by 1h15 at 180V until the Xylene Cyanole reaches the bottom edge of the plate. The adapter migrates with the lower edge of the Xylene Cyanole. The gel is stained with Syber Gold for 10-15 min minutes before being transferred to the BlueBox ™ Pro transilluminator (MiniPCR bio #QP-1700-03-US) for cutting. The slice is cut over the 124 bp adapters, so between 140-175bp (avoid the first steady-state RNA visible on the RNA gel). The gel slices are shredded by spinning them through Gelbreaker tubes [IST Engineering Inc 3388-100] which are placed in a 2ml RNase-free Eppendorf LowBind tube, and centrifuge for three minutes at RT at max speed. Then, 150 µl of diffusion buffer (0.5 M ammonium acetate, 10 mM magnesium acetate, 1 mM EDTA, pH 8.0, 0.1% SDS) is added, and elution is done by incubation O/N at 37°C on shaker at 1000 rpm (buffer modification of Meyer, M.K. & al. (2026)). The slurry is transferred to a 0.45 µm spin column (UltraFree MC, Millipore UFC30HV25) using a wide-orifice p1000 pipette tip. The gel is then rinsed with an additional 100 µl of diffusion buffer. The column containing the slurry is transferred into a new 2 ml RNase-free Eppendorf LowBind tube and spun at 1000 g for 1 minute at room temperature (RT). The spin column is discarded and the 250µl sample is transferred to a new 2 ml RNase-free Eppendorf LowBind tube for ethanol precipitation at -20 °C overnight: add 25ul NaOAC 3M (0.1X) and 1000ul 100% EtOH (>3X vol). The DNA is pelleted by spinning at >20,000 g at 4 °C for 45 minutes. The supernatant is then removed and the pellet is washed with 500 µL of 75% ethanol. The pellet is then dried for 3–5 minutes. The purified DNA is resuspended in 6 µL of H₂O and accurately quantified using an Invitrogen Qubit 4 Fluorometer with 1X dsDNA High Sensitivity (#Q33230). The profiles were evaluated using the Agilent DNA HS Kit protocol (Cat. No. 5067-4626) on a Bioanalyzer. The csRNA-seq libraries were pooled in accordance with the recommendations of the UCSD-IGM Genomics Centre (University of California, San Diego Institute for Genomic Medicine, Genomics Centre), and sequenced as 150 bp paired-end on there Illumina NovaSeq X plus platform.

***Total RNA-seq library***

A total of 500 ng of total RNA was used as the starting material for library preparation for each sample, according to the Illumina® Stranded Total RNA Prep Ligation Kit with Ribo-Zero Plus (#20040529) protocol. The purified DNA is resuspended in 6 µL of H₂O and accurately quantified using an Invitrogen Qubit 4 Fluorometer with 1X dsDNA High Sensitivity (#Q33230). The profiles were evaluated using the Agilent DNA HS Kit protocol (Cat. No. 5067-4626) on a Bioanalyzer. RNA-seq libraries were pooled in accordance with the recommendations of the UCSD-IGM Genomics Centre (University of California, San Diego Institute for Genomic Medicine, Genomics Centre), and sequenced using a paired-end approach with a read length of 150 bp on there Illumina NovaSeq X Plus 25B platform.

***RT-qPCR***

Cells were washed with PBS 1× and total RNA was extracted using the E.Z.N.A. Total RNA Kit (Omega Bio-Tek R6834-02) according to the manufacturer's instructions. RNA was quantified using a Nanodrop spectrophotometer. Reverse transcription was performed using the qScript cDNA Synthesis Kit (Quantabio 95048-500) with 1 µg of total RNA in a 20 µL reaction volume containing 4 µL of qScript reaction mix/enzyme. The thermal cycling program consisted of 22°C for 5 minutes, 42°C for 30 minutes, and 85°C for 5 minutes. The resulting cDNA was diluted 1:10 in nuclease-free water.

Quantitative PCR was performed on a Corbett Rotor-Gene 6000 using PerfeCTa SYBR Green SuperMix (Quantabio 95054-02K) in a 20 µL reaction containing 5 µL of diluted cDNA, 2 µl of Qiagen primers (HPRT1 #QT00059066, AHNAK #QT01680994, PDIA6 #QT00037086), 10 µL of PerfeCTa SYBR Green SuperMix, and 3 µL of nuclease-free water. The cycling conditions were 95°C for 15 minutes, followed by 40 cycles of 94°C for 10 seconds, 55°C for 30 seconds, and 72°C for 30 seconds. Relative gene expression was quantified using the ΔΔCt method, normalized to a housekeeping gene.

***Perturb-seq***

Perturb-seq was performed on HAVICs from 2 donors, following the CROPseq protocol^7^ for library construction and the STINGseq protocol^8^ for the plasmid and the 10X single cell 5’ CRISPR capture.

*Guide RNA library construction*

A custom single-guide RNA (sgRNA) library targeting candidate regulatory elements was designed with FlashFry V1.15^9^ within a 100 bp window around the SNV, 4 guides per target excluding guides with off targets. gRNA were then synthesized as single-stranded oligonucleotide pools (IDT oligo pool for the first experiment; Agilent HiFi OLS for the second). The lentiviral sgRNA backbone lentiGuide-FE-Puro (Addgene #170069) was linearized by BsmBI-V2 (NEB #R0739) digestion (1 µg plasmid, 1 hour at 55°C followed by 20 minutes at 80°C for heat inactivation). The digested vector was resolved on a crystal violet–stained agarose gel (1.8 µg/mL crystal violet (Sigma #C0775), no UV exposure) to preserve DNA integrity. The linearized band was excised and purified using the FavorPrep Gel/PCR Purification Mini Kit (#FAGCK 001-1), and DNA was quantified by Nanodrop.

The sgRNA oligonucleotide pool was cloned into the linearized vector by Gibson assembly using NEBuilder HiFi DNA Assembly 2X Master Mix (NEB E5520S) at a 1:200 vector-to-insert molar ratio (0.005 pmol vector, 1 pmol ssDNA oligonucleotides), incubated at 50°C for 1 hour. The assembly product was desalted by dialysis on a 0.05µm MF-Millipore™ Membrane Filter (Sigma #VMWP04700) against HPLC-grade water (Sigma #270733) for 30 minutes, then electroporated into Lucigen Endura electrocompetent cells (Lucigen #60242-1). Approximately 25 µL of the bacteria and 5 µL of DNA were transferred to pre-chilled 1 mm electroporation cuvettes and electroporated using the EC1 program (Biorad Micropulser Electroporator #1652100). Cells were immediately recovered in pre-warmed Lucigen recovery medium and incubated at 37°C with shaking 1h. Transformed bacteria were pooled and plated across ten 10 cm LB agar plates supplemented with ampicillin, yielding appropriate number of colonies (~100× coverage per sgRNA). A 1:100 dilution plate was prepared in parallel to estimate colony counts. After overnight growth at 37°C, colonies were scraped, pooled, and plasmid DNA was extracted by maxiprep (GeneBio Systems, Inc. # FAFTE 001-1- EFG).

*Library quality control*

Library representation was verified by targeted sequencing using MiniSeq Mid Output Kit (Illumina # FC-420-1004) on an Illumina MiniSeq. The sgRNA cassette was first amplified from the plasmid pool (10 ng/µL input) using indexed CROPseq library QC primers with SYBR Green Master Mix. To prevent overamplification, a 20 µL aliquot was first run on qPCR (98°C for 30 s; 40 cycles of 98°C for 10 s, 72°C for 45 s) to determine the plateau fluorescence value. The optimal PCR cycle number was calculated by dividing the plateau value by three and converting to cycle number. The remaining reaction volume was then amplified by standard PCR using the determined cycle number. PCR products were purified using SPRI beads at a 2.0× ratio, washed twice with 80% ethanol, eluted in 15 µL of 0.1× TE buffer, and quantified using an Invitrogen Qubit 4 Fluorometer with 1X dsDNA High Sensitivity (#Q33230). Fragment size was analysed using an Agilent 2100 Bioanalyzer with an Agilent DNA HS Kit (#5067-4626). Sequencing on the MiniSeq confirmed uniform representation of all sgRNAs in the library.

*CRISPRi cell line generation and perturbation*

The CRISPRi effector was delivered by transducing HAVICs with lentiviral particles carrying lentiCRISPRi(v2)-Blast (Addgene #170068; dCas9-KRAB-MeCP2) at a multiplicity of infection (MOI) of 5. HAVICs were parallelly transduced with the sgRNA lentiviral library at the desired MOI in lentivirus packaging medium supplemented with 8 µg/mL polybrene. Twenty-four hours post-transduction, medium was replaced with DMEM supplemented with 10% FBS. Puromycin and blasticidin selection was initiated 48 hours post-transduction and maintained throughout the experiment with medium changes every 2–3 days. Cells were cultured for 10 days to allow maximal CRISPRi-mediated silencing of the targeted regulatory elements.

*Single-cell RNA-seq and sgRNA capture*

Single-cell gene expression and sgRNA capture libraries were prepared using the Chromium Next GEM Single Cell 5' Kit v2 (10x Genomics #1000265) according to the manufacturer's protocol. Cells were loaded onto two Chromium Chip K lanes (10x Genomics #1000287) for the first experiment and four lanes for the second experiment. During 10X Chromium Controller cycle, cells were partitioned into gel beads-in-emulsion (GEMs), where poly-adenylated mRNA and sgRNA transcripts bearing a capture sequence were reverse-transcribed. Following GEM breakage and cDNA cleanup, full-length cDNA was amplified. The sgRNA-derived cDNA (~300 bp) was separated from the larger gene expression cDNA by SPRI Select size selection: a first round of SPRI Select bead cleanup was performed to retain the high-molecular-weight gene expression cDNA fraction on the beads, while the supernatant containing the smaller sgRNA cDNA fragments was collected. The sgRNA fraction was further purified by a second round of SPRI Select cleanup. The 5' gene expression library (10X Genomics 5' CRISPR Kit #1000451) was generated from the on-bead fraction by enzymatic fragmentation, end repair, A-tailing, adaptor ligation, and sample index PCR. The sgRNA enrichment library was constructed from the SPRI-selected small fragment fraction by sample index PCR using feature barcode-specific primers. Both libraries were accurately quantified using an Invitrogen Qubit 4 Fluorometer with 1X dsDNA High Sensitivity (#Q33230). The profiles were evaluated using the Agilent DNA HS Kit protocol (Cat. No. 5067-4626) on a Bioanalyzer. The libraries were pooled at the appropriate ratio in accordance with the recommendations of the UCSD-IGM Genomics Centre (University of California, San Diego Institute for Genomic Medicine, Genomics Centre), and sequenced as 150 bp PE on there Illumina NovaSeq X Plus platform.

*Sequencing*

This publication includes data generated at the UC San Diego IGM Genomics Center utilizing both Illumina NovaSeq 6000 and Illumina NovaSeq X Plus that were both purchased with funding from a National Institutes of Health SIG grant (#S10 OD026929).

***DPI-ELISA***

To assess transcription factor binding to specific DNA sequences, a streptavidin-capture DNA-protein binding ELISA was performed^10^. Complementary biotinylated oligonucleotide probes (IDT) were annealed at 2 µM in annealing buffer (40 mM Tris-HCl pH 8.0, 20 mM MgCl₂, 50 mM NaCl) by heating at 95°C for 5 minutes followed by slow cooling to room temperature overnight.

Nuclear protein extracts were prepared from 5–10 × 10⁶ cells (iMSC3 for TEAD1 and mAVICs for GATA4). Cells were lysed in Gough I buffer (10 mM Tris-HCl, 150 mM NaCl, 1.5 mM MgCl₂, 0.65% NP-40, 1× protease inhibitor cocktail, 10 mM DTT) and incubated on ice for 10–15 minutes. Following centrifugation (12,000 × g, 5 min, 4°C), the nuclear pellet was resuspended in high-salt buffer C (20 mM HEPES, 25% glycerol, 400 mM NaCl, 1.5 mM MgCl₂, 0.2 mM EDTA, 1× protease inhibitor cocktail, 10 mM DTT) and incubated under rotation for 2 hours at 4°C. After centrifugation, the nuclear extract supernatant was diluted with buffer D (20 mM HEPES, 20% glycerol, 400 mM KCl, 0.2 mM EDTA, 1× protease inhibitor cocktail, 10 mM DTT) and protein concentration was determined by Bradford assay (Bio Basic Kit # SK3041).

Greiner Bio-One 96-well High Binding Standard ELISA Microplates (Fisher 7000101) were coated overnight at 4°C with 100µl of streptavidin (Invitrogen 43-4301) at 10 µg/ml in PBS, then blocked with 5% BSA in TBS-T (20 mM Tris-HCl pH 7.5, 180 mM NaCl, 0.1% Tween-20) for 1 hour at room temperature. Double-stranded biotinylated probes (100 pmol per well) were immobilized at 37°C for 1 hour. After blocking with 5% BSA in TBS-T for 30 minutes, nuclear extracts (10–50 µg per well) were added and incubated for 1 hour at room temperature. Bound proteins were detected by sequential incubation with a TEAD1 (ActiveMotif #61644) or GATA-4 (ActiveMotif #39894) primary antibodies (1:500 in PBS-T, 1 hour) followed by Goat anti-Rabbit HRP-conjugated secondary antibody (TransGen #HS101-01) at 0.01 µg/ml for 1 hour. Between each step, wells were washed three times with 150µl TBS-T or PBS-T. Colorimetric detection was performed by adding 60µl of OPD (Thermo Fisher #34005) substrate solution (4 mg OPD and 3µl 30% H₂O₂ (Sigma # H1009-5ML) in 6ml citrate-phosphate buffer, pH 5.0) for up to 30 minutes in the dark. The reaction was stopped with 2N HCl and absorbance was measured at 492 nm using a microplate reader.

***Luciferase essay***

To assess the transcriptional regulatory activity of candidate enhancer elements harboring GWAS-associated variants, dual-luciferase reporter assays were performed in HAVICs using the pLenti-basP:fLuc-TK:rLuc lentiviral reporter vector (Addgene # 138369)^11^, which contains a firefly luciferase gene driven by a minimal promoter for enhancer activity measurement and a constitutively expressed Renilla luciferase cassette under the TK promoter as an internal control for lentiviral integration.

The original gateway cloning cassette was removed by double digestion with SbfI (NEB #R3642S) and ApaI (NEB #R0114S) and replaced with a unique BamHI restriction site to enable Gibson assembly-based cloning. Enhancer sequences of 200 bp centered on the variant of interest, flanked by 15 bp homology arms on each side for Gibson assembly, were synthesized (IDT) as single-stranded DNA oligonucleotides, converted to double-stranded DNA by PCR amplification (5 cycles), and cloned into the BamHI-linearized vector by Gibson assembly using NEBuilder HiFi DNA Assembly Master Mix (NEB E5520S) at a 1:7 vector-to-insert molar ratio (0.005 pmol vector). Assembly products were transformed into NEB 5-alpha competent *E. coli* by heat shock, and individual colonies were screened by Sanger sequencing to confirm correct insertion and allele identity.

Lentiviral particles carrying the reporter constructs were produced in HEK 293T cells as described above. HAVICs were transduced with lentiviral reporter particles in lentivirus packaging medium supplemented with 8 µg/mL polybrene. Three days post-transduction, cells were washed with PBS and lysed using the Passive Lysis Buffer provided in the Dual-Luciferase Reporter Assay System (Promega, #E1910). Renilla and firefly luciferase activities were measured sequentially on a microplate luminometer. Firefly luciferase activity (reporting enhancer-driven transcription) was normalized to Renilla luciferase activity (reporting lentiviral integration efficiency) to control for differences in transduction efficiency and cell number. Normalized luciferase activity was compared between reference and alternative allele constructs, as well as against a random sequence control.

***Scar-in-a-jar***

Scar-in-a-Jar is an in vitro fibrosis model that consists of highly accelerated ECM deposition by fibroblast cells under culture conditions combining macromolecular crowding (MMC) and fibrotic stimuli such as TGFβ1. High-content optical screening is then performed to quantify the collagen deposited. Our protocol is adapted from the Simon Stebler and Michael Raghunath protocol^12^. Briefly, HAVIC cells were seeded into 12-well plates at a density of 15 000 cells per well in 1ml of DMEM 10% FBS, then incubated at 37°C and 5% CO₂ for 48 hours. The medium was then replaced with 1ml of assay medium per well [DMEM supplemented with 10% FBS; 100 μM L-ascorbic acid 2-phosphate (Sigma #A8960); a mixture of 37.5 mg/mL Ficoll™ 70 (Sigma # F2878-50G) and 25 mg/mL 400 (Invitrogen # B22095.09); 5 ng/mL TGFβ1 (Gibco # PHG9214) and incubated at 37°C and 5% CO₂ for 6 days to ensure sufficient MEC deposition. The medium was renewed every 2 to 3 days with 1ml per well of freshly prepared assay medium. The collagen deposition is quantified by immunocytochemistry. Cells were first fixed 10 min at -20°C with ice-cold methanol and then wash 3 X with PBS 1X containing 0.05% Tween-20. Then blocking solution (PBS1X with 0.05% Tween-20, and 5% BSA) is added to the wells and block at room temperature for 1h with rotation. The type I collagen primary antibody (Sigma #C2456) is diluted in blocking buffer (1: 500) and add to cells for ∽ 2h incubation at RT with rotation. The primary antibody is then removed and the cells are wash three times with PBS 1X containing 0.05% Tween-20 for 10 minutes with rotation. The Alexa Fluor 647 dye conjugated to goat anti-mouse secondary antibody (Invitrogen #A-21236) is diluted in blocking buffer (1:1000), with also a 1:1000 dilution of DAPI (Thermo Fisher #62248) for nuclear staining. The cells were incubated in the dark with rotation O/N at 4°C. The next day, the cells were wash three times with PBS 1X containing 0.05% Tween-20 for 10 minutes with rotation, plus one quick wash with H2O to remove all residues. The samples are immediately analyzed or store at 4 C in the dark for not longer than 48 h prior to analysis. Zeiss microscope widefield Axio Observer Z1 driven by the Zen Pro Imaging software (Objective 10x (NA 0.25), Zeiss, ON, Canada) was used for acquisition of 16 separate images per well (4 x 4 grid) covering a total area of 0.183 cm2 were taken for each condition. The appropriate fluorescence intensity thresholds for type I collagen (CY5; Refl 50) and DAPI (Refl 49) were defined using the well with the highest fluorescence signal, and apply to all sample. The images to be analyzed are saved in .tiff format and then processed and quantified using ImageJ 1.54p (NIH, United States) as follows: 1) Nuclei are counted using the StarDist 2D plug-in on the blue channel of each resized image (Set Width: 2048). 2) Type I collagen fluorescence is quantified using the red channel of each image. After subtracting background noise, the integrated density (IntDen) of the entire image is measured (considering both intensity and surface area, the latter being similar in all images). 3) The integrated density is then reported relative to the number of nuclei. The compilation of each replicate (n=6) is plotted on a graph and statistics are calculated.

***Single-cell RNA-seq of TGFβ-induced myofibroblast differentiation***

To profile myofibroblast differentiation at single-cell resolution, HAVICs from three independent donors were seeded and cultured in Fibroblast Growth Medium 3 (PromoCell #C-23025) for 48 h to allow adaptation to the fibroblast culture conditions. Cells were then treated with either 10 ng/ml recombinant human TGFβ1 (Gibco #PHG9214) or vehicle control for 48 h. Following treatment, cells were washed with PBS, trypsinized, pelleted, and resuspended in PBS supplemented with 0.04% BSA. Cell viability and concentration were assessed by trypan blue exclusion, and only suspensions with viability >90% were processed further.

Single-cell gene expression libraries were prepared using the Chromium Next GEM Single Cell 3′ Kit v3.1 with the Chromium Next GEM Chip G Single Cell Kit (10x Genomics #1000269, #1000127) according to the manufacturer's instructions. Cells were loaded onto Chromium Next GEM Chip G lanes targeting a recovery of approximately 8,000 cells per sample. During the 10X Chromium Controller run, cells were partitioned into gel beads-in-emulsion (GEMs), where polyadenylated mRNA was captured on barcoded gel beads and reverse-transcribed to generate barcoded full-length cDNA. Following GEM breakage and cDNA cleanup with SPRIselect beads, full-length cDNA was amplified by PCR. The 3′ gene expression library was generated from the amplified cDNA by enzymatic fragmentation, end repair, A-tailing, adaptor ligation, and sample index PCR, following the manufacturer's protocol.

Libraries were accurately quantified using an Invitrogen Qubit 4 Fluorometer with the 1X dsDNA High Sensitivity assay (#Q33230), and library size profiles were evaluated using the Agilent DNA HS Kit (#5067-4626) on an Agilent 2100 Bioanalyzer. Libraries were pooled at the appropriate ratio, and sequenced as 150 bp paired-end reads on an Illumina NextSeq 2000 P3 platform.

**QUANTIFICATION AND STATISTICAL ANALYSIS**

***Statistics***

Statistical analyses were performed using R. Comparisons of continuous distributions between two groups were performed using two-sided Kolmogorov–Smirnov tests. Comparisons of continuous variables between two groups were performed using two-sided Wilcoxon rank-sum tests. Categorical comparisons between two groups (e.g., as-caQTLs versus balanced variants) were performed using two-sided Fisher's exact tests. Gene set enrichment in predefined pathways was assessed using hypergeometric tests. Linear correlations were quantified using Pearson's correlation coefficient, and Spearman's rank correlation was used when relationships were non-linear or when variables were not normally distributed. Unless otherwise specified, p-values were adjusted for multiple testing using the Benjamini–Hochberg procedure.

***RNA-seq data processing and quality control***

Paired-end RNA-seq FASTQ files were preprocessed using fastp (version 0.23.4) with default parameters for adapter trimming and quality filtering. Read quality was assessed using FastQC (version 0.12.1). Trimmed reads were aligned to the reference genome using STAR v2.7.9a in default alignment mode, producing unsorted BAM files. Aligned reads were sorted by genomic coordinate using samtools. Gene-level read quantification was performed using featureCounts from the Subread package^13^, with paired-end mode enabled (-p), allowing reads to be assigned to overlapping features (-O), and counting junction reads (-J). Reads were assigned to features defined by the specified annotation (GTF) using the designated feature type and gene identifier attribute. The resulting raw count matrix was extracted for downstream differential expression analysis.

***ATAC-seq data processing and quality control***

Raw paired-end ATAC-seq FASTQ files were preprocessed using fastp with default parameters. Reads were aligned to the hg19 reference genome using Bowtie2 (version 2.5.2) with default parameters. SAM files were converted to BAM format, sorted by genomic coordinate, and indexed using samtools (version 1.20). Peaks were called using MACS2 (version 2.2.9.1) with the parameters --broad --broad-cutoff 0.01. Inter-sample reproducibility was assessed using multiBigWigSummary from deeptools (version 3.5.4). To generate a consensus dataset, BAM files from all samples were merged using samtools merge and peaks were called on the merged file using the same parameters, yielding a total of 335,497 chromatin-accessible regions. Signal tracks (bigWig) were generated using bamCoverage from deeptools. The fraction of reads in peaks (FRiP) was calculated using bedtools (version 2.31.1) multicov to quantify the number of reads overlapping the consensus peak set.

*Transcription factor footprinting*

Transcription factor footprinting analysis was performed using TOBIAS^14^ on the merged ATAC-seq BAM file from 16 HAVIC samples. Duplicate reads were removed using Picard MarkDuplicates. Tn5 insertion bias was corrected using TOBIAS ATACorrect with the hg19 reference genome, the merged ATAC-seq consensus peak set, and ENCODE blacklist regions excluded. Footprint scores were computed across all accessible regions using TOBIAS FootprintScores on the bias-corrected signal. Transcription factor binding was detected using TOBIAS BINDetect with 755 position frequency matrices from the JASPAR 2024 CORE Homo sapiens collection, restricted to autosomal peaks. Bound sites across all motifs were concatenated and sorted to generate a genome-wide TF footprint map.

***Allele-specific chromatin accessibility analysis***

Allelic imbalance at heterozygous SNVs within ATAC-seq peaks was assessed using the following pipeline. Heterozygous SNVs were identified from ATAC-seq BAM files aligned to the hg19 reference genome. Variant calling was performed using bcftools (version 1.23) mpileup (max depth 10,000, minimum base quality 10) followed by bcftools call. Only biallelic heterozygous SNVs located on autosomes (chr1–22) with a quality score ≥ 20 were retained.

To mitigate reference mapping bias, reads overlapping heterozygous SNVs were processed through the WASP pipeline (version 0.3.4)^15^. Briefly, reads overlapping SNV positions were identified and their alleles flipped, then remapped to the reference genome using Bowtie2. Reads that did not remap to the same genomic position were discarded. Filtered reads were sorted and deduplicated using WASP's rmdup_pe module.

WASP-filtered BAM files were further processed to exclude variants falling within ENCODE blacklist regions. Remaining SNVs were annotated with rsIDs from dbSNV build 151 (common_all_20180423), and only annotated variants were retained. Allele-specific read counts at heterozygous sites were quantified using GATK ASEReadCounter (v4.1.8.1)^16^. SNVs with fewer than 5 reads supporting either allele with a minimum of 20 reads overlapping the SNV and at least 5 reads supporting each allele.

Background allelic dosage (BAD) was estimated using Babachi (version 2.0.26)^17^ to account for local copy number variation and aneuploidy. Only SNVs with a BAD score of 1 (i.e., diploid regions) overlapping ATAC-seq peaks were retained for downstream analysis.

Statistical testing for allelic imbalance was performed using MixALime(version 2.27.3)^18^. Allele-specific count data were modeled using a mixture of negative binomial distributions (MCNB). Model fitting, testing, and combination of p-values across samples were conducted using the MixALime framework (mixalime fit, test, combine). Significant SNVs (FDR < 0.10) were defined as allele-specific chromatin-accessible QTL (as-caQTL), and conversely, non-significant SNVs were defined as balanced QTL (Pval > 0.1).

***Haplotype phasing and allele-specific signal tracks***

Read-based haplotype phasing was performed using WhatsHap ^19^. For each sample, ATAC-seq BAM files were sorted and indexed with samtools, and heterozygous variants from the allelic imbalance VCF files were phased using whatshap phase with the hg19 reference genome (--ignore-read-groups). Phased VCF files were compressed with bgzip and indexed with tabix. Reads were assigned to parental haplotypes using whatshap haplotag, and haplotype-specific BAM files were generated using whatshap split. Haplotype-resolved signal tracks (bigWig) were produced from each haplotype BAM using bamCoverage (deeptools) for visualization of allele-specific chromatin accessibility at heterozygous loci.

***ChIP-seq data processing and quality control***

*Data processing*

Raw paired-end ChIP-seq FASTQ files were preprocessed using fastp with default parameters. Reads were aligned to the hg19 reference genome using Bowtie2 with default parameters. SAM files were converted to BAM format, sorted by genomic coordinate, and indexed using samtools. ChIP-seq was performed for six histone modifications (H3K27ac, H3K27me3, H3K4me1, H3K4me3, H3K36me3, H3K9me3; n = 3 biological replicates each), the transcription factor TEAD1 (n = 3), and CTCF (n = 2). Replicates from each mark were merged using samtools merge, and signal tracks (bigWig) were generated using bamCoverage from deeptools. Peak calling was performed using MACS2 with input controls for background normalization. Narrow peak calling was used for TEAD1 and CTCF. Broad peak calling (--broad --broad-cutoff 0.01) was used for all six histone modifications.

*Chromatin state annotation*

Chromatin states were defined using ChromHMM (version 1.25)^20^ with the 18-state expanded model based on the six histone marks, as recommended by the Roadmap Epigenomics Consortium. State annotations were assigned based on emission probabilities and genomic enrichment patterns following the nomenclature described in the original publication.

*Allelic imbalance of TF*

Allelic imbalance at heterozygous SNVs within ChIP-seq data was assessed using the same pipeline as described for ATAC-seq allelic imbalance analysis, with a minimum of 20 reads overlapping the SNV and at least 3 reads supporting each allele.

***HiChIP and HiC data processing and quality control***

Raw paired-end FASTQ files from four HiChIP-H3K27ac samples and two Hi-C samples were preprocessed using fastp with default parameters. Reads were then processed through the HiC-Pro pipeline (version 2.11.4)^21^ with default settings and a minimum mapping quality threshold of 10 (MIN_MAPQ = 10), using MboI as the restriction enzyme. HiC-Pro performed adapter trimming, alignment to the hg19 reference genome, assignment of reads to MboI restriction fragments, filtering for valid interaction pairs, and generation of binned contact matrices.

*HiChIP-H3K27ac Chromatin loop calling and ABC*

Chromatin loops were identified from the merged HiC-Pro output using FitHiChIP (version 8.1)^22^. Loop calling was performed in peak-to-all mode with stringent background modeling, using a bin size of 5,000 bp and considering interactions within a genomic distance of 20 kb to 2 Mb. Significant loops were identified at an FDR threshold of ≤ 0.01.

*Activity-by-Contact (ABC) model*

Enhancer–gene regulatory connections were predicted using the Activity-by-Contact (ABC) model (version 1.0.0)^23^. The ABC model integrates enhancer activity with chromatin contact frequency to quantify the regulatory effect of candidate enhancers on target gene expression. Enhancer activity was estimated by combining ATAC-seq accessibility and H3K27ac ChIP-seq signal at each candidate regulatory element. Three-dimensional contact frequency between enhancers and gene promoters was derived from Hi-C contact matrices generated by HiC-Pro. The ABC score for each enhancer-gene pair was computed as the product of enhancer activity and contact frequency, normalized by the sum of ABC scores across all candidate enhancers for a given gene. Enhancer-gene pairs with an ABC score ≥ 0.025 were retained as high-confidence regulatory connections.

***csRNA-seq data processing and quality control***

Raw csRNA-seq reads were trimmed using HOMER homerTools (version 4.11.1)^24^ to remove 3' adapter sequences (AGATCGGAAGAGCACACGTCT) with a maximum of 2 mismatches, a minimum match length of 4, and a minimum read length of 1. Trimmed reads were aligned to the hg19 reference genome using STAR v2.7.9 with parameters optimized for short capped RNA fragments (--outSAMstrandField intronMotif --outMultimapperOrder Random --outSAMmultNmax 1 --outFilterMultimapNmax 10000 --limitOutSAMoneReadBytes 10000000). Matched total RNA-seq reads were aligned with STAR v2.7.9 using default parameters. Tag directories were generated from aligned reads using HOMER makeTagDirectory for both csRNA-seq and total RNA-seq samples.

Transcription start sites (TSSs) were identified using the HOMER findcsRNATSS.pl module, which leverages the enrichment of csRNA-seq signal relative to total RNA-seq background to distinguish bona fide sites of transcription initiation from processing artifacts. TSS calling was performed using the csRNA-seq and total RNA-seq tag directories, with gene annotation from Gencode v35 (hg19 liftover).

***Genotyping and eQTL data analysis***

GenomeStudio 2.0 was used to processed genotyping data. Samples with a low call rate, sex mismatch with self-reported sex, or discordance between blood-based genotyping and RNA-based genotyping from cultured cells were excluded. Variants were filtered based on the following criteria: call rate < 0.97, minor allele frequency (MAF) < 0.01, GenCall 10th percentile < 0.1, Hardy–Weinberg equilibrium p-value < 1 E-07, or location on sex chromosomes, yielding 454,278 variants. Genotype imputation was performed using the TOPMed r3 reference panel, and imputed variants with an imputation quality score (Rsq) ≤ 0.3 were excluded.

*cis-eQTL analysis*

Genes expressed at > 1 TPM in at least 20% of samples were retained. Raw read counts were normalized across samples using the log-transformed trimmed mean of M-values (TMM) method implemented in edgeR. Principal component analysis (PCA) was performed on normalized expression values after regressing out sex and disease status. cis-eQTL mapping was performed for SNVs located within 100 kb upstream or downstream of the transcription start site (TSS) of autosomal genes (chromosomes 1–22). Only variants with MAF > 0.1 were considered. Sex, disease status, the first three expression principal components, and the first genotype principal component were included as covariates. Associations were tested using QTLtools v1.3.1 with the cis-QTL nominal pass function. Benjamini-Hochberg multiple testing correction was applied, and eQTLs with a false discovery rate (FDR) < 10% were considered significant.

*Comparison with bulk aortic valve eQTLs*

To compare the directionality of regulatory effects between HAVIC-derived eQTLs and previously reported bulk aortic valve (AV) eQTLs (n = 500)^25^, a Z-test was applied. SNVs were classified as HAVIC-specific if they reached significance in HAVICs (FDR < 0.10) but not in bulk AV tissue (p > 0.05), with a significant Z-test. SNVs were classified as having opposite regulatory effects if they were significant in both datasets (HAVIC FDR < 0.10 and AV p < 0.05) but showed discordant effect directionality, confirmed by a significant Z-test.

***TF disruption analysis***

To assess the impact of genetic variants on transcription factor (TF) binding motifs, we used PerfectosAPE (Predicting Regulatory Functional Effect of SNVs by Approximate P-value Estimation)^17^. For each variant, a 50-bp sequence centered on the SNV position was extracted from the hg19 reference genome using samtools faidx, with 25 bp of flanking sequence on each side. Both the reference and alternative alleles were encoded into the input sequence. TF binding motif models were obtained from the JASPAR 2024 CORE collection for Homo sapiens (755 position count matrices; downloaded August 12, 2024). PerfectosAPE SNVScan was run to evaluate allele-specific changes in motif binding affinity for each variant–motif pair, with (--pvalue-cutoff 0.0005 --fold-change-cutoff 5). The tool computes approximate p-values for the match of each allele to each motif and reports the fold change in binding affinity between alleles.

***Stratified LD Score Regression (S-LDSC****)*

Partitioned heritability enrichment analysis was performed using stratified LD score regression (S-LDSC)^26^ to evaluate whether genomic annotations derived from HAVIC data were enriched for the heritability of complex traits and diseases.

*Annotation file generation*. Binary annotations derived from HAVIC as-caQTL data (±500bp) were generated for each autosome (chr1–22). The baselineLD v2.2 model annotation files (downloaded from the Alkes Price lab repository)^27^ were used as the SNV coordinate template: for each chromosome, the baselineLD annotation file was parsed to extract SNV positions and all baseline annotations, and reformatted as a BED file (chr, start = position − 1, end = position). Custom annotation values were assigned to each SNV position by overlap with the input annotation intervals using bedtools map with the -c 4 -o max option, assigning a value of 1 to SNVs falling within an annotated region and 0 otherwise. The resulting file was reformatted to match the LDSC annotation format (CHR, BP, SNV, baselineLD annotations, custom annotation).

*LD score computation*. Per-chromosome LD scores were computed for each annotation using ldsc.py --l2 with the 1000 Genomes Phase 3 European reference panel (1000G.EUR), a 1 cM LD window (--ld-wind-cm 1), and restricted to HapMap3 SNVs (--print-SNVs hapmap3_SNVs/hm.$chr.SNV).

*GWAS summary statistics munging*. GWAS summary statistics were harmonized to the LDSC format using munge_sumstats.py with the HapMap3 SNV list (w_hm3.SNVlist) as the merge-alleles reference and the --a1-inc flag to indicate that the effect allele column corresponded to the increasing allele.

*Partitioned heritability enrichment*. Partitioned heritability was estimated for each GWAS trait using ldsc.py --h2 with the custom per-chromosome LD scores (--ref-ld-chr), HapMap3 regression weights excluding the HLA region (weights_hm3_no_hla/weights.), the 1000 Genomes Phase 3 European allele frequency files (1000G_Phase3_frq/1000G.EUR.QC.), and the --overlap-annot flag to account for overlap with the baselineLD v2.2 annotations. For each GWAS–annotation pair, the heritability enrichment (fold change), and the associated standard errors and p-values were extracted from the. enrichment.results output. Results were aggregated across GWAS traits and sorted by p-value.

***Phenome-wide association scans***

Phenome-wide association scans for candidate variants were performed using the Cardiovascular Disease Knowledge Portal (CVDKP; <https://cvd.hugeamp.org/>). For each variant, cross-trait associations were retrieved from the variant-level summary pages, and significant associations were reported.

***SuRE data***

SuRE (Survey of Regulatory Elements) allele-specific regulatory activity data from HepG2 cells were obtained from van Arensbergen et al. (2019)^28^. Allele-specific regulatory effects were extracted for candidate variants.

***ADASTRA allele-specific TF binding***

Allele-specific transcription factor binding (TF-ASB) calls were retrieved from the ADASTRA database (Abramov et al., 2021; <https://adastra.autosome.org/>)^17^. Variants with FDR < 0.05 were considered significant TF-ASB variants.

***UDACHA and QTLbase***

Allele-specific chromatin accessibility calls were retrieved from UDACHA^18^ (<https://udacha.autosome.org/>) by merging ATAC-seq, DNase-seq, and FAIRE-seq data; variants with FDR < 0.1 were retained. Cross-tissue molecular QTLs were retrieved from QTLbase (Zheng et al., 2020; <http://www.mulinlab.org/qtlbase>)^29^ using the variants reported in the resource. Additional caQTLs from Wenz et al. (2026)^30^ were included, retaining variants at FDR < 0.05.

***Whole-genome sequencing data***

Summary statistics from the most recent UK Biobank whole-genome sequencing association study^31^ were obtained and lifted over from GRCh38 to GRCh37/hg19 using the UCSC liftOver tool. Variants were filtered on minor allele frequency (MAF < 0.01), annotated with the Ensembl Variant Effect Predictor (VEP) to exclude exonic SNVs, and intersected with the HAVIC ATAC-seq consensus peak set using bedtools intersect to retain variants located in open chromatin regions.

***Mendelian randomization***

Two-sample Mendelian randomization (MR) analyses were performed using the LaScaMolMR Julia package (<https://samuelmathieu-code.github.io/LaScaMolMR/>)^32^. Instrumental variables (IVs) were selected from exposure GWAS summary statistics at a genome-wide significance threshold of P < 5E-08. IVs were clumped for linkage disequilibrium using 1000 Genomes Phase 3 European reference genotypes at an r² threshold of 0.01. Causal effects were estimated using four complementary methods: inverse-variance weighted (IVW), MR-Egger, and weighted median. MR-Egger intercept was used to assess the presence of pleiotropy.

***Locus plot***

Multi-track locus plots were generated using pyGenomeTracks^33^ to visualize the convergence of functional genomic annotations at candidate loci. The pyGenomeTracks framework was modified to incorporate LocusZoom-style association plots displaying GWAS −log₁₀(p-values) with linkage disequilibrium-based coloring relative to the lead variant, enabling the joint visualization of genetic association signals and underlying epigenomic landscape at each locus within a single figure.

***Tag density and signal enrichment plots***

Tag density profiles at genomic features of interest were generated using HOMER annotatePeaks.pl with the -hist option to compute average signal density across fixed-width windows centered on as-caQTLs or TSSs. The density of eQTLs and footprints in the vicinity of allele-specific chromatin accessibility QTLs (as-caQTLs) were computed using the GenomicRanges R package^34^ and visualized using ggplot2.

***Microarray data processing***

Publicly available microarray expression data from human fetal heart (GSE75985) were processed using the NetworkAnalyst platform^35^ (<https://www.networkanalyst.ca/>). Raw data were background-corrected, log₂-transformed, and VSN-normalized. Probe-to-gene mapping was performed using the platform annotation file, and probes mapping to multiple genes were removed. Differential expression analysis between OFT and the four heart chambers was performed using the limma framework, and genes with an adjusted p-value < 0.05 were considered differentially expressed.

***Predictive deep learning model analysis***

To predict the impact of genetic variants on chromatin accessibility, we fine-tuned the Enformer^36^ deep learning model on ATAC-seq data from human aortic valve interstitial cells (HAVICs). Enformer is a transformer-based architecture that takes 196,608 bp of DNA sequence as input and predicts epigenomic tracks at 128-bp resolution across 896 output bins.

We used the gReLU framework^37^ to load the pretrained Enformer model with 11 transformer layers and adapted the final prediction head for a single-task regression output corresponding to ATAC-seq signal. Because loading entire genomic sequences into memory was not feasible, we implemented a custom lazy-loading script that extracts sequences from the hg19 reference genome (FASTA) and corresponding ATAC-seq signal from a CPM-normalized BigWig file on the fly. ATAC-seq peaks from 16 merged HAVIC samples served as training intervals. Peaks overlapping ENCODE blacklist regions were excluded. Intervals whose 196,608-bp extraction window exceeded chromosome boundaries were also removed.

Training, validation, and test sets were split by chromosome, using chr10 for validation and chr11 for testing, with remaining autosomes used for training. Target signal was binned into 896 bins of 128 bp each and log-transformed (log1p) prior to training. Reverse complement data augmentation was applied during training.

Fine-tuning proceeded in two phases. In the first phase (transfer learning), only the prediction head was trained using the Adam optimizer (learning rate = 1 × 10⁻³) with a Poisson negative log-likelihood loss, where both predictions and targets were converted from log-space to count-space prior to loss computation. Early stopping with a patience of 3 epochs was applied based on validation loss. In the second phase, the 11 transformer layers and the head were unfrozen and trained end-to-end using the Adam optimizer (learning rate = 5 × 10⁻⁵) with gradient clipping (max norm = 1.0). Training was performed on a single NVIDIA RTX 5090 GPU (32 GB) with a batch size of 1.

Model performance was evaluated on the held-out test set (chr11) by computing Pearson and Spearman correlations between predicted and observed ATAC-seq signal at the three central bin of each peak, after converting values back to count-space (predictions via exp, targets via expm1).

*Variant effect prediction*

To predict the effect of genetic variants on chromatin accessibility, we used the fine-tuned Enformer model through the gReLU variant effect prediction framework. For each variant, a 196,608-bp sequence centered on the variant position was extracted for both the reference and the alternative allele. Model predictions were aggregated over three central output bins (corresponding to the region surrounding the variant) and variant effects were quantified as the log₂ fold change between alternative and reference allele predictions (log₂FC). Variants located near chromosome boundaries whose extraction window exceeded the reference genome were excluded prior to prediction. Variants with an absolute log₂FC exceeding a threshold of 0.24 (0.95 specificity versus balanced) were considered to have a significant predicted effect on chromatin accessibility

***Perturb-seq analysis***

To functionally validate candidate regulatory variants, we performed a CRISPR-based perturbation screen coupled with single-cell RNA sequencing (Perturb-seq) in HAVICs. Guide RNAs (gRNAs) were designed to target candidate regulatory elements harboring GWAS-associated variants, along with positive control gRNAs targeting transcription start sites (TSS) of known genes. Libraries were processed across two and four sequencing lanes using Cell Ranger v7.0.0 (10x Genomics) with the GRCh38-2024-A reference transcriptome, jointly quantifying gene expression and gRNA capture.

*cis-regulatory effect analysis*

Differential expression analysis of gRNA perturbations was performed using SCEPTRE (version 0.10.3)^38^. Data were jointly imported with a high multiplicity of infection (MOI) setting. gRNA-to-target assignments were defined based on genomic coordinates of the targeted regulatory elements. Positive control pairs were defined as gRNAs targeting TSS of known genes paired with the corresponding gene. Discovery pairs were constructed in cis within a 1 Mb window around each gRNA target using the construct_cis_pairs function, excluding positive control pairs. gRNA assignment was performed using a union integration strategy. Quality control filters included a mitochondrial threshold of 5%, a minimum of 15 non-zero cells in both treatment and control groups, and UMI count range between the 5th and 95th percentiles. Calibration and power checks were performed prior to the discovery analysis. left-sided tests were applied, and results were extracted from the discovery analysis.

*trans-regulatory effect analysis*

To identify distal transcriptional effects of perturbations, trans-regulatory analyses were conducted for each positive control target independently. For each targeted element, all expressed genes genome-wide were tested as potential response genes using construct_trans_pairs. The same QC parameters and two-sided testing framework were applied. Calibration checks were performed with 10 calibration pairs per target. Discovery results for each perturbation target were exported separately.

*Gene co-expression network analysis to identify programs*

To characterize transcriptional programs associated with specific perturbation targets, weighted gene co-expression network analysis (WGCNA) (version 1.73)^39^ was performed on single-cell expression data. Raw counts were first normalized by library size and denoised using MAGIC (version 3.0.0)^40^ to mitigate single-cell dropout effects. For each locus of interest, a subset of genes identified from the trans-regulatory analysis was selected. Biweight midcorrelation (bicor) was computed between all gene pairs, and hierarchical clustering (average linkage) was performed on the resulting distance matrix. Programs were defined by cutting the dendrogram into clusters, and program eigengenes were computed to summarize expression patterns.

*TF enrichment*

To identify transcription factors whose binding is enriched among genes of interest, we adapted the distance-weighted enrichment approach described in ChEA3^41^. ChIP-seq peak files for candidate transcription factors were obtained from the ChIP-Atlas database and from our HAVIC ChIP-seq. Peaks mapping to canonical chromosomes were retained and sorted by genomic position.

For each transcription factor, a binding score was computed for every protein-coding gene^41^. TSS was extracted from Gencode v35, lifted to hg19. Peak summits were defined as the midpoint of each peak interval. For each peak *i* of a given transcription factor *j* located within a 50 kb window around the transcription start site (TSS) of gene *k*, a peak score was calculated as s*ᵢ,ⱼ,ₖ* = 1 − (|d*ᵢ,ₖ*| / 50,000), where d*ᵢ,ₖ* denotes the distance between the peak summit and the TSS. For each gene–transcription factor pair, individual peak scores were summed to produce a cumulative score t*ⱼ,ₖ*. Target genes were defined as those ranked in the top 5% of nonzero cumulative scores, with a maximum cap of 1,500 targets per transcription factor.

To test whether genes identified from the CRISPRi Perturb-seq analysis or WGCNA co-expression programs were enriched among predicted transcription factor targets, we performed Fisher's exact tests. The background was defined as all protein-coding genes with at least one ChIP-seq peak within 50 kb of their TSS. Genes from the test set absent from this background were excluded. Enrichment was assessed using a two-sided Fisher's exact test, and results were reported as odds ratios with 95% confidence intervals.

***Genetically Informed Perturbation Program Score (GIPPS)***

For each co-expressed ciGene program, we examined its relationship to the risk of aortic valve disease, inspired by prior genetic aggregation approaches^42,43^. We leveraged gene-level loss-of-function (LOF) burden effect sizes derived from whole-exome sequencing in the UK Biobank (n = 394,841)^44^, using ICD code I35 (nonrheumatic aortic valve disorders). We reasoned that aggregation of LOF burden effects across genes within a program provides a genetically informed estimate of the program’s relationship to disease risk. Accordingly, for each ciGene in a program, we extracted its effect size from the LOF burden test and transformed it to reflect gene dosage by flipping the sign: β_dosage_ = −β_LOF_, representing the effect per unit increase in gene function.

Next, we harmonized each ciGene β_dosage_ to account for its direction of regulation under CRISPRi perturbation of the upstream regulator. Specifically, the sign of β_dosage_ was aligned with whether the ciGene was up- or down-regulated by the perturbation, such that the harmonized effect reflects the expected change in disease risk mediated by the perturbation (harmonized β_dosage_). For example, a gene for which increased function is associated with increased disease risk (β_dosage_ > 0) and that is downregulated by CRISPRi is expected to contribute to a reduction in disease risk at the program level considering the upstream regulation.

We defined the program effect (β_program_) as the mean of the harmonized β_dosage_ values across all ciGenes in the program, representing the genetically informed effect of perturbing the upstream regulator through that program.

Statistical significance was evaluated using an empirical null distribution. For each tested program, we sampled without replacement from the set of genes passing quality control in the single-cell screen that were also present in the exome burden dataset, drawing the same number of genes as in the program. Sampling without replacement was used to ensure that each null gene set comprised distinct genes, matching the biological structure of real programs. For each of 10,000 iterations, we computed a mean harmonized β_dosage_, yielding an empirical null distribution from which a two-sided *P*-value was obtained^45,46^.

We summarize program-level genetic evidence using a signed −log_10_ *P*-value, where the sign is determined by the direction of β_program_, referred to as the genetically informed perturbation program score (GIPPS). A GIPPS with absolute value > 1.30 corresponds to a nominal significance threshold of *P* < 0.05 (since −log_10_(0.05) ≈ 1.30). GIPPS is reported as a point estimate; confidence intervals are derived for β_program_^47,48^.

To quantify the uncertainty around β_program_, we performed a within-program bootstrap. In each of 10,000 iterations, we resampled the program’s ciGenes with replacement^5^ and computed a bootstrapped β_program_. The 2.5th and 97.5th percentiles of the resulting bootstrap distribution defined the 95% confidence interval for β_program_. Note that this resampling scheme differs by design from the without-replacement sampling used to construct the null distribution: the null assesses whether the observed program effect exceeds what would be expected from a random set of distinct background genes, whereas the bootstrap quantifies the precision of the program effect estimate given its constituent ciGenes. A confidence interval excluding zero provides corroborating evidence for the significance of the program’s genetic association with disease risk.

***Single-cell RNA-seq analysis of TGFβ-induced myofibroblast differentiation***

Raw sequencing reads from each sample (three donors × two conditions: TGFβ-treated and vehicle control) were aligned to the GRCh38-2020-A reference transcriptome and quantified using Cell Ranger v7.0.0 (10x Genomics) with default parameters. Filtered feature-barcode matrices were imported and processed using Seurat v5.3.1^49^.

*Quality control and filtering*. For each sample, a Seurat object was created from the filtered count matrix. The percentage of mitochondrial reads (MT-), ribosomal protein S reads (RPS), and ribosomal protein L reads (RPL) was computed for each cell using PercentageFeatureSet. Low-quality cells were excluded based on the following thresholds: nFeature_RNA > 500 and < 14,000, percent.mt < 5%, percent.rps < 15%, and percent.rpl < 15%. Donor and treatment condition metadata were added to each object, and cell barcodes were prefixed with donor and condition identifiers to ensure unique naming after merging.

*Data integration*. The six Seurat objects were restricted to their common feature set and merged into a single object. The RNA assay was split by donor to enable per-donor processing within Seurat v5's layer-based framework. Gene expression was normalized with NormalizeData (LogNormalize), the top 2,000 highly variable features were identified using FindVariableFeatures (vst method), and data were scaled with ScaleData. Principal component analysis was performed with RunPCA (npcs = 100). Donor batch effects were corrected using IntegrateLayers with the CCAIntegration method, using PCA as the input reduction and producing an integrated reduction (integrated.cca).

*Trajectory inference*. Single-cell trajectories were reconstructed using a two-step approach. First, a diffusion map was computed from the integrated CCA embedding using the DiffusionMap function from the destiny R package ^50^. The first two diffusion components were scaled and stored as a custom dimensional reduction (diffmap) in the Seurat object. Second, the Seurat object was converted to a SingleCellExperiment object containing both raw counts and log-normalized expression layers, with the diffusion map embedding stored as reducedDim(sce, "DM") and Seurat cluster identities used as cluster labels. Slingshot^51^ was run on the diffusion map coordinates. Pseudotime values were inverted (max − pt) to align with the expected biological direction of the trajectory and re-imported into the Seurat object as a per-cell metadata column for downstream visualization.

*Gene expression dynamics along pseudotime*. Normalized expression values for stemness markers (*PDGFRA, CD44, CXCL12, GATA4*) and fibroblast/extracellular matrix markers (*ACTA2, FAP, MYH9, SPARC, COL1A1*) were extracted from the RNA assay (data layer) and plotted against Slingshot pseudotime. Smoothed expression trends were fit using generalized additive models (GAMs) with cubic splines (y ~ s(x, k = 6)) and visualized with ggplot2.

*Pseudo-bulk differential expression*. To assess the effect of TGFβ treatment on stemness and fibroblast marker expression, pseudo-bulk profiles were generated by averaging normalized expression for each gene across all cells within each donor–condition combination. For each gene, log₂ fold changes between TGFβ and control conditions were computed per donor, and a one-sample two-sided t-test (against a null mean of 0) was performed on the per-donor log₂ fold changes to assess significance. Significance was annotated as: ** p < 0.01, * p < 0.05, ns otherwise.

**References:**

1.	Brunton, H., Garner, I. M., Bailey, U.-M., Upstill-Goddard, R. & Bailey, P. J. Using Chromatin Accessibility to Delineate Therapeutic Subtypes in Pancreatic Cancer Patient-Derived Cell Lines. *STAR Protoc.* **1**, 100079 (2020).

2.	Arrigoni, L. *et al.* RELACS nuclei barcoding enables high-throughput ChIP-seq. *Commun. Biol.* **1**, 214 (2018).

3.	Mumbach, M. R. *et al.* HiChIP: efficient and sensitive analysis of protein-directed genome architecture. *Nat. Methods* **13**, 919–922 (2016).

4.	Lafontaine, D. L., Yang, L., Dekker, J. & Gibcus, J. H. Hi-C 3.0: Improved Protocol for Genome-Wide Chromosome Conformation Capture. *Curr. Protoc.* **1**, e198 (2021).

5.	Gryder, B. E. *et al.* Miswired Enhancer Logic Drives a Cancer of the Muscle Lineage. *iScience* **23**, 101103 (2020).

6.	Meyer, M. K. *et al.* Profiling active RNA polymerase II transcription start sites from total RNA by capped small RNA sequencing (csRNA-seq). *Nat. Protoc.* 1–25 (2026) doi:10.1038/s41596-025-01285-y.

7.	Datlinger, P. *et al.* Pooled CRISPR screening with single-cell transcriptome readout. *Nat. Methods* **14**, 297–301 (2017).

8.	Morris, J. A. *et al.* Discovery of target genes and pathways at GWAS loci by pooled single-cell CRISPR screens. *Science* **380**, eadh7699 (2023).

9.	McKenna, A. & Shendure, J. FlashFry: a fast and flexible tool for large-scale CRISPR target design. *BMC Biol.* **16**, 74 (2018).

10.	Brand, L. H., Kirchler, T., Hummel, S., Chaban, C. & Wanke, D. DPI-ELISA: a fast and versatile method to specify the binding of plant transcription factors to DNA in vitro. *Plant Methods* **6**, 25 (2010).

11.	Sissaoui, S. *et al.* Genomic Characterization of Endothelial Enhancers Reveals a Multifunctional Role for NR2F2 in Regulation of Arteriovenous Gene Expression. *Circ. Res.* **126**, 875–888 (2020).

12.	Stebler, S. & Raghunath, M. The Scar-in-a-Jar: In Vitro Fibrosis Model for Anti-Fibrotic Drug Testing. *Methods Mol. Biol.* **2299**, 147–156 (2021).

13.	Liao, Y., Smyth, G. K. & Shi, W. featureCounts: an efficient general purpose program for assigning sequence reads to genomic features. *Bioinformatics* **30**, 923–930 (2014).

14.	Bentsen, M. *et al.* ATAC-seq footprinting unravels kinetics of transcription factor binding during zygotic genome activation. *Nat. Commun.* **11**, 4267 (2020).

15.	WASP: allele-specific software for robust molecular quantitative trait locus discovery | Nature Methods. https://www.nature.com/articles/nmeth.3582.

16.	Van der Auwera, G. A. *et al.* From FastQ data to high confidence variant calls: the Genome Analysis Toolkit best practices pipeline. *Curr. Protoc. Bioinforma.* **43**, 11.10.1-11.10.33 (2013).

17.	Abramov, S. *et al.* Landscape of allele-specific transcription factor binding in the human genome. *Nat. Commun.* **12**, 2751 (2021).

18.	Buyan, A. *et al.* Statistical framework for calling allelic imbalance in high-throughput sequencing data. *Nat. Commun.* **16**, 1739 (2025).

19.	Martin, M. *et al.* WhatsHap: fast and accurate read-based phasing. 085050 Preprint at https://doi.org/10.1101/085050 (2016).

20.	Ernst, J. & Kellis, M. Chromatin-state discovery and genome annotation with ChromHMM. *Nat. Protoc.* **12**, 2478–2492 (2017).

21.	Servant, N. *et al.* HiC-Pro: an optimized and flexible pipeline for Hi-C data processing. *Genome Biol.* **16**, 259 (2015).

22.	Bhattacharyya, S., Chandra, V., Vijayanand, P. & Ay, F. Identification of significant chromatin contacts from HiChIP data by FitHiChIP. *Nat. Commun.* **10**, 4221 (2019).

23.	Fulco, C. P. *et al.* Activity-by-contact model of enhancer–promoter regulation from thousands of CRISPR perturbations. *Nat. Genet.* **51**, 1664–1669 (2019).

24.	Duttke, S. H., Chang, M. W., Heinz, S. & Benner, C. Identification and dynamic quantification of regulatory elements using total RNA. *Genome Res.* **29**, 1836–1846 (2019).

25.	Thériault, S. *et al.* Integrative genomic analyses identify candidate causal genes for calcific aortic valve stenosis involving tissue-specific regulation. *Nat. Commun.* **15**, 2407 (2024).

26.	Finucane, H. K. *et al.* Partitioning heritability by functional annotation using genome-wide association summary statistics. *Nat. Genet.* **47**, 1228–1235 (2015).

27.	Gazal, S. *et al.* Linkage disequilibrium–dependent architecture of human complex traits shows action of negative selection. *Nat. Genet.* **49**, 1421–1427 (2017).

28.	van Arensbergen, J. *et al.* High-throughput identification of human SNPs affecting regulatory element activity. *Nat. Genet.* **51**, 1160–1169 (2019).

29.	QTLbase: an integrative resource for quantitative trait loci across multiple human molecular phenotypes | Nucleic Acids Research | Oxford Academic. https://academic.oup.com/nar/article/48/D1/D983/5584691.

30.	Wenz, B. M. *et al.* Expanded chromatin accessibility mapping explains genetic variation associated with complex traits in liver. *Am. J. Hum. Genet.* **113**, 260–275 (2026).

31.	Carss, K. *et al.* Whole-genome sequencing of 490,640 UK Biobank participants. *Nature* **645**, 692–701 (2025).

32.	Mathieu, S. *et al.* Efficient molecular mendelian randomization screens with LaScaMolMR.jl. 2024.08.29.24312805 Preprint at https://doi.org/10.1101/2024.08.29.24312805 (2024).

33.	Lopez-Delisle, L. *et al.* pyGenomeTracks: reproducible plots for multivariate genomic datasets. *Bioinformatics* **37**, 422–423 (2021).

34.	Lawrence, M. *et al.* Software for Computing and Annotating Genomic Ranges. *PLOS Comput. Biol.* **9**, e1003118 (2013).

35.	Zhou, G. *et al.* NetworkAnalyst 3.0: a visual analytics platform for comprehensive gene expression profiling and meta-analysis. *Nucleic Acids Res.* **47**, W234–W241 (2019).

36.	Avsec, Ž. *et al.* Effective gene expression prediction from sequence by integrating long-range interactions. *Nat. Methods* **18**, 1196–1203 (2021).

37.	Lal, A., Gunsalus, L., Nair, S., Biancalani, T. & Eraslan, G. gReLU: a comprehensive framework for DNA sequence modeling and design. *Nat. Methods* **22**, 2253–2257 (2025).

38.	Barry, T., Wang, X., Morris, J. A., Roeder, K. & Katsevich, E. SCEPTRE improves calibration and sensitivity in single-cell CRISPR screen analysis. *Genome Biol.* **22**, 344 (2021).

39.	Langfelder, P. & Horvath, S. WGCNA: an R package for weighted correlation network analysis. *BMC Bioinformatics* **9**, 559 (2008).

40.	van Dijk, D. *et al.* Recovering Gene Interactions from Single-Cell Data Using Data Diffusion. *Cell* **174**, 716-729.e27 (2018).

41.	Keenan, A. B. *et al.* ChEA3: transcription factor enrichment analysis by orthogonal omics integration. *Nucleic Acids Res.* **47**, W212–W224 (2019).

42.	Ota, M. *et al.* Causal modelling of gene effects from regulators to programs to traits. *Nature* **650**, 399–408 (2026).

43.	Schnitzler, G. R. *et al.* Convergence of coronary artery disease genes onto endothelial cell programs. *Nature* **626**, 799–807 (2024).

44.	Karczewski, K. J. *et al.* Systematic single-variant and gene-based association testing of thousands of phenotypes in 394,841 UK Biobank exomes. *Cell Genomics* **2**, 100168 (2022).

45.	Phipson, B. & Smyth, G. K. Permutation P-values should never be zero: calculating exact P-values when permutations are randomly drawn. *Stat. Appl. Genet. Mol. Biol.* **9**, Article39 (2010).

46.	North, B. V., Curtis, D. & Sham, P. C. A Note on the Calculation of Empirical P Values from Monte Carlo Procedures. *Am. J. Hum. Genet.* **71**, 439–441 (2002).

47.	DiCiccio, T. J. & Efron, B. Bootstrap confidence intervals. *Stat. Sci.* **11**, 189–228 (1996).

48.	Efron, B. & Tibshirani, R. J. *An Introduction to the Bootstrap*. (Chapman and Hall/CRC, New York, 1994). doi:10.1201/9780429246593.

49.	Hao, Y. *et al.* Dictionary learning for integrative, multimodal and scalable single-cell analysis. *Nat. Biotechnol.* **42**, 293–304 (2024).

50.	Angerer, P. *et al.* destiny: diffusion maps for large-scale single-cell data in R. *Bioinformatics* **32**, 1241–1243 (2016).

51.	Street, K. *et al.* Slingshot: cell lineage and pseudotime inference for single-cell transcriptomics. *BMC Genomics* **19**, 477 (2018).

**Supplementary Legends**

**Supplementary Figure 1: ATAC-seq quality.** **A**, Inter-sample correlation of ATAC-seq signal across biological replicates, supporting the definition of a consensus set of 326,594 chromatin-accessible regions.

**Supplementary Figure 2: as-caQTL identification and chromatin state annotation.** **A**, Schematic of the allelic imbalance approach used to identify as-caQTLs from ATAC-seq at heterozygous SNVs. **B**, Distribution of reference versus alternative allele read counts at heterozygous SNVs; yellow: as-caQTLs, blue: balanced SNVs. **C**, Number of as-caQTLs discovered as a function of sequencing read depth, based on subsampling of the top-sequenced sample. **D**, 18-state ChromHMM model characteristics, derived from ChIP-seq of six histone marks in HAVICs.

**Supplementary Figure 3: 5′csRNA-seq and transcription initiation at as-caQTLs.** **A**, Sequence logo at transcription start sites (TSSs) defined by 5′csRNA-seq, showing the canonical initiator YR dinucleotide. **B**, Distribution of TATA box motif frequency relative to TSSs, peaking 20–30 nucleotides upstream. **C**, Spatial organization of TF motifs around TSSs, distinguishing activator (AP-1, KLF, NFY) and repressor (YY1) positioning. **D**, Empirical cumulative distribution function (ECDF) of as-caQTL position relative to TSSs compared to balanced SNVs and random expectation (Kolmogorov–Smirnov test). **E**, Normalized 5′csRNA-seq read counts within ±1,000 bp of as-caQTLs versus balanced SNVs.

**Supplementary Figure 4: as-caQTLs, reporter activity and gene expression.** **A**, Correlation between allelic effects on chromatin accessibility and SuRE reporter expression across P_SuRE_ < 1E-5 SNVs. **B**, Distribution of mean eQTL density per 10-bp bin around as-caQTLs versus balanced SNVs (P < 1E-308, Kolmogorov–Smirnov test). **C**, Distribution of minor allele frequency (MAF) for as-caQTLs shared with AV eQTLs (as-ca-AVeQTLs) versus unshared as-caQTLs (P=8.52E-49, Wilcoxon rank sum test). **D**, Enrichment of as-ca-AVeQTLs (yellow) versus unshared as-caQTLs (blue) across ChromHMM chromatin states (Fisher's exact test). **E**, Allelic imbalance effect sizes of as-ca-AVeQTLs compared to unshared as-caQTLs. (P=9.99E-32, Wilcoxon rank sum test) **F**, Stratified LD score regression significance (−log₁₀ P) for as-caQTLs across GTEx tissue eQTLs and AV eQTLs. **G**, LDSC heritability enrichment for Aortic Valve eQTL as a function of the number of top-ranked as-caQTLs.

**Supplementary Figure 5: Enformer model performance and variant prioritization at GWAS loci.** **A**, Correlation between Enformer-predicted allelic fold-changes and observed as-caQTL effects for variants uniquely assigned within ATAC-seq peaks (Spearman's ρ = 0.67). **B**, ROC curve for classification of as-caQTLs versus balanced SNVs. **C**, CADD scores of Enformer-predicted as-caQTLs compared to predicted balanced SNVs (P=1.6E-09, Wilcoxon rank-sum test). **D**, Locus plot of predicted as-caQTL rs178004. Tracks show, from top to bottom: CAVD GWAS LocusZoom, H3K27ac HiChIP loops, gene annotations, ATAC-seq, 5′csRNA-seq, H3K27ac, H3K4me1, H3K4me3 and CTCF ChIP-seq, and GERP conservation score. In silico saturation mutagenesis reveals a composite regulatory element containing two canonical AP-1 motifs; attention weight analysis of the T and C alleles shows that the risk C allele disrupts the motif and attenuates enhancer activity.

**Supplementary Figure 6: Quality control of the first CRISPRi gRNA library.** **A**, Number of detected genes per cell; purple bar indicates quality control threshold. **B**, Number of UMIs per cell. **C**, Proportion of mitochondrial reads per cell. **D**, Example of cell assignment as perturbed or unperturbed based on target gRNA read counts. **E**, Number of cells per gRNA (mean = 208). **F**, Number of gRNAs per cell (MOI = 6.81). **G**, −log₁₀(P-value) of perturbation effects on positive control genes versus negative controls. **H**, Distribution of non-targeting and targeting cell counts for each SNV–gene pair. **I**, Quantile–quantile (QQ) plot of CRISPRi discovery P-values (blue) versus non-targeting control P-values (red). **J**, Volcano plot of the first library results; yellow points indicate significant associations (FDR < 0.1).

**Supplementary Figure 7: Quality control of the second CRISPRi gRNA library.** **A**, Number of detected genes per cell; purple bar indicates quality control threshold. **B**, Number of UMIs per cell. **C**, Proportion of mitochondrial reads per cell. **D**, Example of cell assignment as perturbed or unperturbed based on target gRNA read counts. **E**, Number of cells per gRNA (mean = 575). **F**, Number of gRNAs per cell (MOI = 23.41). **G**, −log₁₀(P-value) of perturbation effects on positive control genes versus negative controls. **H**, Distribution of non-targeting and targeting cell counts for each SNV–gene pair. **I**, Quantile–quantile (QQ) plot of CRISPRi discovery P-values (blue) versus non-targeting control P-values (red).

**Supplementary Figure 8: Summary of as-caQTL CRISPRi screen results.** Overview of all SNV–ciGene associations identified across the CRISPRi perturbation screen.

**Supplementary Figure 9: CRISPRi identifies *PDGFRA* as a target gene at a CAVD risk locus lacking eQTL support.** **A**, CRISPRi of the promoter variant rs1800813 (P_GWAS_ = 1.39E-19; LD r² = 1.0 with the lead SNV) downregulates *PDGFRA* (log₂FC = −0.17, P = 1.5E-9). Tracks show, from top to bottom: CAVD GWAS LocusZoom, H3K27ac HiChIP loops, gene annotations, ATAC-seq, 5′csRNA-seq, H3K27ac, H3K4me1, H3K4me3 and CTCF ChIP-seq, and GERP conservation score. **B**, Luciferase reporter assay showing that the protective G allele increases transcriptional activity (Wilcoxon rank-sum test, n = 6 replicates from 2 donors, *P<0.05). **C**, Log₂ fold-change (TGF-β/control) of pseudobulk expression from single-cell RNA-seq for myofibroblast and stemness marker genes (n = 3 donors) (t-test, *P<0.05, **P<0.01). **D**, Pseudotime ordering based on diffusion map dimensionality reduction. **E**, Expression of stemness genes across pseudotime. **F**, Expression of myofibroblast genes across pseudotime.

**Supplementary Figure 10: Developmental context of the *ADAMTSL4* locus and Mendelian randomization.** **A**, Single-cell ATAC-seq co-accessibility linking rs6693567-RE to the *ADAMTSL4* promoter in fetal cardiac stromal cells; the relevant link is highlighted in red. **B**–**F**, Gene expression across the four cardiac chambers (left atrium (LA), left ventricle (LV), right atrium (RA), right ventricle (RV)) and the outflow tract (OFT) during embryogenesis (fetal heart, week 11) for *ADAMTSL4* (**B**), *ELN* (**C**), *LOX* (**D**), *FBN1* (**E**) and *FBLN5* (**F**). **G**, Mendelian randomization showing that higher AoVmax is positively associated with CAVD risk (β = 0.15, P_IVW_ = 3.0E-3).

**Supplementary Figure 11: Convergence of common regulatory variants at fibrillin genes.** **A**, CRISPRi of rs363832/rs3929051-RE identifies *FBN1* as the target gene (log₂FC = −0.12, P = 1.03E-7). Tracks show, from top to bottom: iAoAstj GWAS LocusZoom, gene annotations, ATAC-seq, 5′csRNA-seq, H3K27ac, H3K4me1, H3K4me3 and CTCF ChIP-seq, and GERP conservation score. **B**, CRISPRi of rs73348278-RE and rs78821460-RE reduced expression of *FBN2* (log2FC = −0.64, P_SCEPTRE_ = 2.1E−6 and log2FC = −0.80, P_SCEPTRE_ = 3.2E-09). Tracks show, from top to bottom: iAoAstj GWAS LocusZoom, FBN2 eQTL LocusZoom, gene annotations, ATAC-seq, 5′csRNA-seq, H3K27ac, H3K4me1, H3K4me3 and CTCF ChIP-seq, and GERP conservation score.

**Supplementary Figure 12: Representation of the GIPPS framework and omnigenic model. A, Schematic representation of the GIPPS methodology. B, Schematic representation of the omnigenic model.**

**Supplementary Figure 13: Core HAVIC programs: TEAD1 enrichment, functional validation and *RNFT1*-regulated iron transport program.** **A**, Enrichment of the three programs for SMAD3 ChIP peaks (ChIP-Atlas) and TEAD1 ChIP-seq peaks in HAVICs. **B**, Full-field images of the scar-in-a-jar collagen production assay (the main figure shows a zoomed view). **C**, RT-qPCR quantification of *AHNAK* expression (normalized to *HPRT1*) in control HAVICs versus *AHNAK* CRISPRi HAVICs (n = 6; Wilcoxon rank-sum test, ** P ≤ 0.01). **D**, RT-qPCR quantification of *PDIA6* expression (normalized to *HPRT1*) in control HAVICs versus *PDIA6* CRISPRi HAVICs (n = 6; Wilcoxon rank-sum test, ** P ≤ 0.01). **E**, Example locus: CRISPRi of rs76419616-RE (P_GWAS_ = 3.87E-6) downregulates *RNFT1* in cis (log₂FC = −0.85, P = 1.11E-16). Tracks show, from top to bottom: CAVD GWAS LocusZoom, gene annotations, H3K27ac HiChIP loops, ATAC-seq, 5′csRNA-seq, H3K27ac, H3K4me1, H3K4me3 and CTCF ChIP-seq, and GERP conservation score. **F**, Heatmap of co-expression among 103 trans-regulated ciGenes downstream of rs76419616-RE perturbation, with CRISPRi effect sizes and hierarchical clustering identifying two programs (blue and red). **G**, Reactome pathway enrichment analysis for each program (top three enrichments per program; hypergeometric test). **H**, Forest plot of the genetically informed perturbation program score (GIPPS) for the two programs; the blue iron uptake and transport program containing *RNFT1* is associated with CAVD risk (GIPPS = 2.41).

**Supplementary Figure 14: Summary.** Schematic representation of key findings, linking target genes to biological pathways and to disease traits. Solid lines indicate functional evidence provided in this study; dashed lines indicate associations supported by the literature.

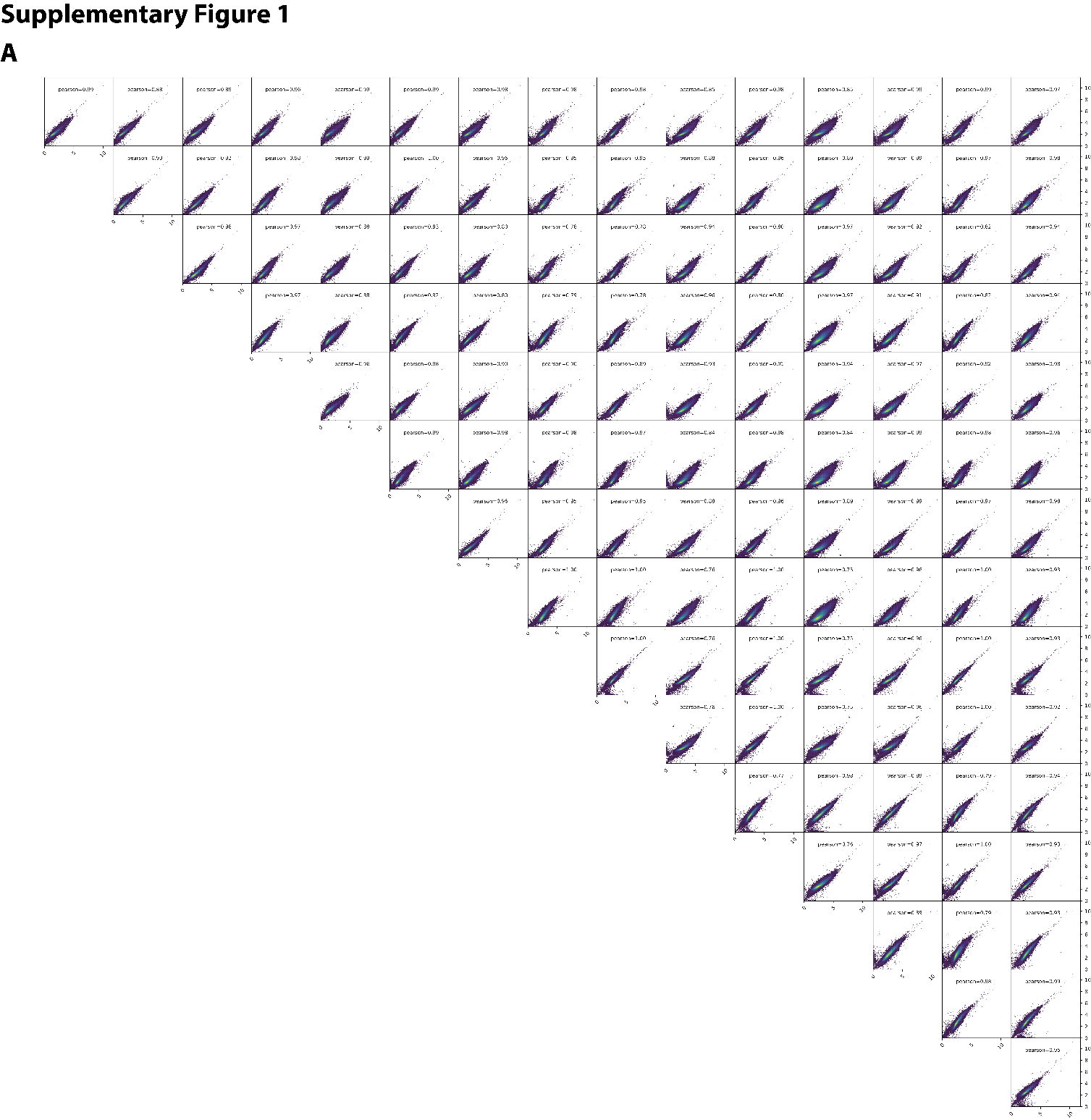
**Supplementary Figures**

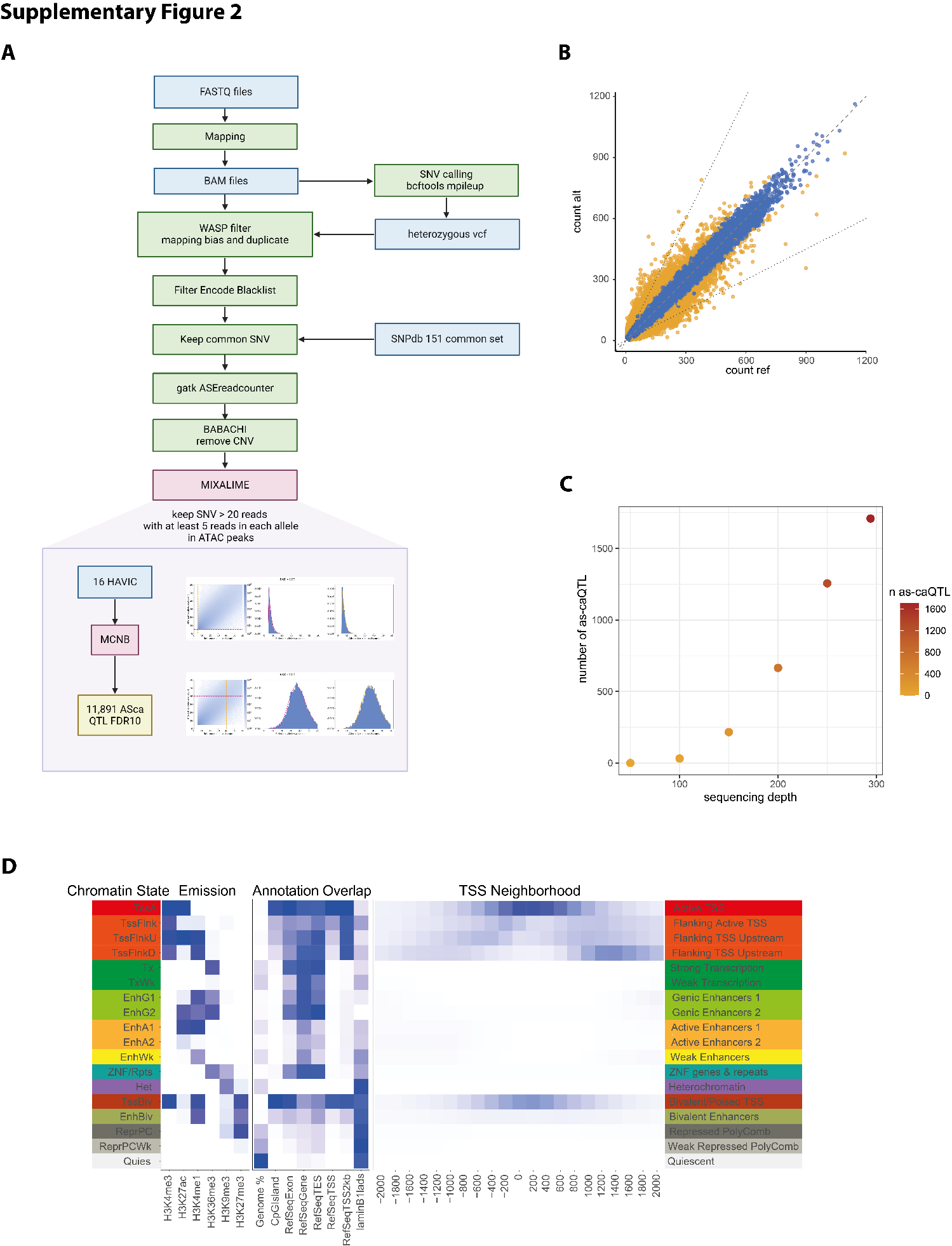

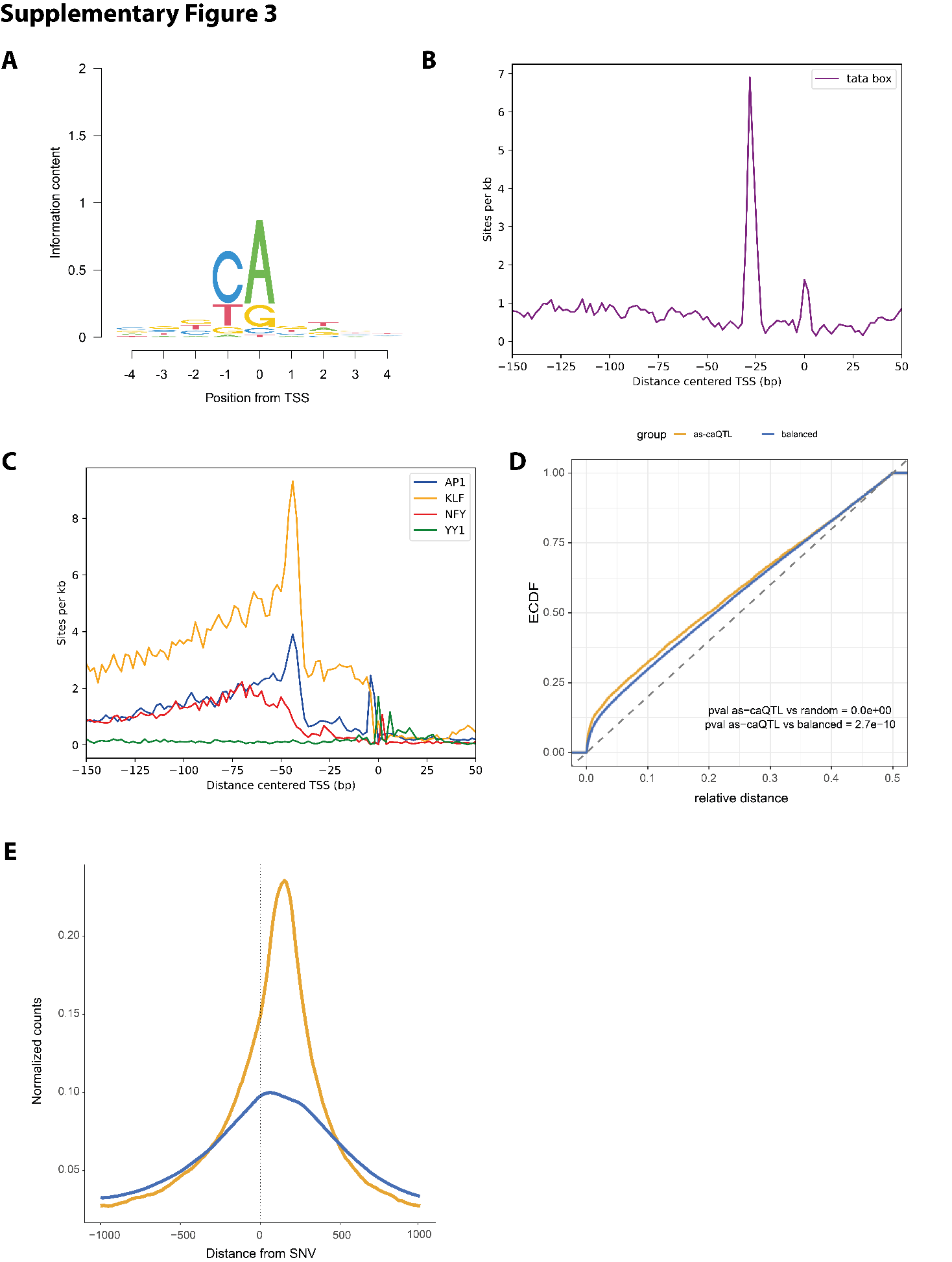

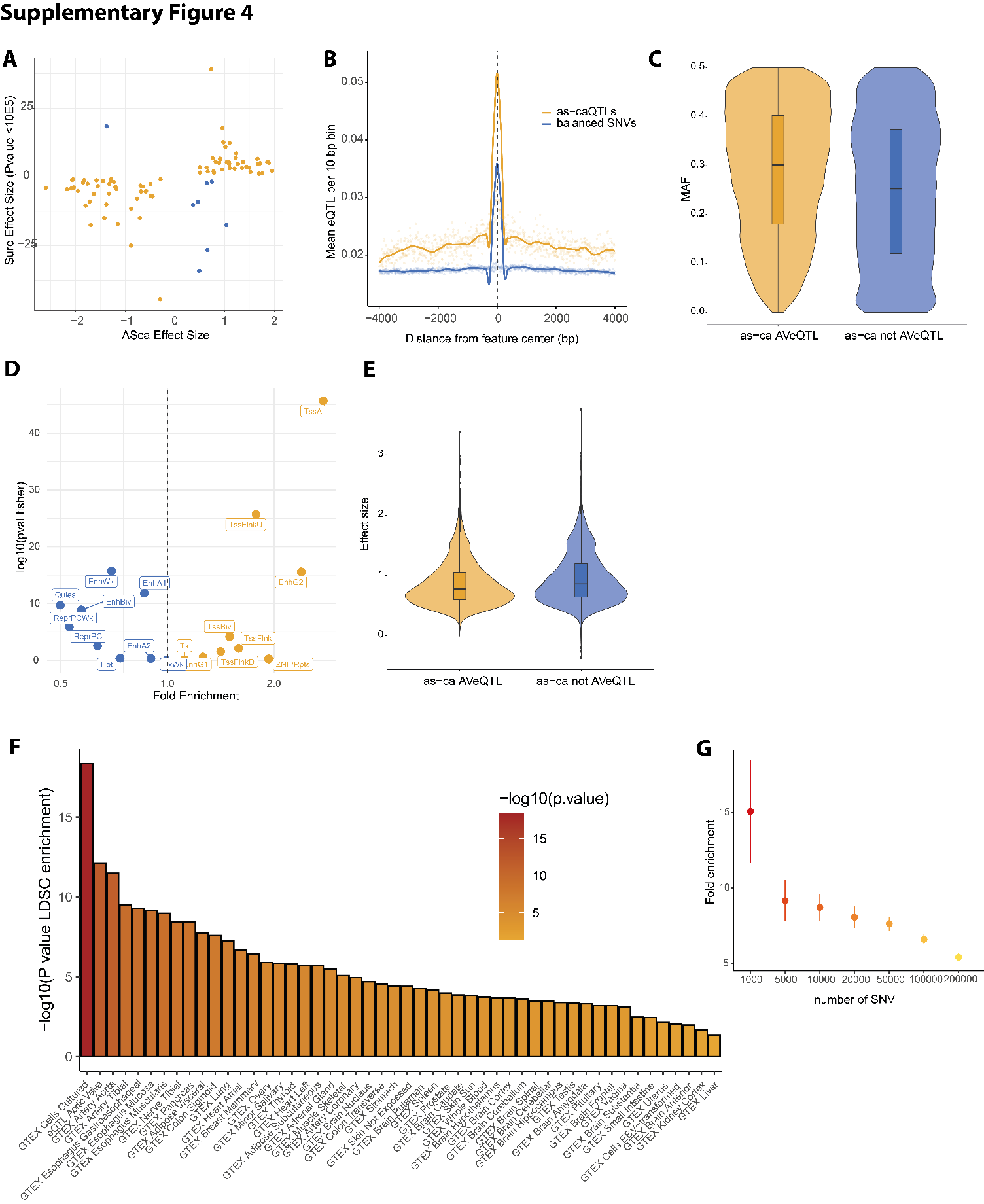

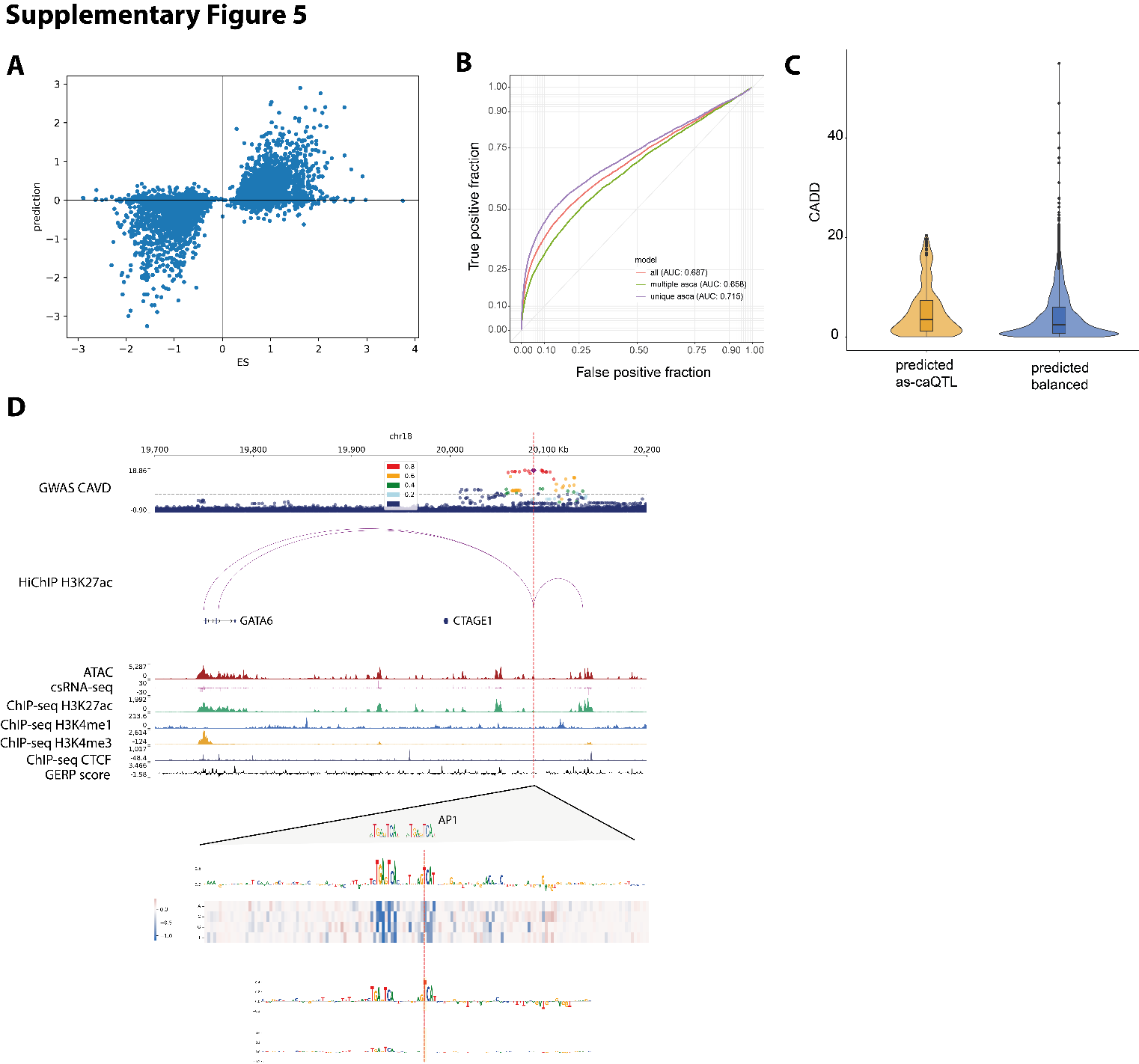

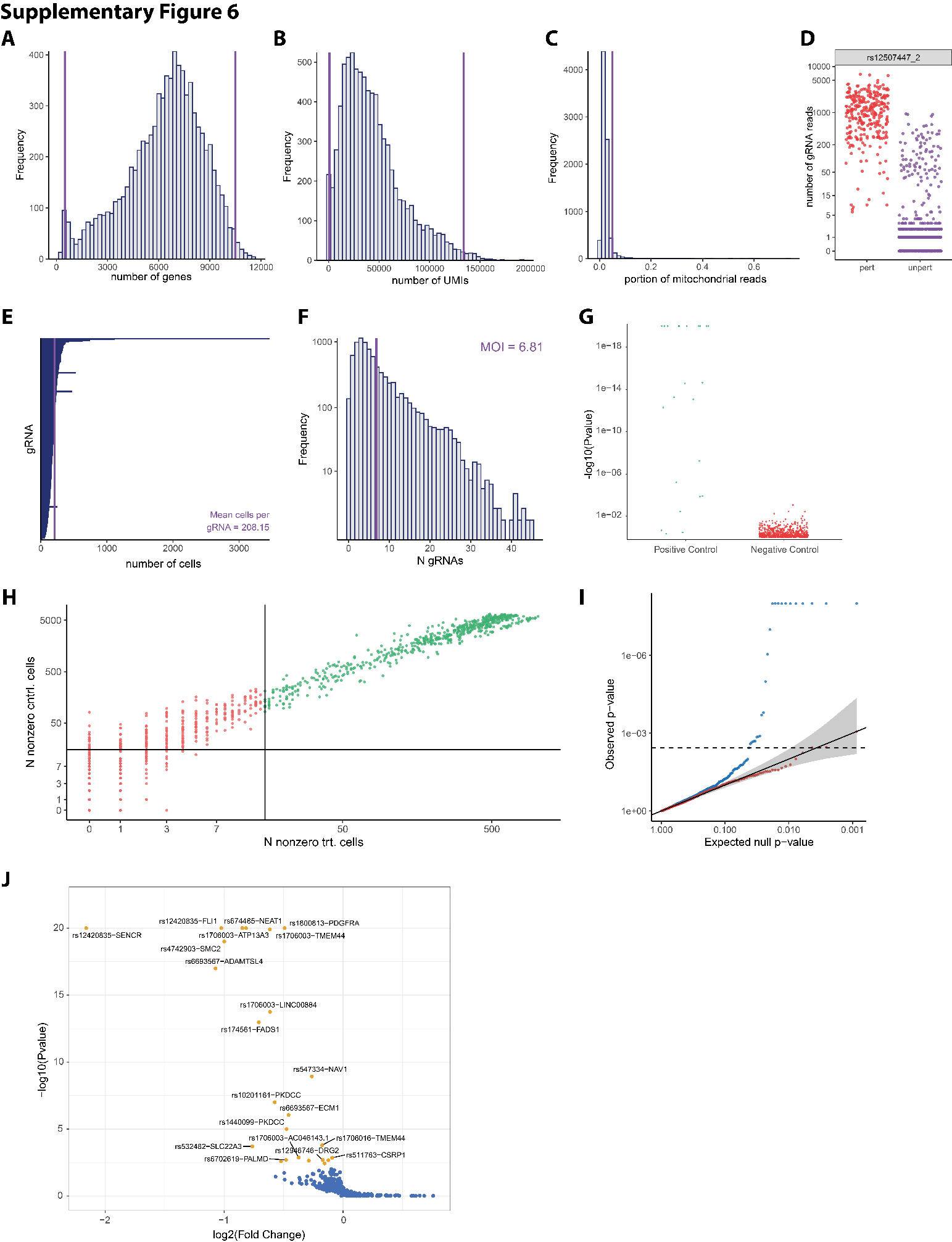

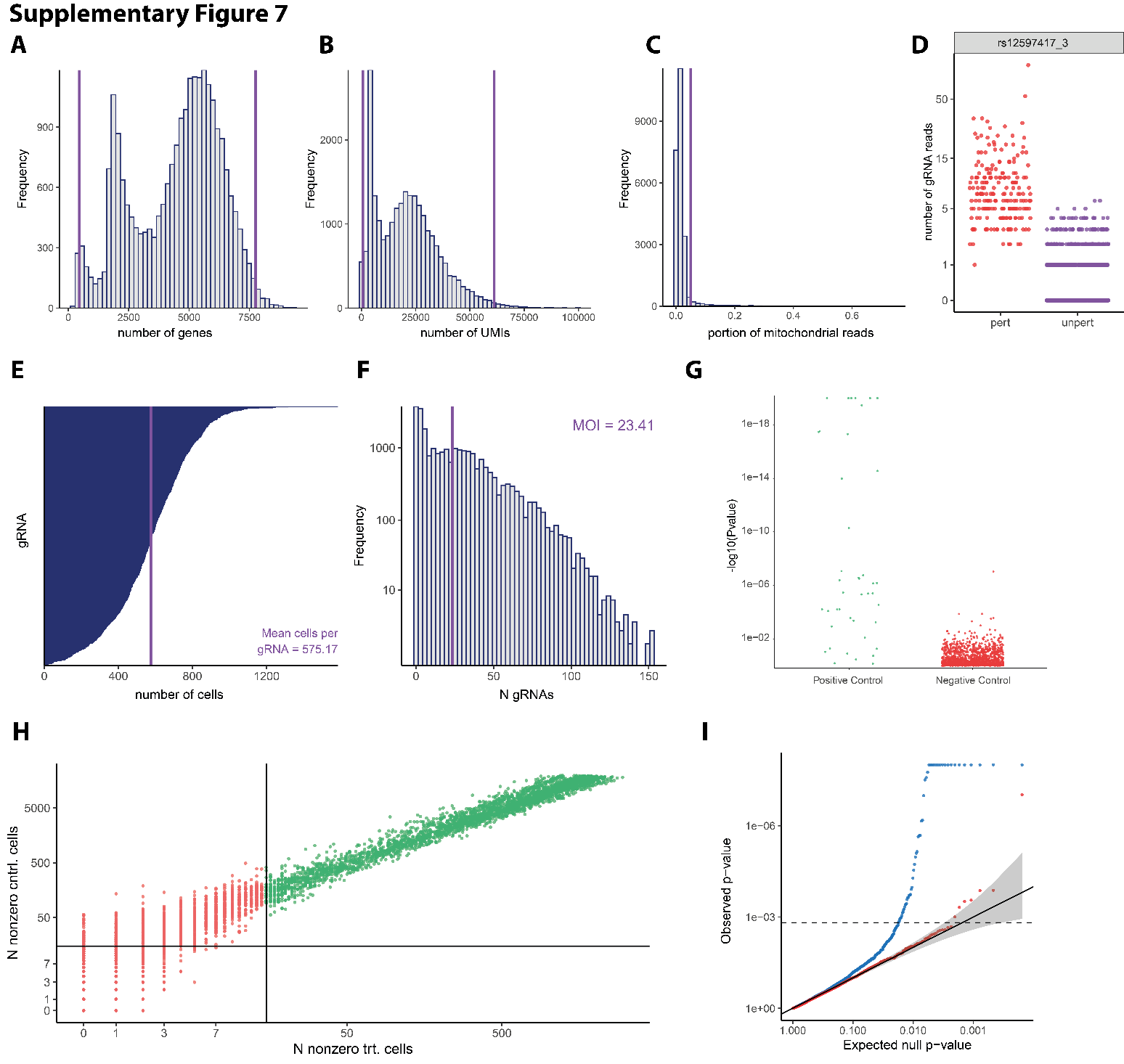

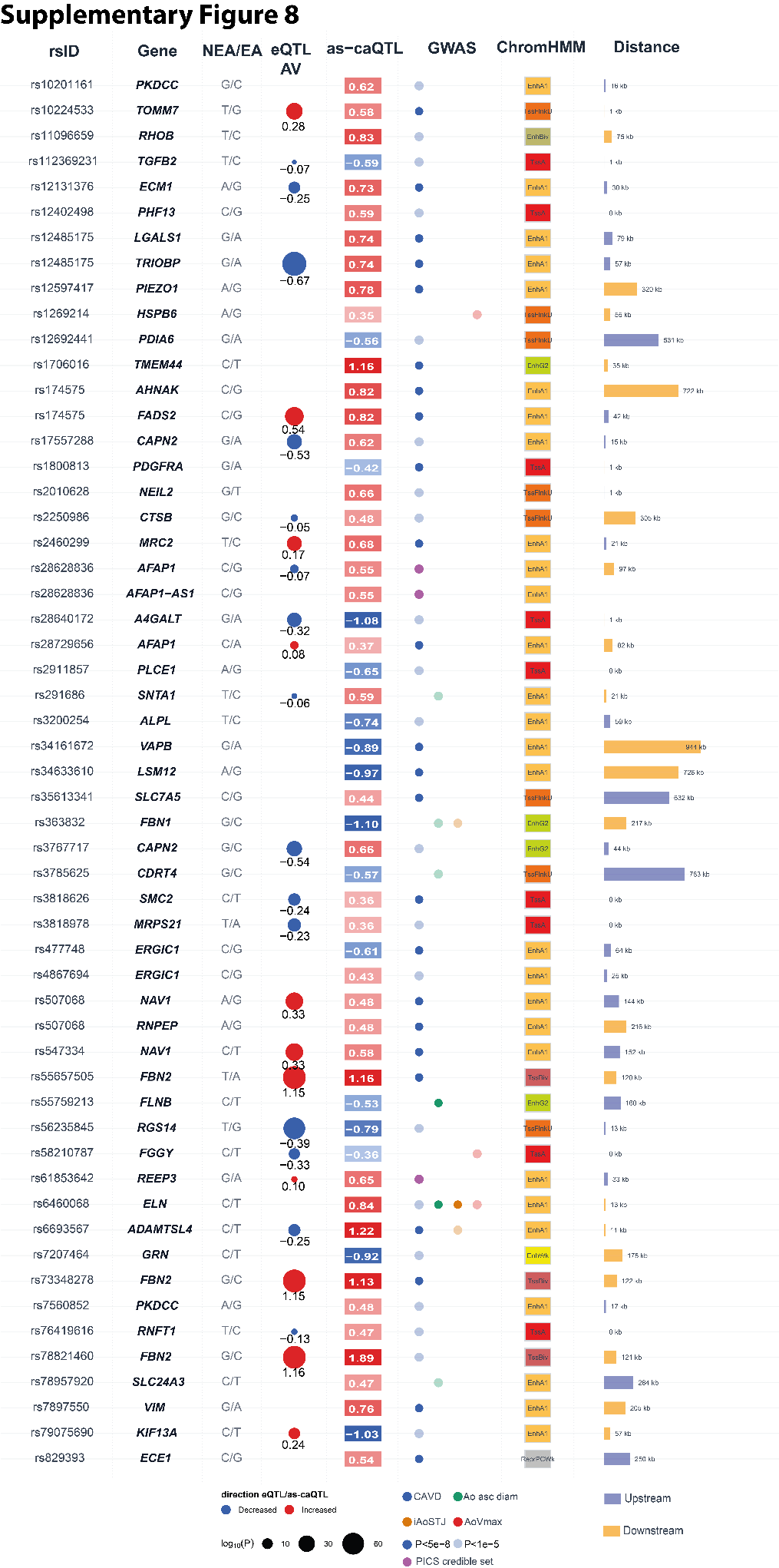

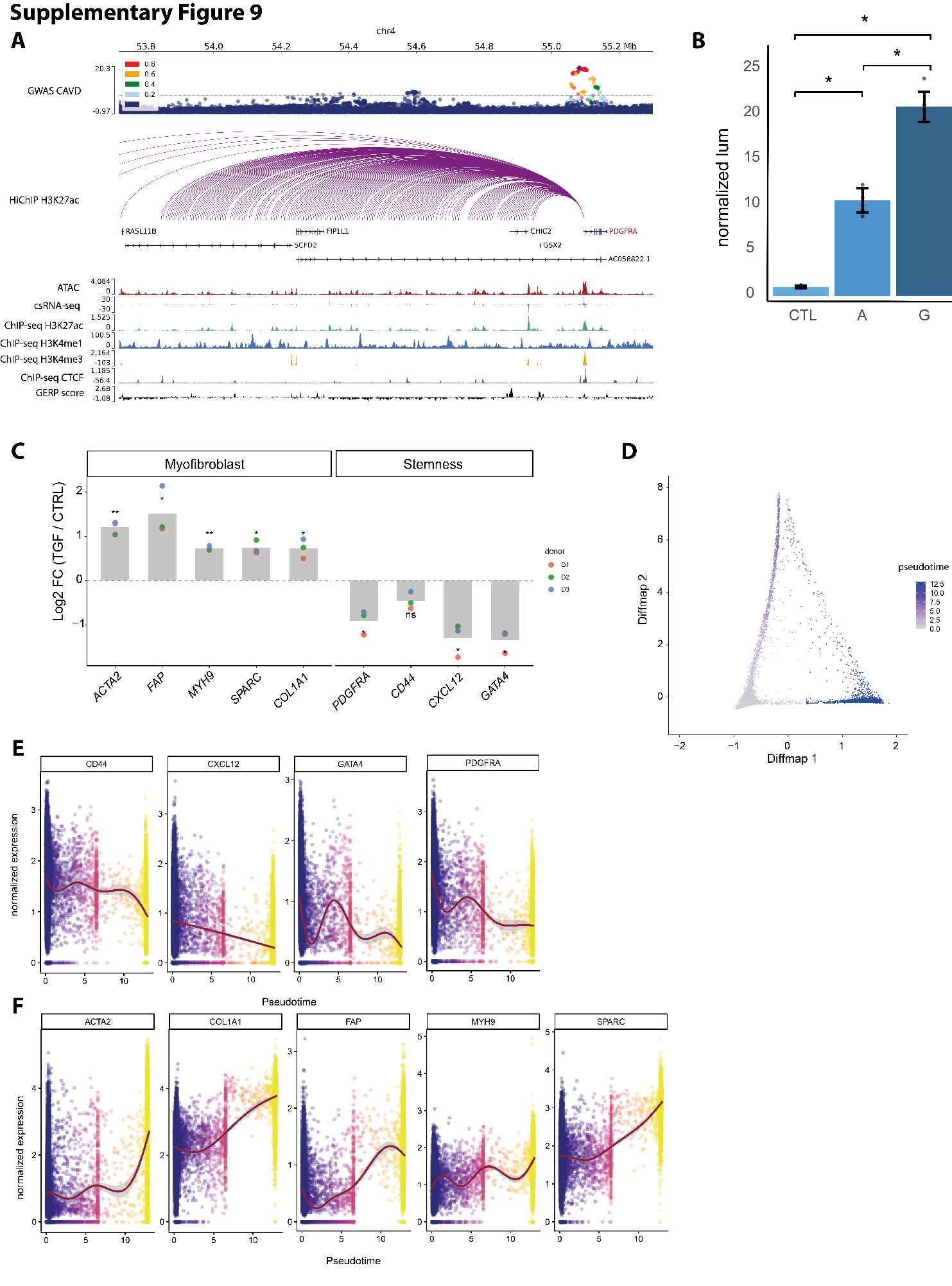

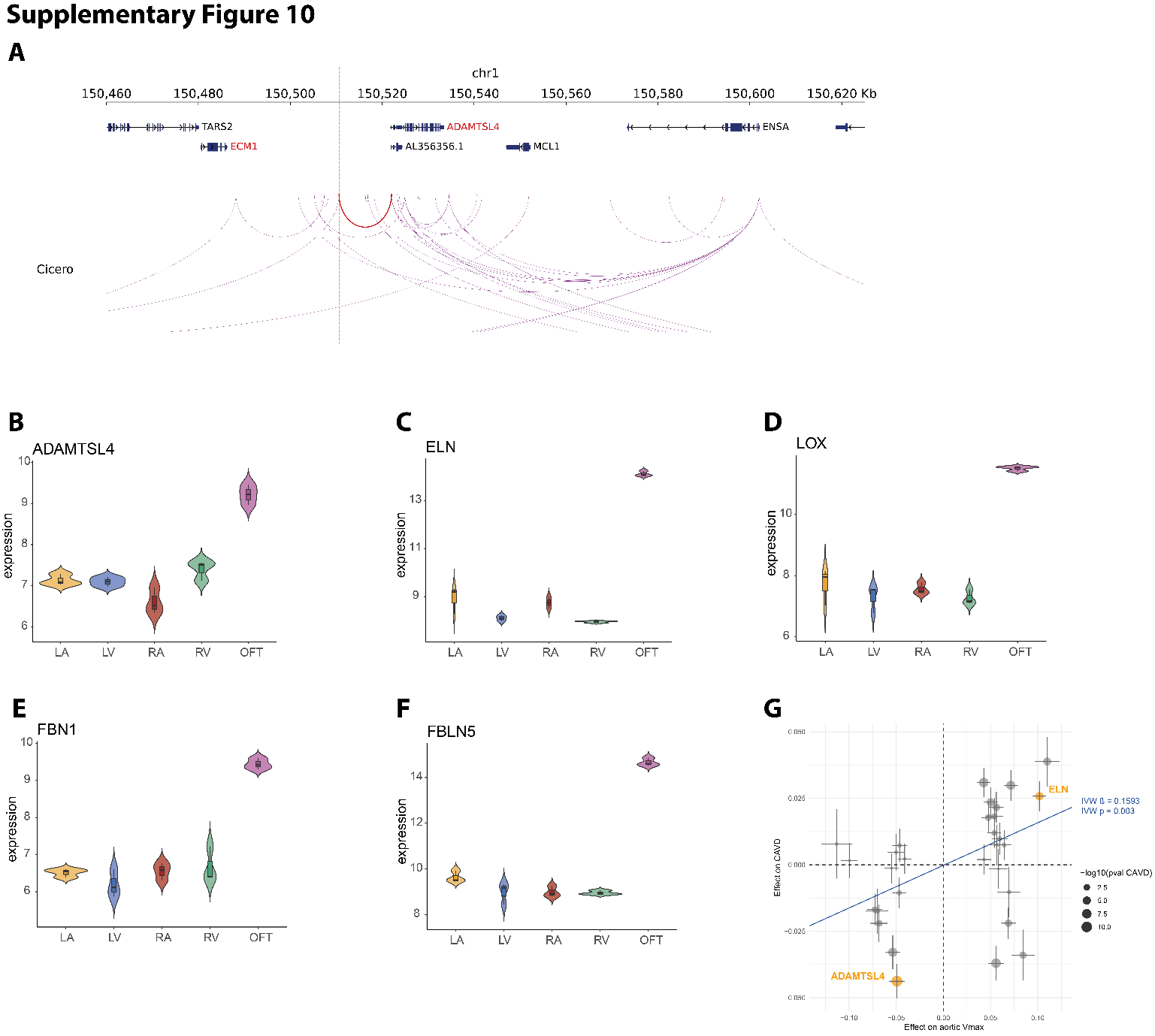

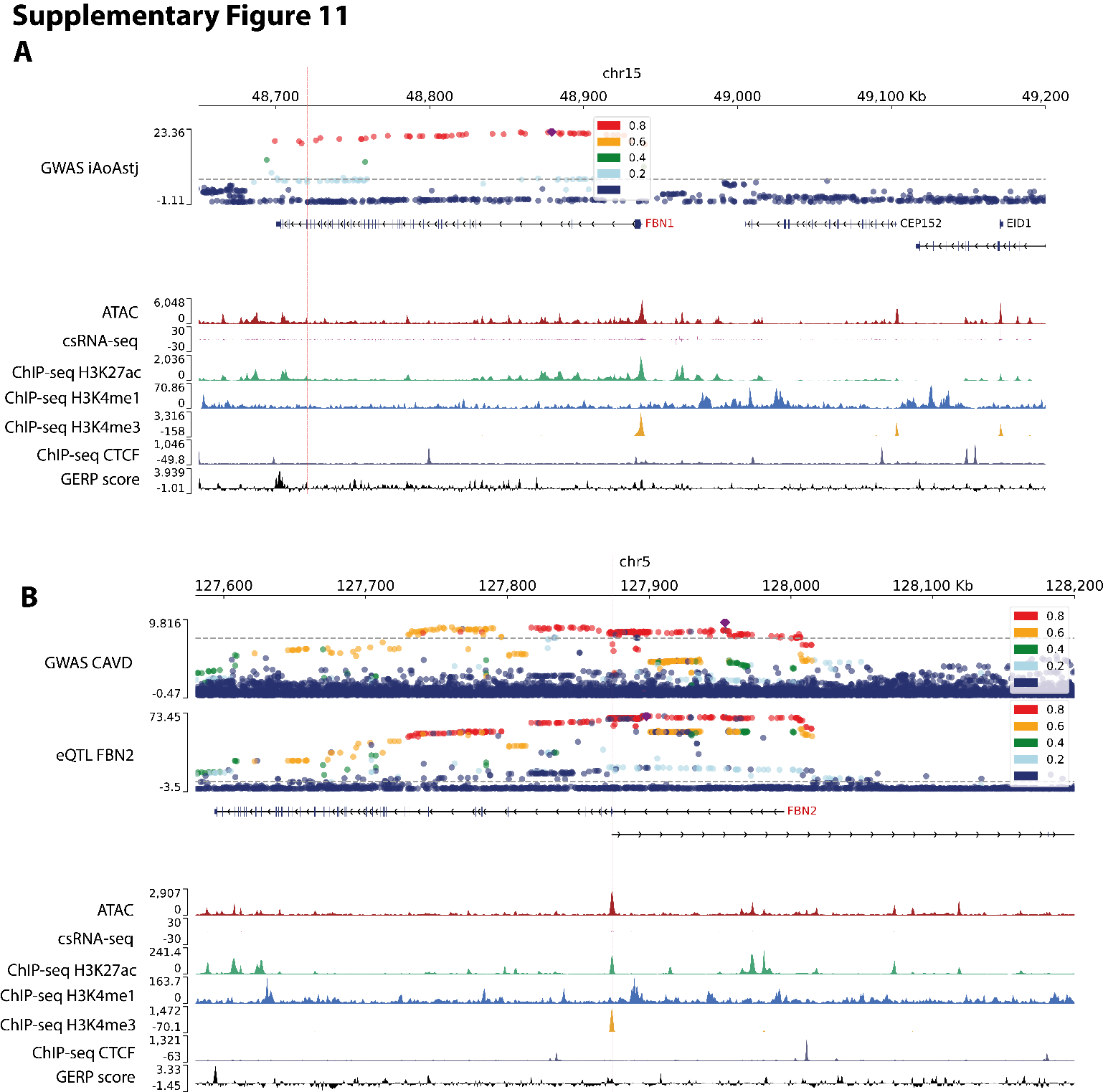

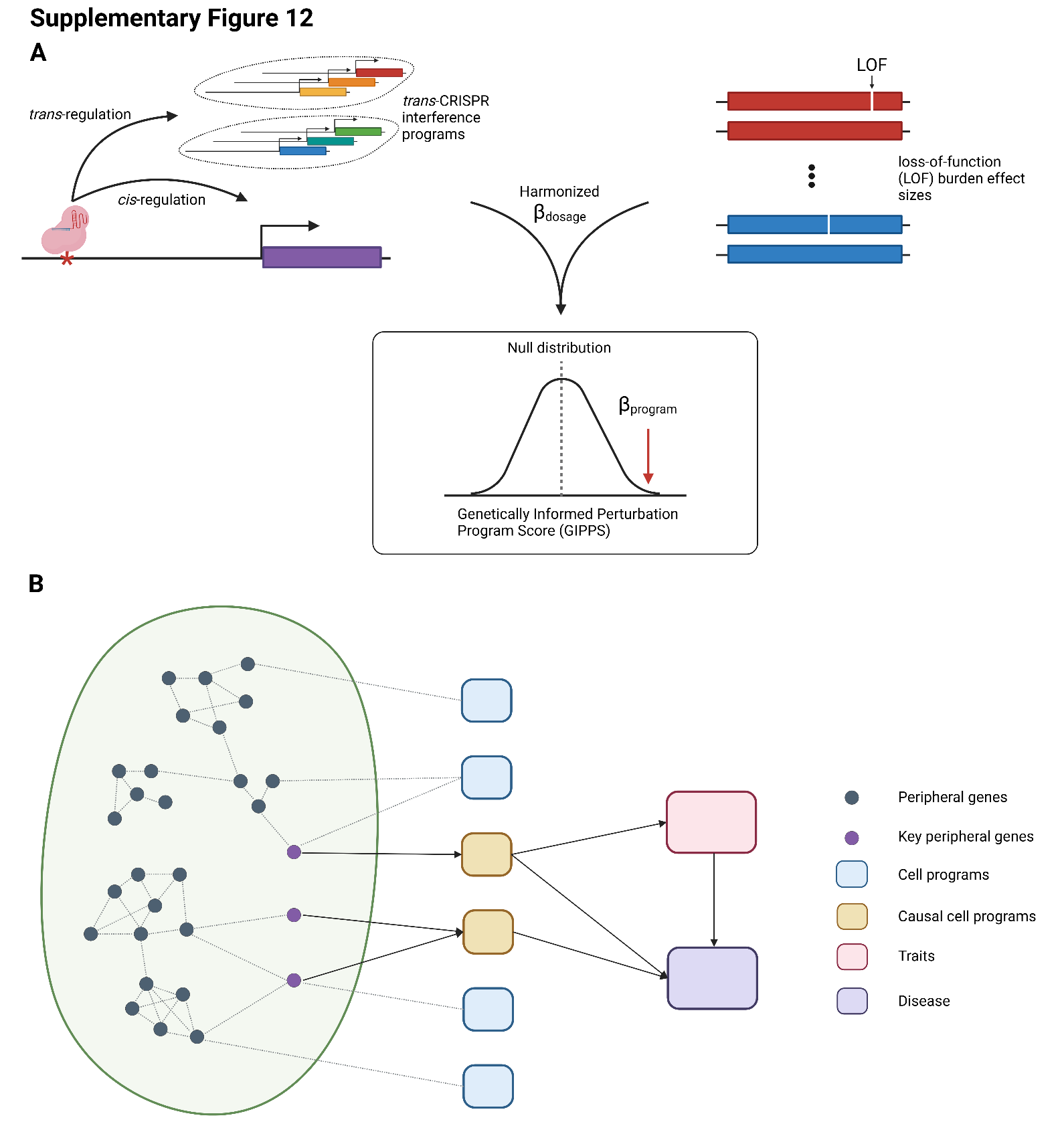

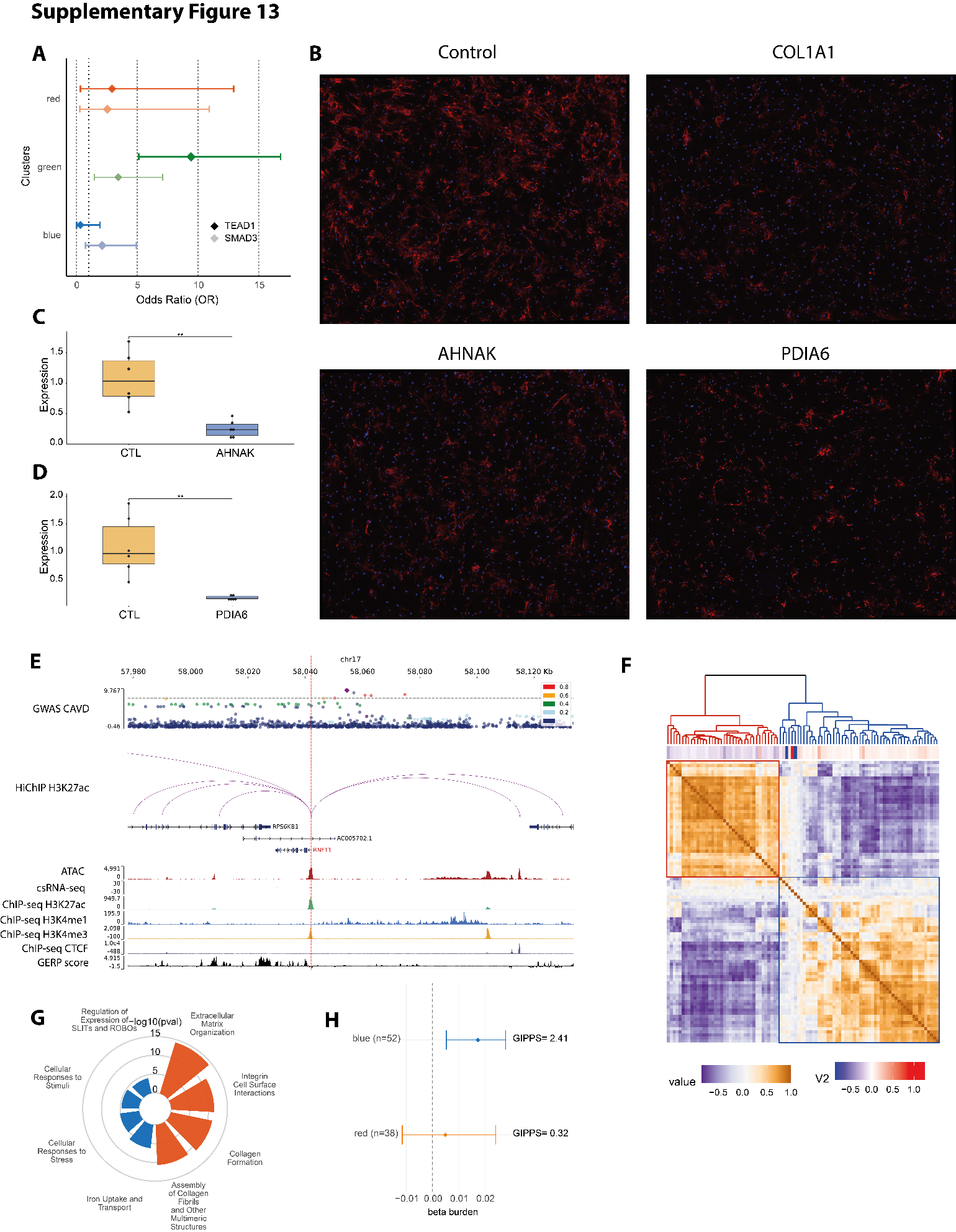

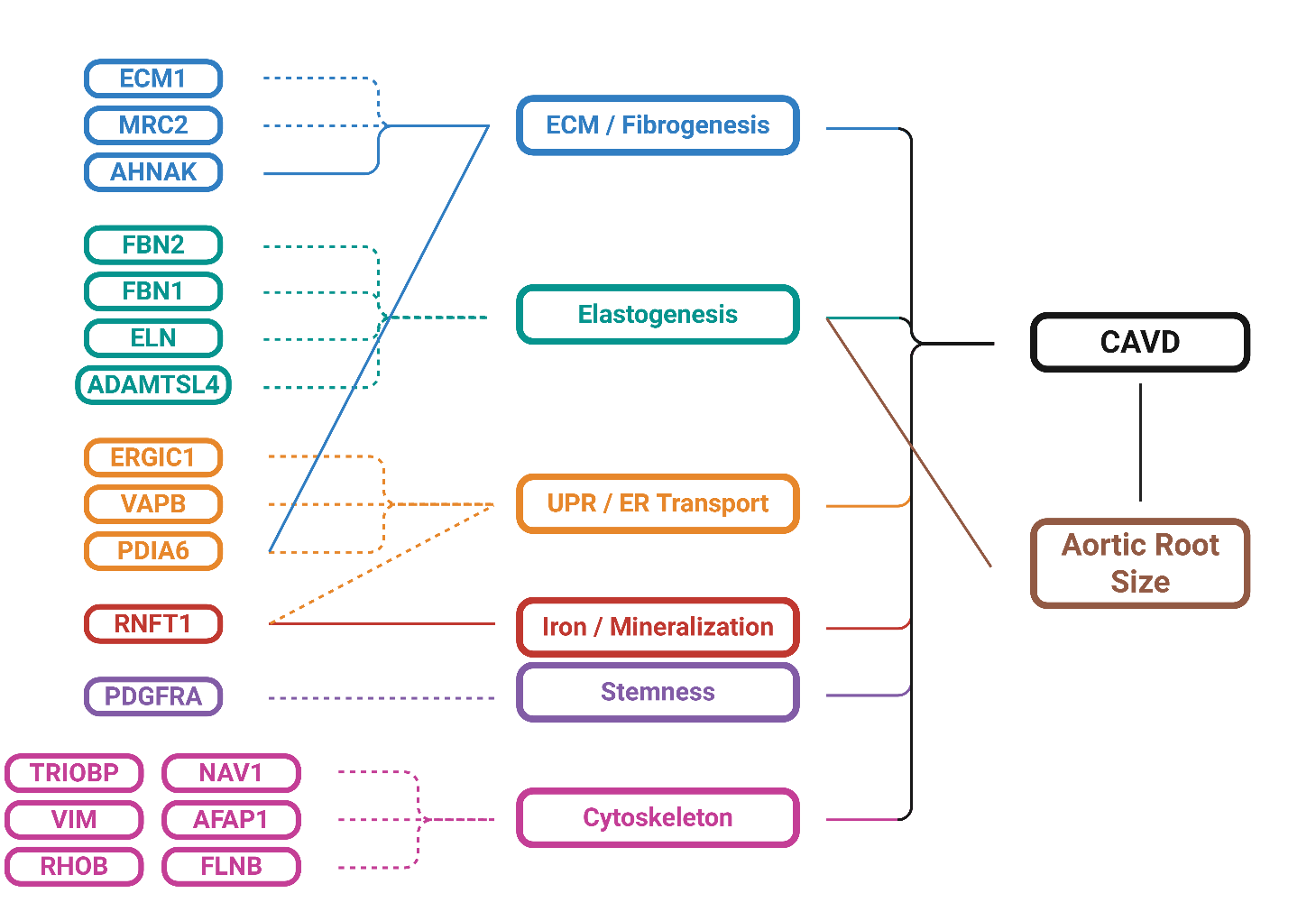
**Supplementary Figure 14**
